# Improved prime editors enable pathogenic allele correction and cancer modelling in adult mice

**DOI:** 10.1101/2020.12.15.422970

**Authors:** Pengpeng Liu, Shun-Qing Liang, Chunwei Zheng, Esther Mintzer, Yan G. Zhao, Karthikeyan Ponnienselvan, Aamir Mir, Erik J. Sontheimer, Guangping Gao, Terence R. Flotte, Scot A. Wolfe, Wen Xue

## Abstract

Prime editors (PEs) mediate genome modification without utilizing double-stranded DNA breaks or exogenous donor DNA as a template. PEs facilitate nucleotide substitutions or local insertions or deletions within the genome based on the template sequence encoded within the prime editing guide RNA (pegRNA). However, the efficacy of prime editing in adult mice has not been established. Here we report an NLS-optimized SpCas9-based prime editor that improves genome editing efficiency in both fluorescent reporter cells and at endogenous loci in cultured cell lines. Using this genome modification system, we could also seed tumor formation through somatic cell editing in the adult mouse. Finally, we successfully utilize dual adeno-associated virus (AAVs) for the delivery of a split-intein prime editor and demonstrate that this system enables the correction of a pathogenic mutation in the mouse liver. Our findings further establish the broad potential of this genome editing technology for the directed installation of sequence modifications *in vivo*, with important implications for disease modeling and correction.

## Introduction

Disease-associated genetic variations, including deletions, insertions and base substitutions, require precise gene correction strategies that are both robust and flexible^1^. Homology-direct repair (HDR) enables precise genome editing through an exogenous donor DNA. However, HDR is inefficient in most therapeutically relevant cell types, especially in post-mitotic cells^2-4^. Base editing enables efficient nucleotide transitions without inducing double-strand breaks (DSBs)^5^. However, targeted nucleotide transversions, deletions and insertions are not easily facilitated by well-established editing systems. In addition, depending on the local sequence context, base editing systems can also convert “bystander” nucleotides within the same editing window, which may be mutagenic, leading to the creation of unproductive or counter-productive alleles.

The prime editor (PE) is a genome editing tool that can produce template-directed local sequences changes in the genome without the requirement for a DSB or exogenous donor DNA templates ^6^. A PE comprises a fusion protein consisting of a catalytically impaired SpCas9 nickase (H840A) fused to an engineered reverse transcriptase (RT). A prime editing guide RNA (pegRNA) targets the PE to the desired genomic sequence and encodes the primer binding site (PBS) and reverse transcriptase template to enable the RT to copy the new genetic information into the target genomic locus ^6^. Prime editing enables nucleotide conversion and targeted sequence insertions and deletions based on the RT template sequence encoded within the pegRNA^6^. PE2 harbors five mutations within the M-MLV RT that improve editing efficiency^6^. PE3 uses an additional sgRNA to direct SpCas9^H840A^ to nick the non-edited DNA strand to encourage the edited strand to be utilized as a repair template by DNA repair factors, leading to further increases in editing efficiency^6^.

The ability to precisely install or correct pathogenic mutations regardless of their composition makes prime editing an intriguing approach to perform somatic genome editing in model organisms to study disease processes or to utilize for therapeutic applications. Previous studies have used prime editing to recode loci in cultured cells^6^, plants^7^, stem cells^8,9^, and mouse zygotes^10,11^. However, PE delivery in adult animals has not yet been described. In this study, we report a nuclear localization signal sequence optimized PE2 (referred to as PE2*) with higher editing efficiency than PE2. We demonstrate that PE2* enables somatic genome editing in the liver of adult mice, where it can correct a pathogenic disease allele or introduce a directed mutation to drive tumor formation to facilitate cancer modeling. The size of a prime editor precludes its packaging in a single AAV vector. We show that dual AAV-mediated delivery of a split-intein prime editor is functional *in vivo* for gene editing in the mouse liver. These data demonstrate the feasibility of employing PE *in vivo* for the targeted, precise alteration of genomic sequence with potential utility both in model organisms and as a therapeutic modality.

## Results

### Optimized nuclear localization signal sequence composition improves prime editing

PEs have the remarkable ability to introduce a variety of different types of sequence alterations into the genome. Their editing efficiency is influenced by a variety of different parameters (PBS length, position of the RT initiation site relative to the desired sequence alteration, composition of the desired sequence alteration, relative position of the alternate strand nick, etc.). Even under optimal conditions the incorporation rate of the desired edit into the genome is incomplete^6^. Previously, we and others have noted that the composition and number of nuclear localization signal (NLS) sequences within a Cas9 effector can influence its efficiency of genome editing^3,12-16^. The original PE2 contains two bipartite SV40 (BP-SV40) NLS sequences^6^. In transient transfection assays of original PE2, we observed incomplete variable nuclear localization based on immunofluorescence: ∼60% of the protein is present in the nucleus in U2OS cells and ∼85% is present in the nucleus in HeLa cells; (**Supplementary Fig. 1**). We found that addition of a N-terminal c-Myc NLS^17^ and inclusion of both a variant bipartite SV40 NLS (vBP-SV40)^18^ and SV40 NLS at the C terminus of PE give rise to nearly complete nuclear localization of the prime editor (PE2*) (**Fig. 1a, Supplementary Fig. 1c,d**).

**Fig. 1.**
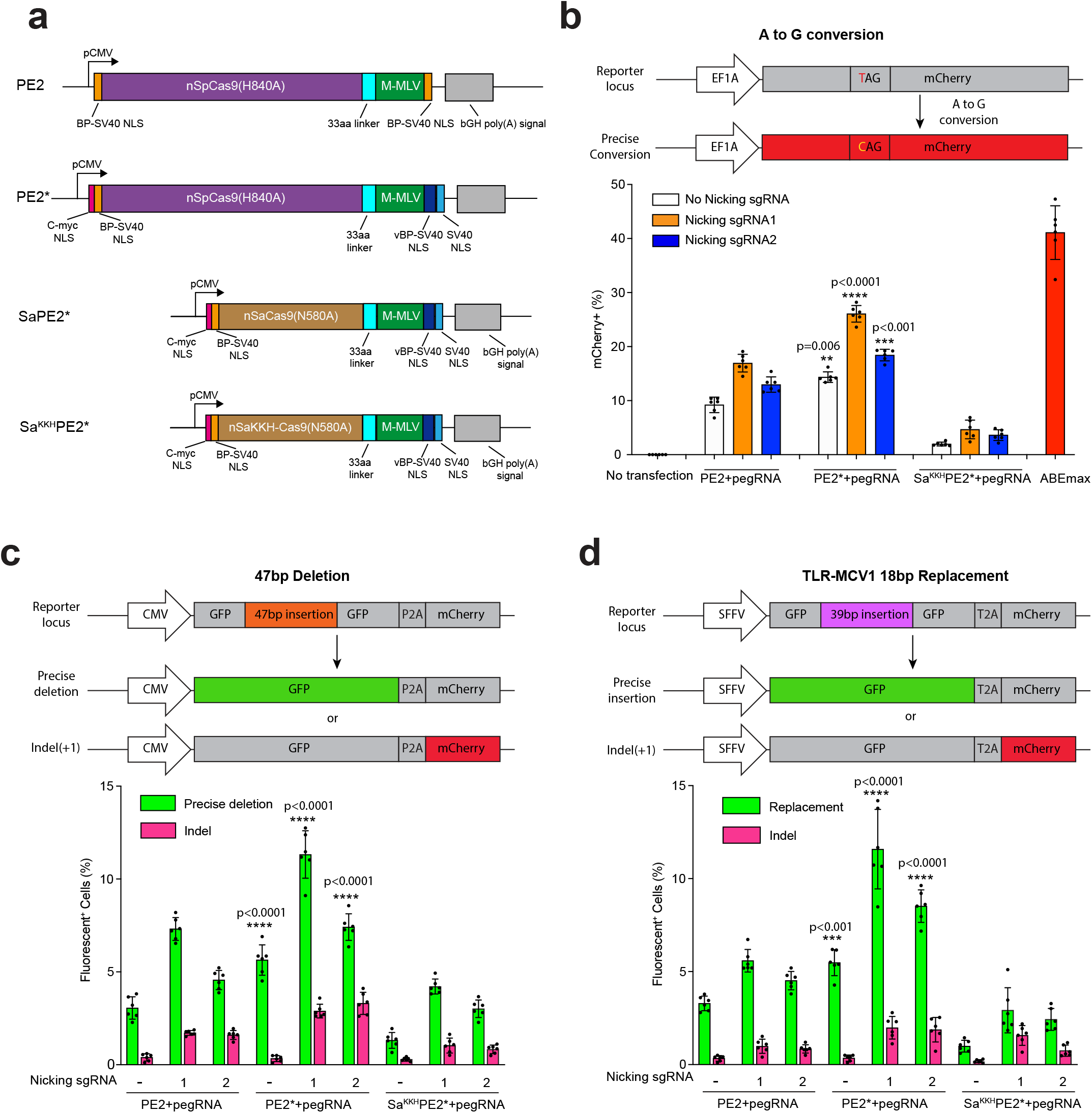
Improved NLS composition enhances prime editing efficiency. (a) Schematic representation of the original primer editor (PE2) and optimized prime editor (PE2*) carrying additional NLS sequences at the N-terminus and C-terminus. The M-MLV reverse transcriptase in PE2 was fused to a SaCas9 nickase (SaPE2*) or a SaCas9^KKH^ nickase (Sa^KKH^PE2*) with the same NLS composition to develop orthogonal prime editors. BP-SV40 NLS = bipartite SV40 NLS; vBP-SV40 NLS = variant BP-SV40 NLS. (b) Diagram of the A•T-to-G•C transition required to convert a stop codon to GLN to restore function to an mCherry reporter in HEK293T cells (top). Frequencies of targeted A•T-to-G•C transition by different prime editors (PE2, PE2* and Sa^KKH^PE2*) were quantified by flow cytometry (bottom). (c) Diagram of the deletion reporter in HEK293T cells containing a broken GFP with 47-bp insertion, P2A, and out-of-frame mCherry (top). A targeted, precise deletion of 47bp will restore GFP expression, whereas indels that create a particular reading frame alteration produce mCherry expression. Frequencies of precise deletion (GFP+) and indel (mCherry+) introduced by different prime editors (PE2, PE2*, and Sa^KKH^PE2*) were quantified by flow cytometry (bottom). (d) Diagram of the insertion reporter in HEK293 cells containing a broken GFP with 39-bp insertion, T2A and mCherry (top). A targeted, precise insertion of 18bp that substitutes for a disrupting sequence can restore GFP expression, whereas indels that create a particular reading frame alteration produce mCherry expression. Frequencies of targeted 18-bp replacement and indel generation by different prime editors (PE2, PE2*, and Sa^KKH^PE2*) were quantified by flow cytometry (bottom). All expression vectors were delivered by transient transfection. The presence of sgRNAs to promote nicking of the complementary strand is indicated in each figure legend. Results were obtained from six independent experiments and presented as mean ± SD. *\*\*P*<0.01, \**\*\*P*<0.001, \**\*\*\*P*<0.0001 by one-way ANOVA with Tukey’s multiple comparisons test between each PE2 and PE2* using the same nicking sgRNA. Source data are provided as a Source Data file.

In parallel, we constructed PE2* frameworks using the orthogonal Staphylococcus aureus (SaCas9) nickase (N580A)^19^ to replace the SpCas9 nickase (H840A). To expand the potential targeting range of these alternate prime editors systems, we utilized both the standard SaCas9 backbone that recognizes an NNGRRT PAM (SaPE2*)^19^ and the SaCas9^KKH^ variant that broadens the targeting to an NNNRRT PAM (Sa^KKH^PE2*)^20^ (**Fig. 1a**). By immunofluorescence staining using the incorporated 3xHA-tag we observed that SaPE2* or Sa^KKH^PE2* proteins were localized to the nucleus (**Supplementary Fig. 1a,b**).

To determine whether the observed improvements in nuclear localization translate into increases in editing efficiency, we evaluated the rate of nucleotide conversion for PE2 and PE2* in HEK293T mCherry reporter cells that contains a premature TAG stop codon that prevents translation of a functional protein (**Supplementary Fig. 2a,d**). PE2 and PE2* were programmed with a pegRNA designed to revert the TAG codon to CAG and delivered without and with different nicking sgRNAs. 3 days after transfection, we performed flow cytometry to quantify prime-editing efficiency. PE2* produced a 1.5-1.6 fold increase in editing efficiency (14.3% to 26.4%) relative to PE2 (9.2% to 16.5%; **Fig. 1b**). A compatible PAM is also present for Sa^KKH^PE2* within the reporter to evaluate nucleotide conversion rates, with which we observed more modest editing efficiencies (1.8% to 4.7%). All PE systems displayed lower editing activity than an adenine base editor system (ABEmax)^13^ for restoration of reporter function (**Fig. 1b**).

Next, we tested the efficiency for generating a targeted deletion using PE2, PE2* and Sa^KKH^PE2* in an HEK293T reporter line that can quantify both precise deletions and indel formation. This reporter design shares similarities to the traffic light reporter system ^21,22^. A precise deletion of 47bp will remove a sequence insertion disrupting GFP expression, whereas indels that produce a particular reading frame alteration restore mCherry expression (**Supplementary Fig. 2b,d**). PE2* produced a 1.6-1.9 fold increase in the level of precise deletions (5.6%-11.3%) compared to PE2 (3.0%-7.3%; **Fig. 1c**). The relative level of undesired indel formation was roughly proportional to the overall activity levels for PE2 and PE2*. We also observed that Sa^KKH^PE2* could generate precise 47bp deletion with efficiencies ranging from 1.3% to 4.2% (**Fig. 1c**).

Finally, we tested the efficiency for generating a targeted insertion using PE2, PE2* and Sa^KKH^PE2* in a different HEK293T reporter line (TLR-MCV1) that can quantify both precise insertions and indel formation^23^. A targeted, precise replacement of 39bp disruption sequence with a 18bp missing sequence element can restore GFP expression, whereas indels that produce a different reading frame alteration restore mCherry expression (**Supplementary Fig. 2c,d**). PE2* led to a 1.7 to 2.1-fold increase in the level of precise insertions (5.5%-11.6%) compared to PE2 (3.2%-5.5%; **Fig. 1d**). Again, the relative level of undesired indel formation was roughly proportional to the overall activity levels for PE2 and PE2*. We also observed that Sa^KKH^PE2* could generate 18bp replacement with efficiencies ranging from 1.3% to 4.2% (**Fig. 1d**). Across all of these reporter systems, we observed that nicking the nonedited strand (PE3 format) increased the editing efficiency by 1.5-to 2.4-fold and the indel rate by 0.2 % to 3.3 % compared to pegRNA only in both PE2 and PE2* (**Fig. 1b-d**), consistent with previous observations^6^. Together, these results demonstrate that PE2* performed nucleotide conversion, sequence deletion or insertion more efficiently than PE2. In addition, the new SaCas9-based PEs displayed appreciable genome editing activity.

### Improved prime editor increases editing efficiency at endogenous loci

Next, we compared the editing efficiency of PE2 and PE2* for the creation of previously described nucleotide substitutions, deletions and insertions at the *EMX1* locus^6^. HEK293T cells were transfected with different prime editors, pegRNAs and different nicking sgRNAs. Genomic DNA was isolated and editing outcomes at each target site were quantified by high-throughput sequence (HTS). PE2* (3.1%-6.5%) led to an average 1.9-fold increase in the rate of point mutation introduction compared to PE2 (1.5%-3.7%) (**Fig. 2a**). Targeted 3bp deletions were generated at 1.4 to 2.1-fold higher rate by PE2* (2.1%-6.0%) than PE2 (1.5%-2.9%) (**Fig. 2a**). Targeted 6bp insertions were generated at 1.7 to 2.4-fold higher rate by PE2* (2.2%-3.1%) than PE2 (0.9%-1.8%) (**Fig. 2a**). As observed with the various reporter systems, the level of indel formation was roughly proportional to the activity levels of PE2 and PE2*. Together, these observations suggest that PE2* has broadly improved editing efficiency at endogenous loci.

**Fig. 2.**
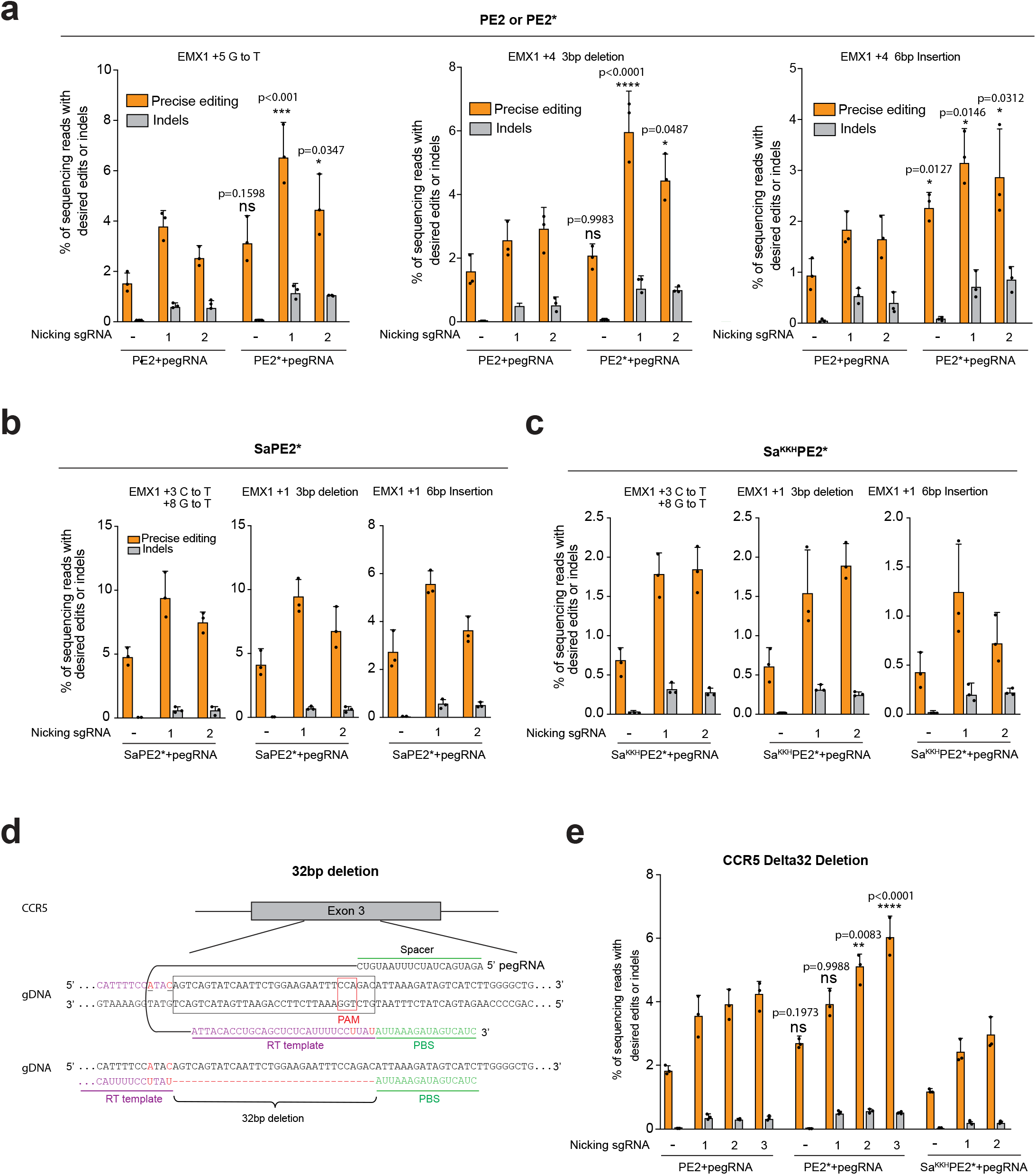
Improved PE2* increases editing efficiency at endogenous loci. (a) Comparison of editing efficiency for nucleotide substitution, targeted 3-bp deletion, and 6-bp insertion with PE2 and PE2* at *EMX1* locus in HEK293T cells. Indels broadly indicates mutations to the endogenous sequence that do not result in the desired sequence alteration. (b-c) Editing efficiency for nucleotide substitution, targeted 3-bp deletion, and 6-bp insertion with SaPE2* (b) and Sa^KKH^PE2* (c) at *EMX1* locus in HEK293T cells. (d) Sequence of CCR5 locus and pegRNA used for the 32bp deletion. Two mutations in red were included to demonstrate that sequence collapse was not a function of nuclease-induced microhomology mediated deletion and to reduce re-cutting of deletion allele. Bottom panel shows the alignment of pegRNA with the *CCR5* sense strand. (e) Comparison of efficiency for generating a targeted 32-bp deletion with PE2, PE2*, and Sa^KKH^PE2* within *CCR5* in HeLa cells. All expression vectors were delivered by transient transfection. The presence of sgRNAs to promote nicking of the complementary strand is indicated in each figure panel. Results were obtained from three independent experiments and presented as mean ± SD. *\*P*<0.05, *\*\*P*<0.01, \**\*\*P*<0.001, \**\*\*\*P*<0.0001 by one-way ANOVA with Tukey’s multiple comparisons test. ns, not significant. Source data are provided as a Source Data file.

We also compared the editing efficiency of SaPE2* and Sa^KKH^PE2* for the creation of similar nucleotide substitutions, deletions and insertions at the *EMX1* locus. Notably, SaPE2* installed point mutations at two positions in the *EMX1* locus with editing efficiency from 4.7% to 9.3% and a modest indel rate (0.0%-0.5%). Targeted 3-bp deletion and 6-bp insertion were introduced by SaPE2* with an editing efficiency of 4.1% to 9.4% and 2.7% to 5.5%, respectively. Indel induction generated by SaPE2* ranged from 0.0% to 0.6% (**Fig. 2b**).Overall, Sa^KKH^PE2* exhibited lower editing efficiency at the *EMX1* locus than SaPE2* with the same set of pegRNAs (typically between 1 and 2%; **Fig. 2c**). Notably, at these loci the editing efficiencies for SaPE2* were similar to the rates obtained with PE2*, suggesting that the SaPE2* platform has the potential to broaden the scope of available prime editing systems.

A homozygous 32-bp deletion in the *CCR5* gene is associated with resistance to human immunodeficiency virus (HIV-1) infection^24-26^. We evaluated the utility of PE2 and PE2* to generate a large, therapeutically relevant 32-bp deletion within *CCR5* that recapitulates the HIV-1 resistance allele (**Fig. 2d**). We utilized linear amplification to incorporate unique molecular identifiers (UMI) prior to sequencing^27^ to avoid PCR amplification bias for the assessment of the deletion rate in the population of treated cells. Both PE2 and PE2* were able to generate the desired 32-bp deletion within the *CCR5* locus in HeLa cells, where the PE2* editor displayed higher deletion rates than PE2 (average 1.4-fold across all conditions) with a maximum efficiency of about 6.0% (**Fig. 2e, Supplementary Fig. 3**). Sa^KKH^PE2* exhibited lower editing efficiency than PE2 for the generation of the 32-bp deletion, with a maximum efficiency of 2.9% (**Fig. 2e, Supplementary Fig. 3**). Overall, these results demonstrate that PE can introduce a therapeutically relevant deletion in the *CCR5* gene in human cells.

### Improved PE increases the correction efficiency of a pathogenic mutation *in vivo*

Alpha-1 antitrypsin deficiency (AATD) is an inherited disorder that is caused by mutations in the Serpin Peptidase Inhibitor Family A member 1 (*SERPINA1*) gene^28^. The E342K mutation (G to A) in *SERPINA1* (PiZ allele) is the most frequent mutation and causes severe lung and liver disease^28^. Patients with homozygous mutation in *SERPINA1* (PiZZ) have PiZ protein aggregates in hepatocytes and lack of functional AAT protein in the lung. The PiZ transgenic mouse contains 16 copies of the human *SERPINA1* PiZ allele and is a commonly-used mouse model of human AATD ^29^. For the correction of the E342K mutation, there are some challenges for the utilization of an adenine base editor: no optimal NGG PAM is present nearby for SpCas9, and in addition to the target adenine there are several other adenines that may also be susceptible to base conversion (“bystander effect”)^30^. Consequently, we investigated the utilization of the PE platform to correct this pathogenic mutation. To test the efficiency of different prime editors at this locus, we first evaluated the generation of the pathogenic E342K mutation (via G-to-A conversion) in wildtype *SERPINA1* in HEK293T cells. A series of different nicking sgRNAs were also evaluated in conjunction with the PEs (PE3 strategy^6^). We observed 1.6 to 3.4-fold increase for G-to-A base transition in *SERPINA1* by PE2* (6.4%-15.8%) compared to PE2 (1.9%-9.9%; **Fig. 3a**). The average rate of indel generation with a nicking sgRNA slightly increased with PE2* (0.1% −3.8%) compared to PE2 (0.0%-2.2%). Sa^KKH^PE2* exhibited lower overall editing efficiency for the installation of the E342K mutation (1.1%-4.4%). Indel generation at the target site by Sa^KKH^PE2* ranged from 0.0% to 1.4% (**Fig. 3a**).

**Fig. 3.**
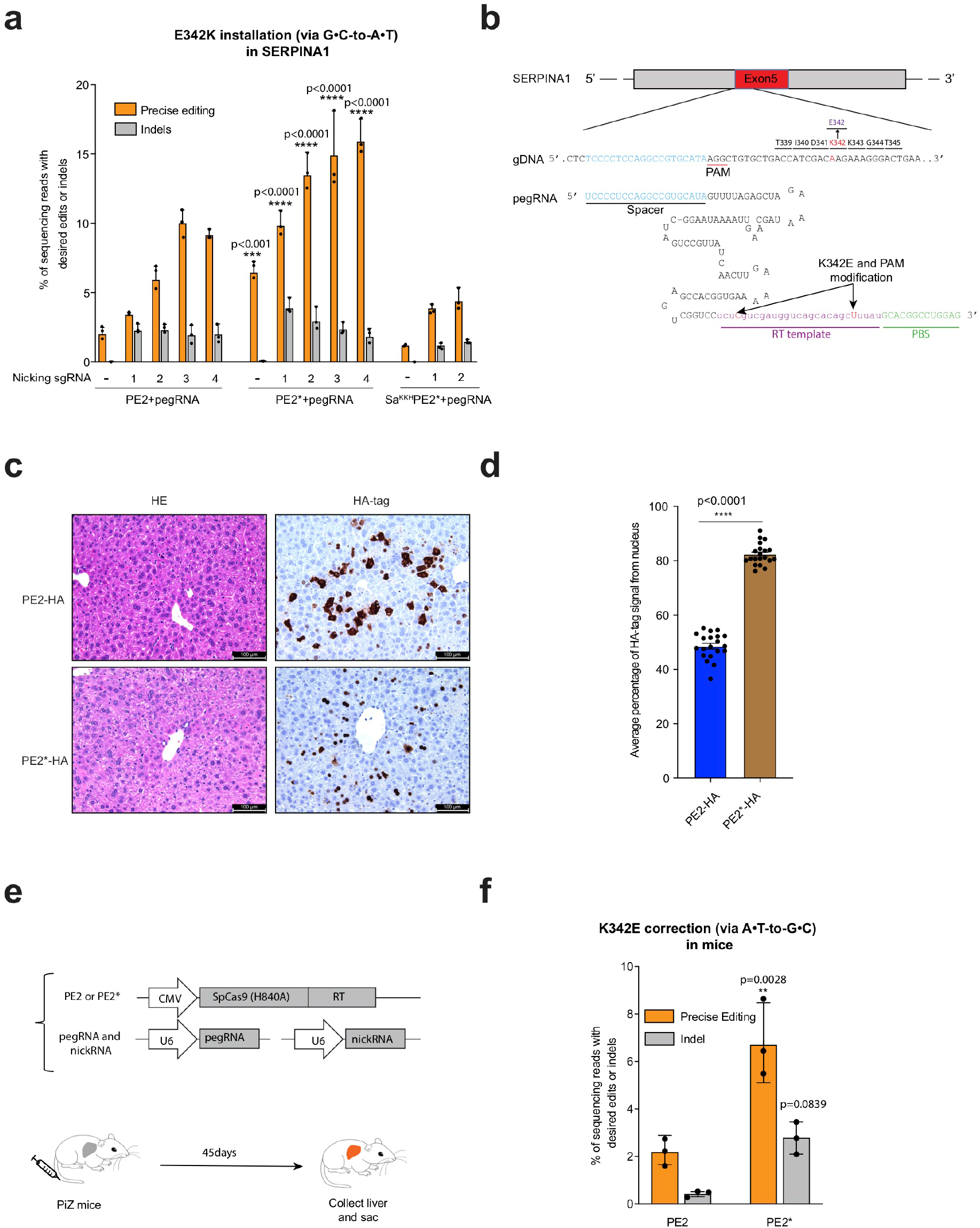
Improved PE2* increases the correction efficiency of a pathogenic mutation *in vivo*. (a) Installation (via G•C-to-A•T) of the pathogenic *SERPINA1* E342K mutation in HEK293T cells using PE2, PE2*, and Sa^KKH^PE2*. Editing efficiencies reflect sequencing reads which contain the desired edit. The presence of sgRNAs to promote nicking of the complementary strand is indicated on the x-axis. Results were obtained from three independent experiments and are presented as mean ± SD. (b) pegRNA used for correction (via A•T-to-G•C) of the E342K mutation includes a spacer sequence, a sgRNA scaffold, a RT template including edited bases (red) and a primer-binding site (PBS). A PAM mutation (AGG to AAG) was introduced to reduce re-cutting of the locus that results in a synonymous codon change. (c) Evaluating PE expression and subcellular distribution in mouse liver. FVB mice were injected with PE2 or PE2* expression plasmids containing a 3xHA-tag. IHC was performed with an HA-tag antibody. Scale bars: 100 µm (20X lens). (d) Average percentage of HA-tag signal from nucleus. Each dot is the average calculated signal intensity within the nucleus relative to the whole cell from all positive cells in a microscopic image. Numbers are mean ± sem (n = 20 total images from 3 mice). (e) Schematic overview of correction strategy of the *SERPINA1* E342K mutation in PiZ transgenic mouse model of AATD. Prime editor, pegRNA and nicking sgRNA plasmid were delivered by hydrodynamic tail-vein injection. (f) Comparison of the efficiency of K342E correction and indels in mouse livers in PE2 or PE2* treatment groups. Precise editing is defined as the fraction of sequencing reads with both A to G prime editing and synonymous PAM modification. Results were obtained from three mice and presented as mean ± SD. *\*\*P*<0.01, \**\*\*P*<0.001, \**\*\*\*P*<0.0001 by one-way ANOVA with Tukey’s multiple comparisons test. Source data are provided as a Source Data file.

Next, we investigated the ability of prime editors to directly correct a pathogenic mutation *in vivo*. Hydrodynamic injection can deliver plasmid DNA to 20-30% of hepatocytes ^31^. Using a PE2 plasmid encoding an HA-tag, we observed that 19.98 ± 0.88% hepatocytes were HA-tag positive (**Supplementary Fig. 4a**). Based on the editing results for *SERPINA1* in HEK293T cells, we chose the nicking sgRNA3 for use with the PEs for the *in vivo* experiments. We utilized a different pegRNA designed to revert the E342K mutation (**Fig. 3b)**. A PAM mutation (AGG to AAG) was also included to reduce re-cutting of the locus that introduced a synonymous codon change (**Fig. 3b)**. In the mouse liver, PE2* shows increased nuclear localization compared to PE2 (**Fig. 3c-d)**. We introduced PE2 or PE2*, pegRNA and nicking sgRNA into the liver of PiZ mice (n=3/group) through hydrodynamic tail vein injection (**Fig. 3e**). 45 days after injection, livers of PiZ mice were collected and DNA was purified for HTS analysis. Notably, we observed 3.1-fold increase for A-to-G correction in PiZ *SERPINA1* by PE2* (6.7% on average) compared to PE2 (2.1%). The indel rate at the locus was also increased from 0.4% to 2.7% (**Fig. 3f**). Interestingly, we observed a low frequency of large deletions between pegRNA and nicking sgRNA (**Supplementary Fig. 4b**). Together, these data demonstrate that prime editors can restore the wild type *SERPINA1* allele thereby enabling pathogenic gene correction in adult mice.

### Improved PE enhances tumor burden by somatic engineering in the mouse liver

*CTNNB1* (β-catenin) is a commonly mutated gene in hepatocellular carcinoma^32^. Overexpression of a mutant *Ctnnb1* and Myc oncogene have been used to generate liver cancer models^14^. To explore the potential of prime editors to drive tumor formation *in vivo*, we delivered PE2 or PE2*, a pegRNA used for installation (via C-to-T) of the oncogenic S45F mutation in *Ctnnb1* (**Fig. 4a**), and a nicking sgRNA to the livers of adult FVB mice (n=4/group) by hydrodynamic tail vein injection (**Fig. 4b**). A MYC transposon and transposase were co-injected to provide a second oncogenic driver necessary for tumor formation in conjunction with the *Ctnnb1* mutation^33^. 25 days after injection, livers of adult mice were collected and tumor nodules on the liver were quantified. PE2-treated animals showed an average 5.5 ± 1.1 tumors per mouse, whereas PE2*-treated mice displayed higher rates of tumor formation, with an average 10.0 ± 2.7 tumors on the liver (**Fig. 4c,d**). Consistent with gain of function of the S45F mutation, liver tumors were positive for nuclear β-Catenin^14^ (**Supplementary Fig.5**). Sanger sequence of gDNA from the tumor nodules showed precise conversion of S45F in *Ctnnb1* (**Fig. 4e**).

**Fig. 4.**
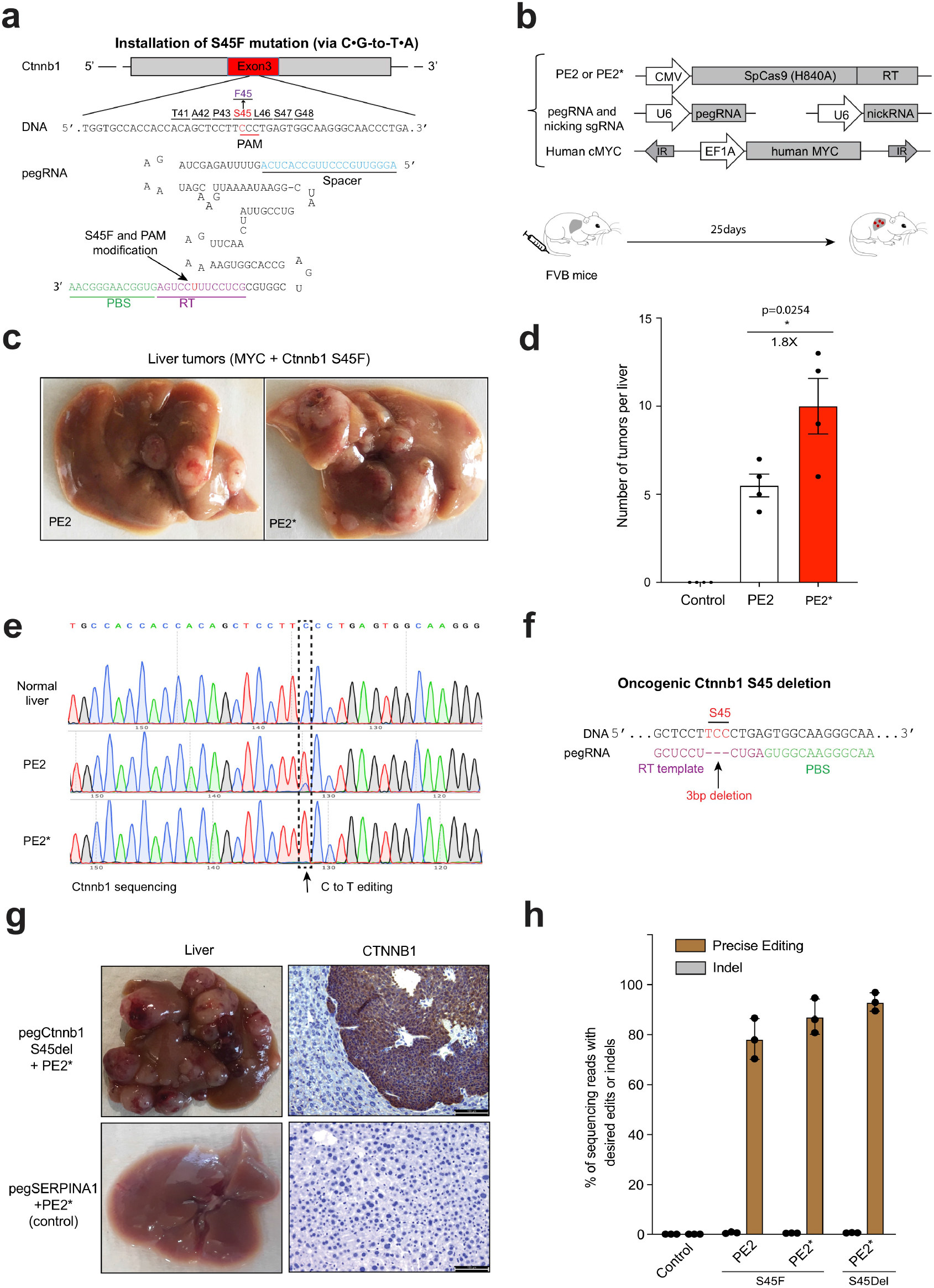
Generating mouse cancer models using improved PE2*. (a) pegRNA used for installation (via C•G-to-T•A) of the oncogenic S45F in *Ctnnb1* in mouse liver. (b) Schematic overview of the somatic cell editing strategy to drive tumor formation. Prime editor (PE2 or PE2*), pegRNA for *Ctnnb1* S45F and nicking sgRNA plasmids were delivered by hydrodynamic tail-vein injection along with the MYC transposon and transposase plasmids. (c) Representative images of tumor burden in mouse liver with PE2 or PE2*. (d) Tumor numbers in the livers of mice 25 days after injection with PE2 or PE2*. Control group was pegRNA only. Results were obtained from 4 mice and presented as mean ± SD. (e) Sanger sequencing from normal liver and representative tumors. The dashed box denotes C to T editing in tumors. *\*P*<0.05 by one-way ANOVA with Tukey’s multiple comparisons test. (f) Schematic of *Ctnnb1* S45 deletion strategy using PE2* (S45del). pegRNA used for 3bp deletion (TCC) is shown. (g) PE2* treatment leads to oncogenic activation of *Ctnnb1*. Prime editor (PE2*), pegRNA (*Ctnnb1* S45del or SERPINA1) and nicking sgRNA plasmids were delivered by hydrodynamic tail-vein injection along with the MYC transposon and transposase plasmids. Mice treated with the peg*Ctnnb1* S45del (n=4) displayed a large number of liver tumors whereas mice treated with pegSERPINA1 as a control displayed no noticeable oncogenic lesions. beta-Catenin (CTNNB1) IHC staining was performed. Scale bars: 100 µm (20X lens). (h) Prime editing efficiency and indels determined by targeted deep sequencing in control liver and representative tumors. Results were obtained from 3 tumors in each group and presented as mean ± SD. Source data are provided as a Source Data file.

Prime editors afford the opportunity to install other types of mutations within the genome. To assess the feasibility of generating deletions *in vivo*, we designed a pegRNA to delete the S45 codon in *Ctnnb1*, which is a previously described oncogenic mutation at this locus ^34^(**Fig. 4f**). The prime editor (PE2*), pegRNA for *Ctnnb1* S45 deletion and nicking sgRNA plasmids were delivered by hydrodynamic tail-vein injection along with the MYC transposon and transposase plasmids. pegRNA *Ctnnb1* S45 deletion-treated animals showed extensive tumor formation, whereas pegRNA SERPINA1-treated animals did not induce any tumor formation (**Fig. 4g**). Deep sequencing showed that more than 80% of tumor gDNA contained precise editing removing the S45 codon (**Fig. 4h**). Together, these results demonstrate that prime editors can be used for generating tumor models by somatic cell engineering *in vivo*, and that PE2* provides a platform with improved editing activity.

### Utilizing a split-intein approach for prime editor delivery

The ∼6.3-kb coding sequence of PE exceeds the ∼4.8-kb packaging size limit of AAV^35^. To deliver prime editors with AAVs, we adapted a split Cas9 dual-AAV strategy^36^ in which the original PE2 prime editor is divided into an amino-terminal (PE2-N) and carboxy-terminal (PE2-C) segments, which are then reconstituted to full length PE by a *trans*-splicing intein^37^ (**Fig. 5a**). To ensure that each prime editor segment is smaller than the AAV packaging size limit, we divided the PE within the SpCas9 amino acid before Ser 714^38^ and used the c-Myc NLS at the N-terminus and BP-SV40 NLS at the C-terminus. We generated AAV8 particles encoding the split-intein PE, a nicking sgRNA and a pegRNA (PE3 strategy^6^) to correct the E342K mutation in SERPINA1 (**Fig. 5a**). We then characterized the performance of the split-intein AAV prime editor *in vivo*. PiZ mice were treated by tail-vein injection of a low dose dual AAV8-PE (2 × 10^11^ viral genome total) (**Fig. 5b**). Livers were harvested at 2 weeks (n=2), 6 weeks (n=3) and 10 weeks (n=3) after injection. By targeted deep sequencing, we detected 0.6 ± 0.0 % precise editing at 2 weeks. The precise editing efficiency increased significantly to 2.3 ± 0.4 % at 6 weeks and 3.1 ± 0.6 % at 10 weeks (**Fig. 5c**).

**Fig. 5.**
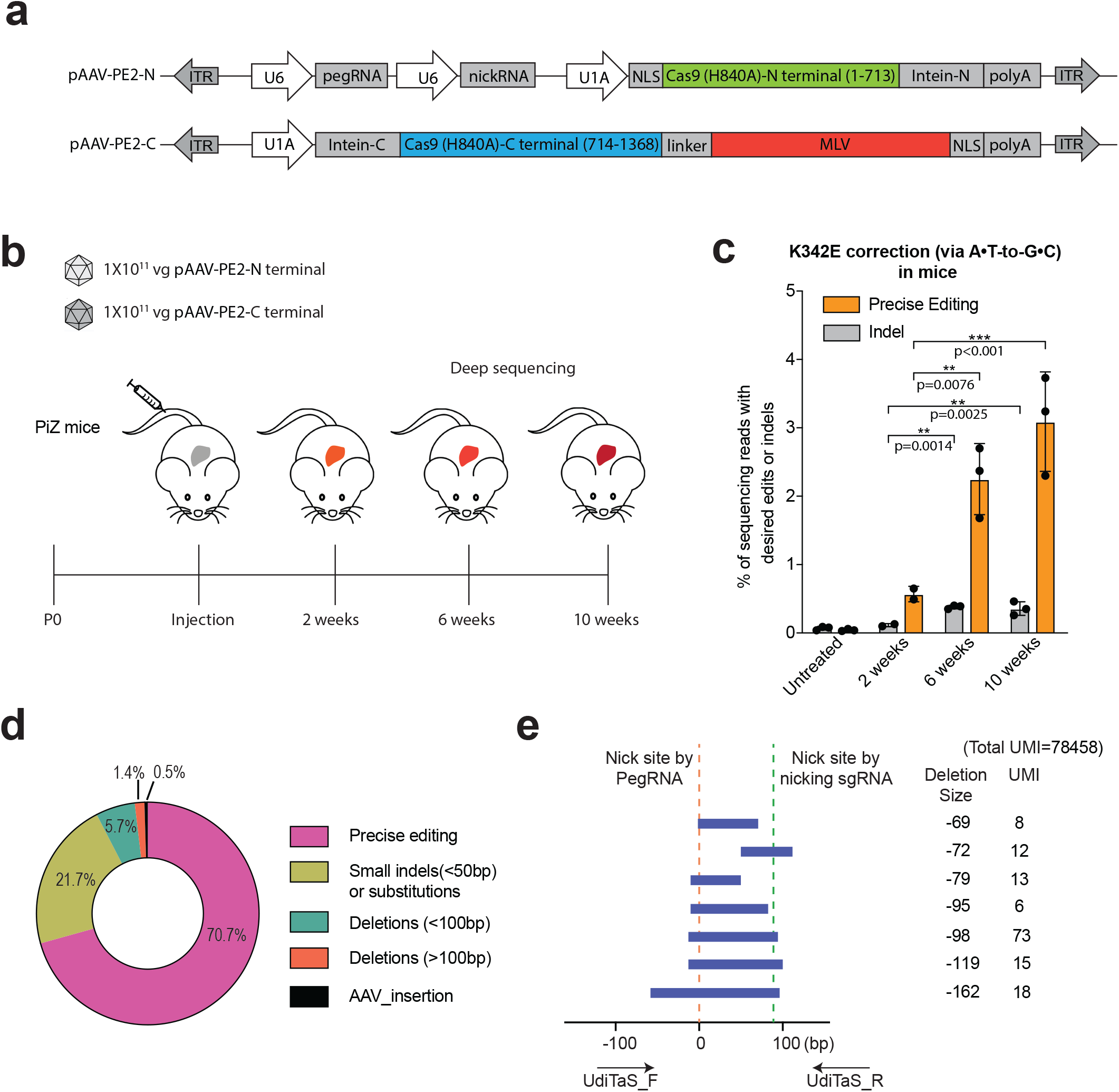
Systemic injection of dual AAV8 split-intein prime editor achieves pathogenic mutation correction in PiZ mice. (a) Schematic of split-intein dual AAV prime editor. Full-length primer editor (original PE2^6^) was reconstituted from two PE2 fragments employing the Npu DNAE split intein^37^. C, carboxy terminal; N, amino terminal. (b) Schematic of the *in vivo* dual AAV8 prime editor injection experiments. Dual AAV8 split-intein prime editor (2 × 10^11^ vg total) was delivered to six-week-old PiZ mice by tail-vein injection. Livers were harvested at 2 (n=2), 4 (n=3) and 10 (n=3) weeks after injection and the genomic DNA was isolated for sequencing. (c) Prime editing efficiency of K342E correction and indels determined by targeted deep sequencing in mouse livers of dual AAV-treated mice. Precise editing is defined as the fraction of sequencing reads with both A to G prime editing and synonymous PAM modification. Results were obtained from two (2 weeks) or three mice (6 and 10 weeks) and presented as mean ± SD. *\*\*P*<0.01, \**\*\*P*<0.001 by one-way ANOVA with Tukey’s multiple comparisons test. (d) Composition of edited alleles at *SERPINA1* by UDiTaS analysis. Circle plot shows the fraction of edits that are precise (intended base conversion), small indels (<50bp) or substitution, deletions between pegRNA and nicking sgRNA sites (<100bp), large deletions (>100bp), and AAV fragment insertion. Numbers are average of 3 mice in 10 week treated cohort. (e) The statistically significant large deletion sequences detected by UDiTaS in the 10 week treated cohort are displayed as bars spanning the sequence that is deleted (a representative liver of n=3 mice). Positions of the pegRNA and nicking sgRNA are indicated by dotted lines and the approximate positions of the locus-specific UDiTaS primers are indicated by arrows below the bar chart. The deletion size and number of UMIs associated with each deletion are indicated to the right of each bar. Statistical significance was calculated as a Benjamini-Hochberg adjusted p-value with a cut-off of 0.05. Source data are provided as a Source Data file.

We observed corresponding increases of indel rates at the target site by split-intein AAVs from 0.1 ± 0.0 % (2 weeks) to 0.4 ± 0.1 % (10 weeks). Utilization of the UDiTaS unidirectional sequencing approach^39^ with locus specific primers for library construction affords the opportunity to assess the rate of large deletions or other types of genomic rearrangements in an unbiased manner. UDiTaS analysis at the SERPINA1 transgene locus revealed primarily precise editing among the modified alleles. As observed by the amplicon deep sequencing, a fraction of the modified alleles contained indels and there were a small number of larger deletions between the two nicking sites or extending beyond these sites, although many of the largest deletions (>100 bp) did not meet the level of statistical significance (**Fig. 5d-e, Supplementary Fig. 6 and Supplementary table 5**). There was also evidence of a very low rate of AAV insertion at the target site (0.014% of total UMIs). Together, these results demonstrate that delivery of split-intein PE by the dual AAV8 enables low rates of precise editing via the PE3 prime editing strategy *in vivo*.

## Discussion

Prime editing systems potentially provide a powerful approach for the template-directed incorporation of a variety of types of alterations (nucleotide changes, insertions, deletions) into genomic DNA sequence without relying on homology directed repair ^6^. In principle, this provides a strategy for the correction of a variety of different disorders, since prime editing should not be dependent on the cell cycle for efficacy as is HDR^2^. Moreover, unlike HDR and MMEJ^40^ approaches for precise sequence insertions, prime editing does not require co-delivery of donor DNA. However, the length of the sequence that can be inserted is limited by the length of the encoded pegRNA. While there are many potential advantages to prime editing, the development of prime editing systems is in its initial stages, and many questions with regard to the utility of this system for genome editing remain to be addressed. This study provides an important demonstration of the *in vivo* utility of prime editors for the template-based modification of genomic sequence with implications for improving the utility of disease model systems and for the eventual translation of this tool to the correction of pathogenic disorders.

The efficacy of genome editing systems is dependent on a number of factors, one of which is the efficiency of nuclear import. In rapidly proliferating cells, the nuclear envelope provides only a modest barrier to entry for genome editing tools. However, in post-mitotic or quiescent cells the nuclear envelope may provide a greater barrier to the entry of Cas9-based systems, such that the number and composition of the NLS sequences can impact editing efficacy ^3,12-16^. By incorporating additional NLSs of varying sequence composition, we developed an improved PE (PE2*) that increases the efficiency of genome editing across multiple endogenous sites relative to the original PE2.Importantly, the observed improvements in genome editing for PE2* *in vivo* correlated with differences in the efficiency of nuclear import based on immunostaining in liver sections. Thus, demonstrating that NLS sequence composition and architecture is an important parameter to consider in the design of prime editing systems to maximize *in vivo* efficacy, as has been observed for other genome editing systems ^3,36^.

This study provides an important proof-of-concept demonstration of the feasibility of *in vivo* editing of somatic cells in mammalian systems by prime editing. We demonstrated the utility of prime editing systems for two different types of applications: the correction of a pathogenic mutation (AATD) and generation of animal cancer model. The ability to precisely correct a pathogenic mutation or install somatic mutations *in vivo* has important implications for both gene therapy and the development of mouse models to study cancer and other disorders^41^. Using hydrodynamic injection, PE2* shows higher editing efficiency (∼6%, **Fig. 3**) than previously published HDR (∼0.5%)^42^ in the mouse liver for installing oncogenic point mutations. Future work will address whether PEs can simultaneously generate cooperating oncogenic point mutations (a list of potential mutations that can be studied is in **Supplementary Table 4**).

Compared with HDR and base editors, prime editing provides a complementary method for generating precise genome alterations. Although a PE is less efficient than base editors for introducing base transitions, a PE can generate precise nucleotide substitutions in target sites where the neighboring sequence composition could potentially be challenging for precise conversion by base editing systems ^6^. Both of our *in vivo* target sites have neighboring nucleotides that would potentially be susceptible to conversion when targeted by a base editor (bystander effects). In the case of the *SERPINA1* locus, these bystander changes would introduce missense mutations that could have undesirable effects.

Prime editing systems can also produce undesired editing outcomes in some instances^11^. The use of the PE2 system in cell culture systems produces primarily precise edits^6^. The rate of precise edits can be increased through the use of an additional nicking sgRNA (PE3 strategy), but this also results in the production of a low rate of indels within the genome^6,11^. Prime editing in plant protoplasts also produces a fraction of undesired editing outcomes when employing either the PE2 and PE3 strategy^7^. Interestingly, in mouse zygotes the PE3 strategy produces alleles containing the desired edit, but a large fraction also harbor deletions of various sizes between the target site and the nicking site ^11^. Reassuringly, for our *in vivo* PE3 editing the majority of modified alleles contain the intended product without additional modifications. Similar to the observation in cell culture experiments, we observe a small fraction of the alleles that contain unintended changes (indels). A small proportion of these (∼6.6% modified alleles) contain deletions between the pegRNA and nicking sgRNA sites. Employing a sequential nicking strategy (PE3b) approach instead of PE3 may reduce the indel rates *in vivo*, as observed in other systems^6,7^, when an overlapping nicking RNA can be designed at the prime editor target site.

We established a dual-AAV system to deliver split-intein prime editors that produces precise editing *in vivo* following a single injection. While we focused on liver editing, the dual AAV-mediated prime editor developed in this study should be applicable to other organ systems. Because AATD requires a high level of gene correction to generate sufficient wildtype protein for disease amelioration^28^, our initial AAV PE design is suboptimal for AATD treatment. Future work will be required to increase PE efficiency *in vivo*. The modest editing efficiency of our AAV system may be due in part to the low vector dose (2×10^11^ vg/kg total). In addition, some aspects of our design of split-PE AAV vectors could be improved. We used c-Myc and BP-SV40 NLSs in our split-intein PE design due to their more compact size for vector packaging – further optimization of the NLSs may improve *in vivo* activity of the split-PE AAV system. Optimizing the position of intein insertion within the PE may also improve activity, as may alteration of the AAV vector layout and promoters that are utilized ^36^. The more compact SaPE2* or Sa^KKH^PE2* systems described herein may also have utility in the context of AAV delivery once they are further optimized. In addition, the development of nanoparticle-mediated delivery or direct RNP delivery could also provide alternate avenues for translation of this tool for *in vivo* therapeutic application without the constraints of AAV cargo capacity. Once the efficiency of prime editing systems *in vivo* is improved, it will also be important to explore in greater depth the frequency of off-target editing before these systems are transitioned to gene therapy applications. In summary, our findings demonstrate the broad potential scope of PEs for *in vivo* genome editing, both for disease gene correction and the development of cancer models in adult animals.

## Methods

### Generation of plasmids

To generate pegRNA expression plasmids, PCR products including spacer sequences, scaffold sequences and 3’ extension sequences were amplified with indicated primers (Supplementary table 2) using Phusion master mix (ThermoFisher Scientifc) or Q5 High-Fidelity enzyme (NewEnglandBioLabs), which were subsequently cloned into a custom vector (Supplementary sequence 2, BfuAI and EcoR I digested) by the Gibson assembly method (NEB). To generate nicking sgRNA expression plasmids, annealed oligos were cloned into BfuAI-digested vector or pmd264 vector (Supplementary sequence 2). Sequences of all pegRNA and nicking sgRNAs are listed in Supplementary Table 1. PE2* was generated through Gibson assembly, by combining SpyCas9(H840A) and the M-MLV RT from PE2 ^6^ with additional NLS sequences by PCR and inserting into a NotI/PmeI-digested pCMV-PE2 backbone (Supplementary sequence 3). SaPE2* and Sa^KKH^PE2* were generated through Gibson assembly, by combining the following three DNA fragments: (i) PCR amplified M-MLV RT with additional NLS sequences from PE2, (ii) a NotI/PmeI-digested PE2 backbone, (iii) a SaCas9 N580A nickase or a Sa^KKH^-Cas9 nickase (supplementary sequence 3). All plasmids used for *in vitro* experiments were purified using Midiprep kit including endotoxin removal step (Qiagen). pCMV-PE2 was a gift from David Liu (Addgene plasmid # 132775) ^6^. AAV-PE-N was generated through Gibson assembly, by combining the following five DNA fragments: (i) gBlock pegRNA driven by U6, (ii) gBlock nicking sgRNA driven by U6, (iii) PCR amplified N-terminal PE2 (amino acid 1-713 of SpCas9 H840A), (iv) gBlock split-intein Npu N-terminal domain, (v) a KpnI/SacI-digested AAV backbone^43^. AAV-PE-C was generated through Gibson assembly, by combining the following four DNA fragments: (i) gBlock split-intein Npu C-terminal domain, (ii) PCR amplified C-terminal PE2 (amino acid 714-1368 of SpCas9 H840A) and M-MLV RT from PE2, (iii) gBlock β-globin poly(A) signal, (iv) a KpnI/NotI-digested AAV backbone. For evaluating nuclear localization, a 3XHA-tag was inserted in PE2 or PE2* before the C-terminal BP-SV40 NLS or vBP-SV40 NLS.

### AAV vector production

AAV vectors (AAV8 capsids) were packaged at the Viral Vector Core of the Horae Gene Therapy Center at the University of Massachusetts Medical School. Vector titers were determined by gel electrophoresis followed by silver staining and qPCR.

### Generation reporter cells and Cell culture conditions

HEK293T cells were purchased from ATCC, and cells were maintained in Dulbecco’s Modified Eagle’s Medium supplemented with 10% FBS. For generation of the mCherry reporter and 47bp-insertion TLR reporter cells, we created single-copy reporter cells using the Invitrogen Flp-In system. Briefly, Flp-In 293T cells were maintained in DMEM, 10% Fetal bovine serum (FBS), 1% pen-strep and 100 μg/ml Zeocin. 1×10^6^ Flp-In 293T cells were plated in a 6 well plate 24 hours before transfection. On the day of transfection, the cells were washed and fresh media without Zeocin was added. The plasmid coding for FLP recombinase and the mCherry reporter or 47bp-insertion TLR reporter plasmid were transfected into the cells at a 9:1 ratio using polyfect (QIAGEN) according to the manufacturer’s protocol (900ng mCherry reporter or 47bp-insertion TLR reporter plasmid and 100ng FLP recombinase plasmid to make 1 μg plasmid in total). 48 hours following transfection, the cells were washed and split into a 10cm dish with fresh media. 100 μg/ml of hygromycin was used to select for cells that contained an integration of the reporter plasmid. Two weeks post selection, hygromycin resistant foci were pooled and propagated for cryopreservation and further experiments. The construction an characterization of the TLR-MCV1 reporter cells was described in ^23^. All cell types were maintained at 37°C and 5% CO_2_ and were tested negative for mycoplasma.

### Cell culture transfection/electroporation and DNA preparation

For transfection-based editing experiments in HEK293T cells or HEK293T reporter cells, cells were plated 100,000 per well on a 48-well plate. 24 hours later, the cells were co-transfected with 540ng of prime editor plasmid, 270ng of pegRNA plasmid and 90ng of Nicking sgRNA plasmid. Lipofectamine 2000 (Invitrogen) was used for the transfection according to the manufacturer’s instructions. FACS analysis was performed 3 days after transfection in HEK293T reporter cells. To detect editing efficiency in endogenous genomic loci, HEK293T cells were cultured for 3 days after transfection, and genomic DNA was isolated using QIAamp DNA mini kit (QIAGEN) according to the manufacturer’s instructions. For *CCR5* PE editing, the same amount of plasmid were delivered (540ng of prime editor plasmid, 270ng of pegRNA plasmid and 90ng of Nicking sgRNA plasmid), where 2 × 10^5^ HeLa cells were treated per electroporation using the Neon® TransfectionSystem 10 L Kit (Thermo Fisher Scientific) with the recommended electroporation parameters: Pulse voltage (1350 v), Pulse width (10 ms), Pulse number (3).

### Fluorescent reporter assay

48 h post-transfection cells are trypsinized and harvested into a microcentrifuge tube. Cells are centrifuged at 500×g for 2 min, washed once with 1× PBS, recentrifuged at 500×g for 2 min and resuspended in 1× PBS for flow cytometry (Becton Dickonson FACScan). 10,000 events were counted from each sample for FACS analysis. Experiments were performed in six replicates on different days. The data were analyzed using Flowjo v10 and are reported as mean values with error bars indicating SD. An example of the gating scheme can be found in Supplementary Note 1.

### Immunofluorescence and immunohistochemistry

HeLa and U2OS cells are transfected in six-well format via Lipofectamine 2000 (Invitrogen) using the manufacturer’s suggested protocol with 300 ng each PE expression plasmid and 150 ng of each pegRNA expression plasmid on a cover slip. 48 h following transfection, transfection media was removed, cells were washed with 1× PBS and fixed with 4% formaldehyde in 1× PBS for 15 min at room temperature. Following blocking (blocking solution: 2% BSA, 0.3% Triton X-100, within 1× PBS), samples were stained with mouse antihemagglutinin (Sigma, H9658, 1:500), and Alexa 488 donkey anti-mouse IgG (H+L; Invitrogen, A-21202, 1:2000), sequentially. VECTASHIELD mounting medium with DAPI (Vector Laboratories, H-1200) was used to stain the nuclei and to mount the samples on the slide. Images were taken with ZEISS LSM 710 Confocal Microscope System. We used Fiji^44^ and the “CMCI-EMBL” plugin to calculate the signal from the nuclear or cytoplasm comparments as previously described^45^ (https://github.com/miura/NucleusRimIntensityMeasurementsV2/). We used >90 cells for each treatment group to determine the nuclear to cytoplasmic ratio for each PE construct, which was calculated as the intensity value from nucleus divided by the intensity value from cytoplasm and nucleus.

IHC staining was performed as previously described^42^ using beta-catenin antibody (BD, 610154, 1:100) or HA tag (CST, 3724, 1:400). The IHC profiler plugin^46,47^ for ImageJ software was used for the analysis of the nuclear fraction of PE2 or PE2* in 20x images of the stained liver sections by comparing the ratio of the intensity of the staining (pixels) in the whole cell (captured by the cytoplasmic contour) relative to the estimated staining within the nucleus of each cell (captured by the nuclear contour) using standard threshold. Each 20x image was divided into 4 quadrants for analysis using IHC profiler and the calculated nuclear signal ratio for each quadrant was averaged.

### Animal studies

All animal experiments were authorized by the Institutional Animal Care and Use Committee (IACUC) at UMASS medical school. No animals were excluded from the analyses. All prime editor plasmids were prepared by EndoFreeMaxi kit (Qiagen) and were delivered through hydrodynamic tail-vein injection. For PiZ correction, eight-week-old PiZ mice were injected with 2.3ml 0.9% saline containing 30μg PE2 or PE2* (n=3), 15μg pegRNA (*SERPINA1*) and 5μg Nicking sgRNA 3. For cancer model generation, FVB/NJ (Strain #001800) was purchased from Jackson Laboratories. Each FVB mouse (n=4) was injected with 2.3ml 0.9% saline containing 30μg PE2 or PE2*, 15μg pegRNA (*Ctnnb1*), 15μg Nicking sgRNA 2 (*Ctnnb1*), 5μg pT3 EF1a-MYC (a gift from Xin Chen, Addgene plasmid # 92046)^33^ and 1μg CMV-SB10 (a gift from Perry Hackett, Addgene plasmid # 24551).

### Deep sequencing and data analysis

Library construction for deep sequencing is modified from our previous report^27^. Briefly, 72 h after transfection or electroporation, cells were harvested and genomic DNA was extracted with GenElute Mammalian Genomic DNA Miniprep Kit (Sigma). Genomic loci spanning the target and off-target sites were PCR amplified with locus-specific primers carrying tails complementary to the Truseq adapters (Supplementary Table 3). 50 ng input genomic DNA was PCR amplified with Q5 High-Fidelity DNA Polymerase (New England Biolabs): (98 °C, 15 s; 67 °C 25 s; 72 °C 20 s) ×30 cycles. For the construction of the CCR5 UMI-based library, 50 ng input genomic DNA was first linearly pre-amplified with 10 nM final concentration 5p-CCR5_UMI primer using the Q5 High-Fidelity DNA Polymerase (New England Biolabs): (98 °C, 60 s; 67 °C, 25 s; 72 °C, 20 s) × 10 cycles. In the same reaction mix, 500 nM final concentration 5p-DS_constant and 3p-CCR5_DS primers were added for further amplification (98 °C, 60 s; 67 °C, 25 s; 72 °C, 20 s) for 30 cycles. Next, 0.1 μl of each PCR reaction was amplified with index-containing primers to reconstitute the TruSeq adaptors using the Q5 High-Fidelity DNA Polymerase (New England Biolabs): (98 °C, 15 s; 67 °C, 25 s; 72 °C, 20 s) x10 cycles. Equal amounts of the PCR products from each experimental condition (identified by different indices) were pooled and gel purified. The purified library was deep sequenced using a paired-end 150 bp Illumina MiniSeq run.

MiniSeq data analysis was done as previously reported ^27^. First, the quality of paired-end sequencing reads (R1 and R2 fastq files) was assessed using FastQC (http://www.bioinformatics.babraham.ac.uk/projects/fastqc/). Raw paired-end reads were combined using paired end read merger (PEAR) (PMID: 24142950) to generate single merged high-quality full-length reads. Reads were then filtered by quality (using Filter FASTQC (PMID: 20562416)) to remove those with a mean PHRED quality score under 30 and a minimum per base score under 24. Each group of reads was then aligned to a corresponding reference sequence using BWA (version 0.7.5) and SAMtools (version 0.1.19). To determine indel frequency, size and distribution, all edited reads from each experimental replicate were combined and aligned, as described above. Indel types and frequencies were then cataloged in a text output format at each base using bam-readcount (https://github.com/genome/bam-readcount). For each treatment group, the average background indel frequencies (based on indel type, position and frequency) of the triplicate negative control group were subtracted to obtain the precise editing and indel frequencies for each group. The fraction of precise editing is calculated as sequencing reads with the desired allele editing/ all reads for the target locus. The results were concatenated and loaded into GraphPad Prism 8.4 for data visualization.

### Tn5 tagmentation and library preparation for UDiTaS

For tagmentation, transposome was assembled as previously described^39^ using purified Tn5 protein and oligonucleotides purchased from IDT. 200ng of genomic DNA was incubated with 2ul of assembled transposome at 55 degree for 7 mins, and the product was cleaned up (20ul) with a Zymo column (Zymo Research, #D4013). Tagmented DNA was used for the 1st PCR using PlatinumTM SuperFi DNA polymerase (Thermo) with i5 primer and gene specific primers (Supplementary table 3). Two different libraries were prepared for gDNA from each mouse with different combinations of primers (i5+Locus_F [UDiTaS], i5+Locus_R [UDiTaS]). The i7 index was added in the 2nd PCR and the PCR product was cleaned up with Ampure XP SPRI beads (Agencourt, 0.9X reaction volume). Completed libraries were quantified by Tapestation and Qubit (Agilent), pooled with equal mole and sequenced with 150 bp paired-end reads on an Illumina MiniSeq instrument.

### UDiTaS data analysis

The analysis pipeline was built using python code. Briefly, the analysis steps are as follows:

i. Demultiplexing. Raw BCL files were converted and demultiplexed using the appropriate sequencing barcodes, allowing up to one mismatch in each barcode. Unique molecular identifiers (UMIs) for each read were extracted for further downstream analysis.
ii. Trimming. Remove 3′ adapters using cutadapt, version 3.0 http://journal.embnet.org/index.php/embnetjournal/article/view/200/479
iii. Create reference sequence based the UDiTaS locus-specific primer position and AAV plasmid map separately. Build index files for the reference using bowtie2-index^48^, version 2.4.0.
iv. Alignment analysis. Paired reads were then globally aligned (end-to-end mode) to all the reference amplicons using bowtie2’s very sensitive parameter. Finally, samtools^49^ (version 0.1.19) was used to create and index sorted bam files. Paired-end reads covering a window between pegRNA targeting site and nicking sgRNA targeting sites were extracted and the total number of unique UMIs were counted. Precise editing or small indels were analyzed as previously described^27^. Pindel^50^ (version 0.2.5b8) was used to detect breakpoints of large deletions. Raw sequencing reads that align to the reference sequence were collapse to a single read by common UMI and categorized as an exemplar for each UMI to a specific category—for example, Wild Type, precise editing, small indel/substitution and Large Deletions. Then the number of UMIs assigned per category is determined to define the ratio of each event.
v. AAV integration. Extract the unmapped reads that did not locally align to the AAV/plasmid in steps 3 and 4 using bedtools bamtofastq. With bowtie2, index the AAV plasmid sequence and then do a local alignment of the reads. Of the reads that locally align to the AAV plasmid, first filter out those reads which are directly adjacent to the UDiTaS primer (on read 2) and do not contain any target locus sequence. This removes reads that are due to false priming. Of the remaining reads, collapse these by UMI and count the UMIs. Classify the exemplar read for each UMI as ‘AAV/Plasmid Integrations’.

### Statistical analysis

The fold changes of editing (precise editing or indels) are calculated between the corresponding groups: pegRNA_only between PE2 and PE2*, or with specific Nicking sgRNA between PE2 and PE2*. All raw data is listed in Source data file and statistical analyses were performed using GraphPad Prism 8.4. Sample size was not pre-determined by statistical methods, but rather, based on preliminary data. Group allocation was performed randomly. In all studies, data represent biological replicates (n) and are depicted as mean ± s.d. as indicated in the figure legends. Comparison of mean values was conducted with one-way ANOVA with Tukey’s multiple comparisons test, as indicated in the figure legends. In all analyses, P values < 0.05 were considered statistically significant.

## Data availability

Illumina Sequencing data have been submitted to the Sequence Read Archive. These datasets are available under BioProject Accession number PRJNA692762 [https://www.ncbi.nlm.nih.gov/bioproject/PRJNA692762/]. The authors declare that all other data supporting the findings of this study are available within the paper and its Supplementary Information files or upon reasonable request. Backbone plasmids used for pegRNA and sgRNA cloning are available from addgene. Source data are provided with this paper.

## Code availability

The software used for data analysis is available at Github:

https://github.com/locusliu/GUIDESeq-Preprocess_from_Demultiplexing_to_Analysis

https://github.com/editasmedicine/uditas

https://github.com/ericdanner/REPlacE_Analysis

https://github.com/locusliu/PCR_Amplicon_target_deep_seq/blob/master/CRESA-lpp.py

## Acknowledgements

We thank C. Mello and P. Zamore for helpful discussions. We thank Y. Liu, J. Xie, and Q. Su in the UMass Morphology and Viral Vector Cores for support. W.X. was supported by grants from the National Institutes of Health (DP2HL137167, P01HL131471 and UG3HL147367), American Cancer Society (129056-RSG-16-093), the Lung Cancer Research Foundation, and the Cystic Fibrosis Foundation. P.Z., E.M., K.P. and S.A.W. were supported in part by the National Institutes of Health (R01GM115911, and UG3TR002668) and the Rett Syndrome Research Trust. A.M. and E.J.S. were supported by UG3TR002668.

## Author contributions

PL and SQL performed the experiments, analyzed data and wrote the manuscript. CZ, EM, YGZ, KP, and AM performed experiments. GG, EJS, TRF, SAW, and WX supervised the study and wrote the manuscript with all co-authors.

## Supplementary data accompanies this paper

### Figure Legends

**Supplementary Fig. 1.**
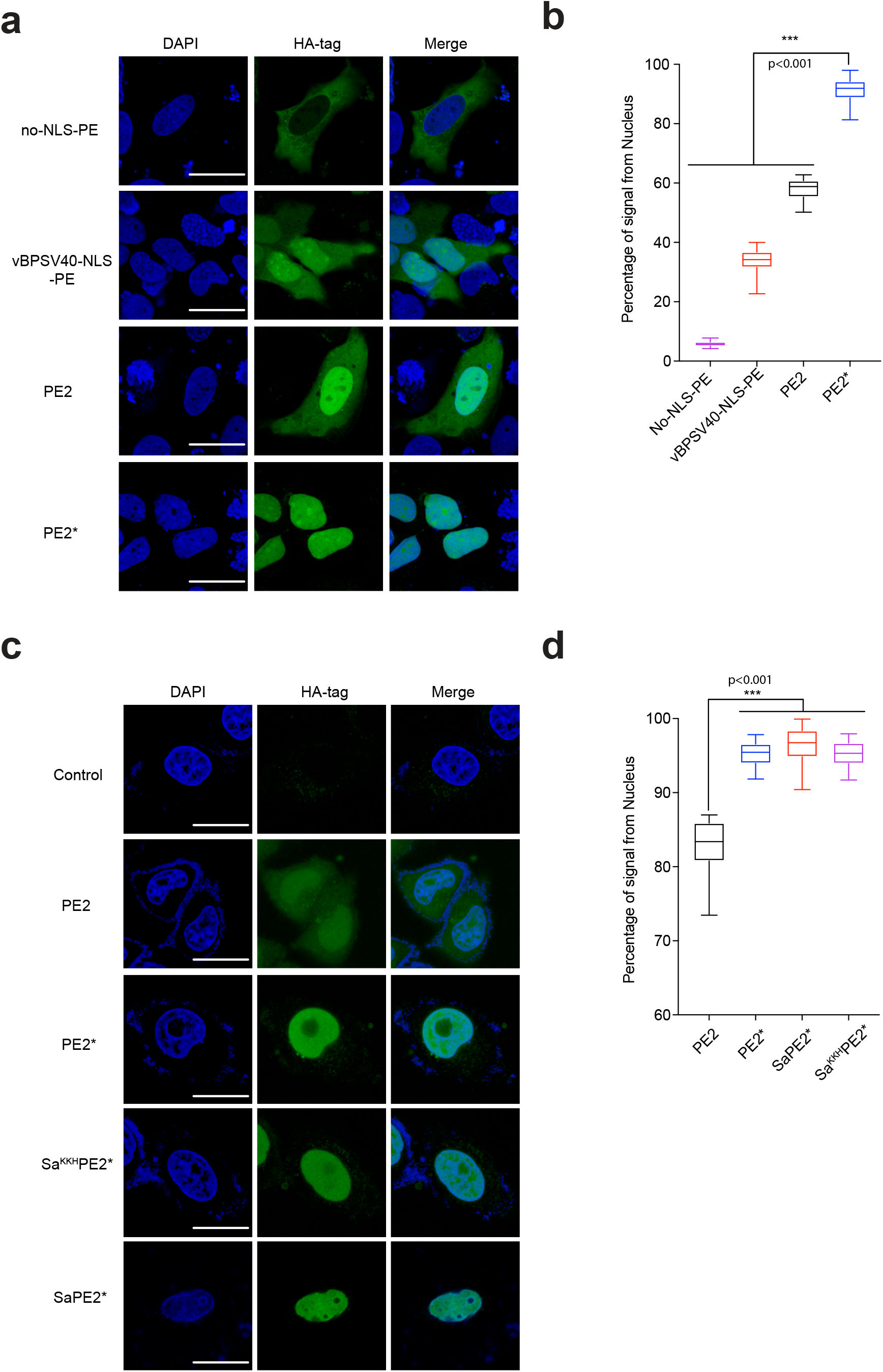
Optimized prime editor increases the degree of nuclear import. (a) Representative immunofluorescence images of prime editors transfected into U2OS cells immunostained with a HA tag antibody to visualize subcellular localization. PE2 contains two Bipartite SV40 NLSs, whereas the PE2* variant include an extra N-terminal c-Myc NLS and a C-terminal variant bipartite SV40 NLS (vBPSV40) and SV40 NLS at the C-terminus. DNA was stained with DAPI. Scale bar, 10μm. (b) Average percentages of HA-tag signal from nucleus for different prime editors. (c-d) Representative immunofluorescence images and quantification in HeLa cells. For (b) and (d), >90 cells were analyzed for each group from 3 independent experiments. \*\*\**P*<0.001 by one-way ANOVA with Tukey’s multiple comparisons test. In the boxes, the top, middle and bottom lines represent the 25, 50 and 75 percentiles, respectively. Whiskers indicated the min and max percentiles and outliers are not shown. Scale bar, 10µm.

**Supplementary Fig. 2.**
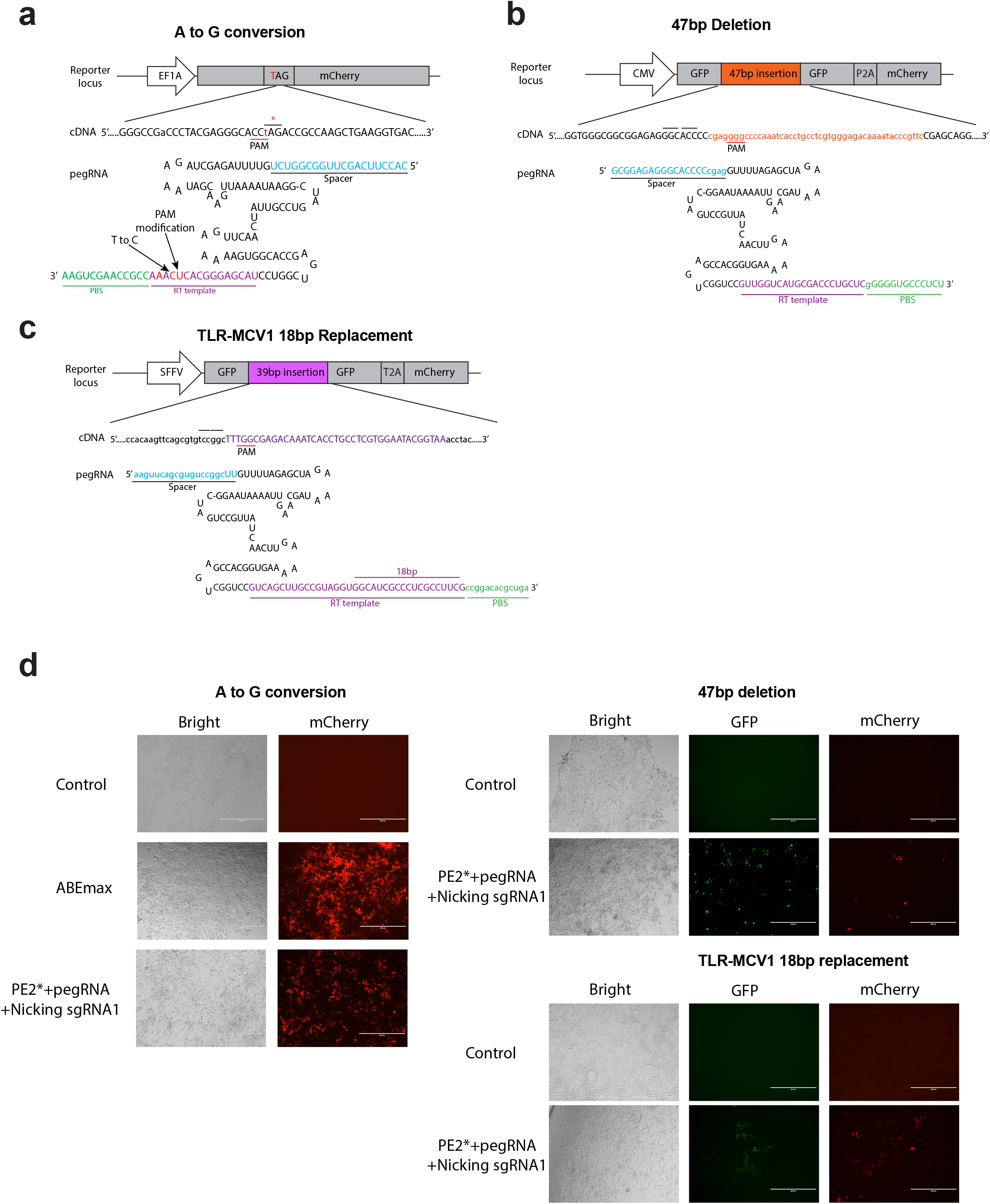
Prime editing in reporter cells by PE2 and PE2*. (a) Sequence of reporter locus and pegRNA used for repair of mCherry reporter in HEK293T cells. Bar above the cDNA indicates the stop codon with the target “t” for conversion indicated in red. Two additional silent mutations are included to discourage recutting of the repaired DNA sequence (b) Sequence of reporter and pegRNA used for generation of a 47bp deletion to restore function of the GFP reporter in HEK293T cells. The bars above the cDNA indicates three nucleotide blocks that correspond to codons in the GFP reporter. (c) Sequence of reporter and pegRNA used for replacement of a 18bp element to restore function of the GFP reporter. The bars above the cDNA indicates three nucleotide blocks that correspond to codons in the GFP reporter. (d) Representative images of HEK293T reporter cells (n=3) transfected with control, ABEmax or PE2*. Scale bar, 400μm.

**Supplementary Fig. 3.**
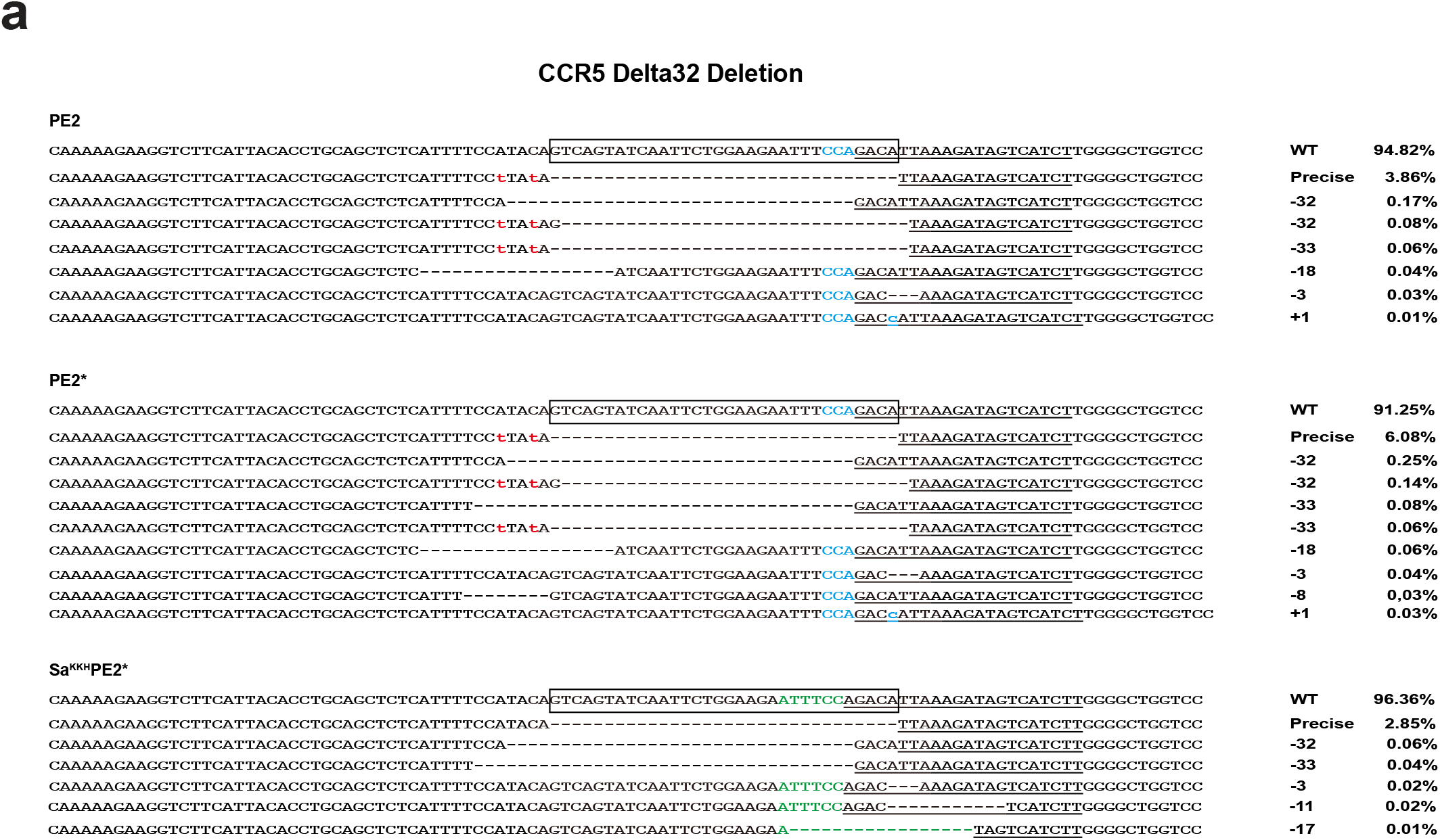
Sequencing analysis of the *CCR5* prime editing by PE2, PE2*, or Sa^KKH^PE2*. The percentage of most common sequences in PE2, PE2* and Sa^KKH^PE2* transfected cells is shown on the right (representative of n=3, determined by Illumina sequencing). The PE target site is underlined. The PAM sequences are indicated in light blue for SpCas9 and green for SaCas9^KKH^. The box denotes the 32bp sequence deleted in CCR5delta32. Two “t” mutations in red were included in the PE2 and PE2* RT template (Fig. 2d) to demonstrate that sequence collapse was not a function of nuclease-induced microhomology mediated deletion and to reduce re-cutting of deletion allele. Deleted bases are indicated by dashes and inserted bases are in blue.

**Supplementary Fig. 4.**
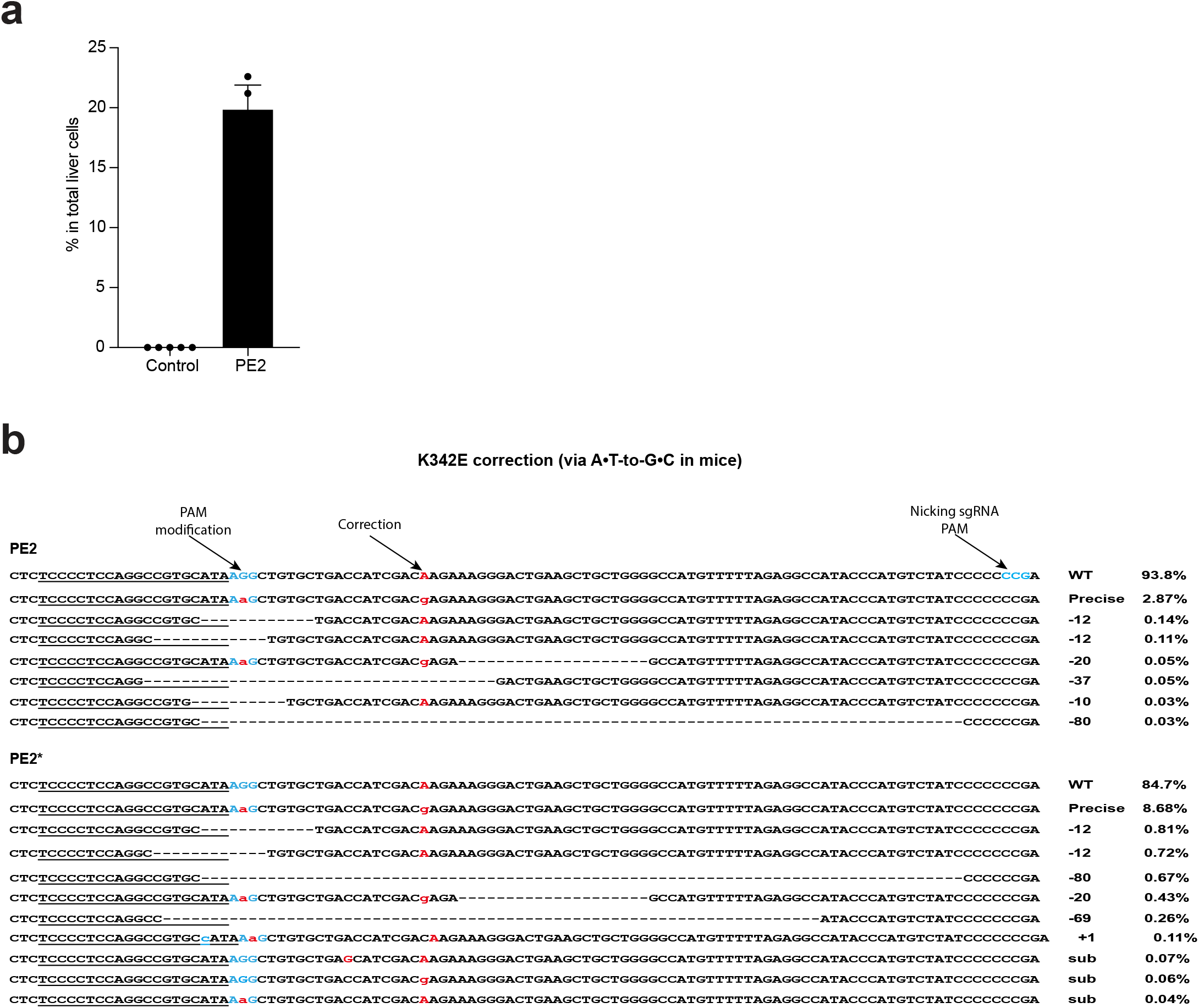
Sequencing analysis of *SERPINA1* editing in the liver of PiZ mouse. (a)Evaluating prime editor expression in mouse liver. FVB mice were injected with 30μg of control vector or PE2 plasmid with an incorporated 3xHA-tag. Livers were harvested at day 2 and IHC staining were performed with an HA-tag antibody. Quantification of HA^+^ cells. Numbers are mean ± sem (n=5 images from 4 mice for each group). (b)The percentage of most common sequences in the liver of PE2 and PE2*-treated mice is shown on the right (representative liver of n=3, determined by Illumina sequencing). The PE target site is underlined. The PAM sequences are in light blue. Nucleotide substitutions are labeled in red. Deleted bases are indicated by dashes. Inserted bases are shown in blue/lower case.

**Supplementary Fig. 5.**
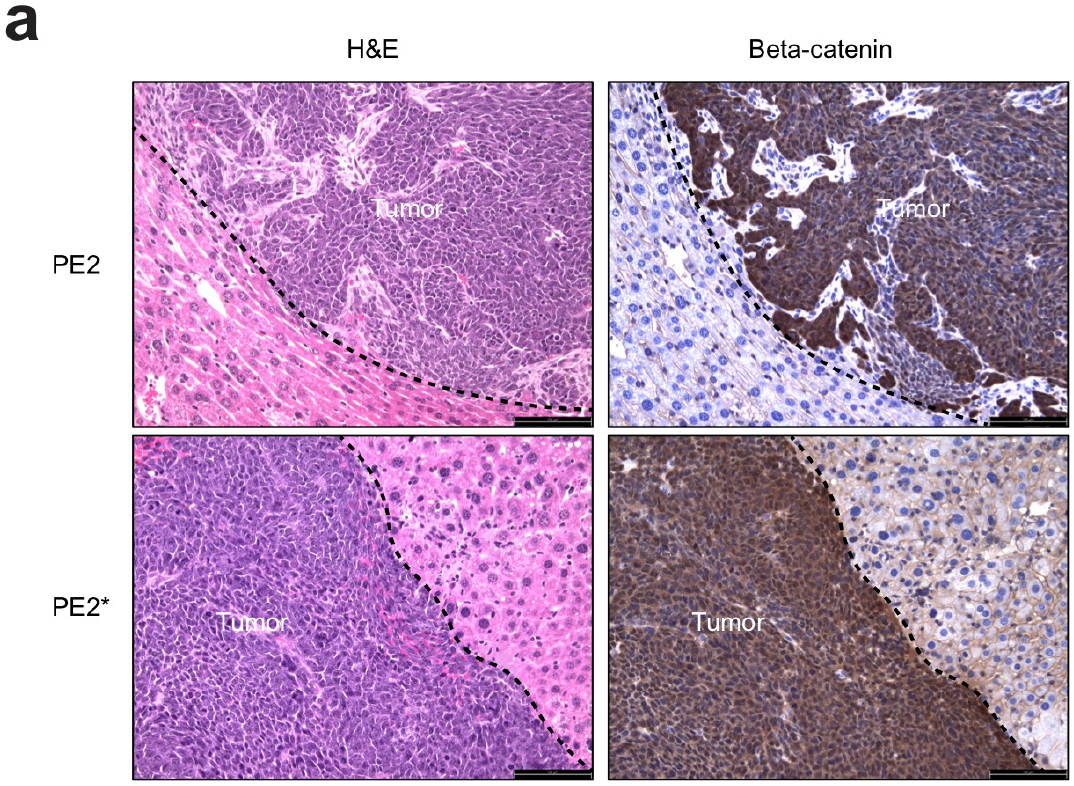
Liver tumors are positive for nuclear beta-Catenin. (a), Representative H&E and beta-catenin IHC staining (n=3) in PE2 or PE2*-induced S45F tumors in Fig. 4. Scale bars: 100 µm (20X lens).

**Supplementary Fig. 6.**
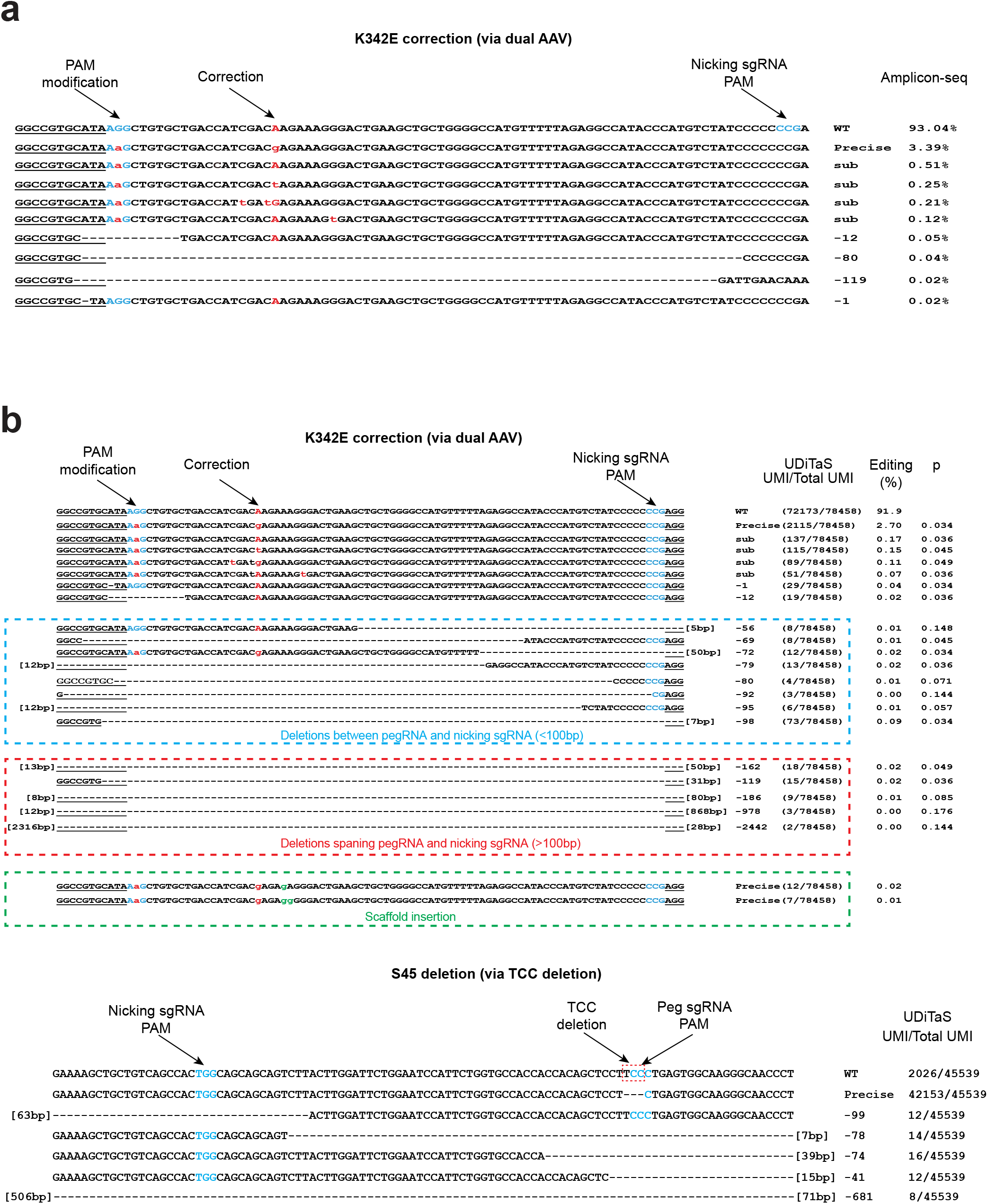
Sequencing analysis of the *Ctnnb1* and *SERPINA1* prime editing by PE2 or PE2*. (a), The percentage of most common sequences in the liver of dual AAV-treated PiZ mice determined by locus amplification is shown on the right (representative liver of n=3). A portion of the PE target site is underlined. The PAM sequences are in light blue. Nucleotide substitutions are labeled in red. Deleted bases are indicated by dashes. (b), Length distribution of precise editing and other indels at *SERPINA1 (K342E correction)* and *Ctnnb1* target sites by UDiTaS. Also included are sequence modifications that may be associated with pegRNA scaffold insertions^7^. Data for a representative liver is shown (n=3 mice).

## Abbreviation

CRISPR/Cas9: clustered regularly interspaced short palindromic repeats/CRISPR-associated protein 9
PEs: prime editors (PEs)
pegRNA: prime editing guide RNA
PBS: primer binding site
RT: reverse transcriptase

