## Supplementary information for "Improved prime editors enable pathogenic allele correction and cancer modelling in adult mice"

**Supplementary Table 1**. Sequences of pegRNAs and sgRNAs used in this study. All sequences are shown in 5' to 3' orientation.

**Supplementary Table 2**. Sequences of primers used for pegRNA cloning

**Supplementary Table 3**. Sequences of primers used for genomic DNA amplification and high throughput sequencing.

**Supplementary Table 4.** Potential genetic engineered liver cancer models using prime editors.

**Supplementary Table 5.** In vivo on target analysis at SERPINA1 site by UdiTaS.

**Supplementary Sequences 1.** Sequences of reporter cell line used in this study

**Supplementary Sequences 2.** Sequence of backbone plasmid used for pegRNA and nicking sgRNA cloning

**Supplementary Sequences 3.** Sequences of prime editors

**Supplementary Note 1**. FACS gating examples for GFP-positive or Cherry-positive cells.

**Supplementary Table 1**. Sequences of pegRNAs and sgRNAs used in this study. All sequences are shown in 5' to 3' orientation.

sgRNA scaffold

GTTTTAGAGCTAGAAATAGCAAGTTAAAATAAGGCTAGTCCGTTATCAACTTGAAAAAGTGGGACCGAGTCGGTCC

| **pegRNA** | **spacer sequence (5'-3')** | **3' extension** | **PBS (nt)** | **RT (nt)** | **Figure** | **PE** |
|---|---|---|---|---|---|---|
| mCherry A to G | CACCTTCAGCTTGGCGGTCT | TACGAGGGCACTCAAACCGCCAAGCTGAAG | 14 | 16 | Figure 1B | PE2 or PE2* |
| mCherry A to G | GGTCACCTTCAGCTTGGCGGT | TACGAGGGCACCCAGACTGCCAAGCTGAAGGTGA | 16 | 17 | Figure 1B | Sa$^{KKH}$PE2* |
| GFP-insertion | AAGTTCAGCGTGTCCGGCTT | GTCAGCTTGCCGTAGGTGGCATCGCCCTCGCCTTCG | 13 | 36 | Figure 1C | PE2 or PE2* |
| GFP-insertion | TGAACTTCAGGGTCAGCTTGCC | ACAAGTTCAGCGTGTCCGGCGAGGGCGAGGGCGATGCCACGTACGGCAAGCTGACCCTGAAGTTC | 18 | 47 | Figure 1C | Sa$^{KKH}$PE2* |
| GFP-deletion | GCGGAGAGGGCACCCCCGA | GTTGGTCATGCGACCCTGCTCGGGGGTGCCCTCTCC | 14 | 22 | Figure 1D | PE2 or PE2* |
| GFP-deletion | GGTCATGCGACCCTGCTCGGA | CGGAGAGGGCACCCCCGAGCAGGGTCGCATG | 15 | 16 | Figure 1D | Sa$^{KKH}$PE2* |
| EMX1 +5 G to T | GAGTCCGAGCAGAAGAAGAA | ATGGGAGCACTTCTTCTTCTGCTC | 14 | 10 | Figure 2A | PE2 or PE2* |
| EMX1 +3 +8 G to T | CAGAAGCTGGAGGAGGAAGGGC | TGCTCGGAATCAGACCCTTCCTCCTCCAGCT | 15 | 16 | Figure 2A | SaCas9 PE2* or Sa$^{KKH}$PE2* |
| EMX1 +4 3bp deletion | GAGTCCGAGCAGAAGAAGAA | ATGTGATGGGAGTTCTTCTTCTGCTC | 14 | 12 | Figure 2B | PE2 or PE2* |
| EMX1 +1 3bp deletion | CAGAAGCTGGAGGAGGAAGGGC | TTCTGCTCGGACTCAGCTTCCTCCTCCAGCT | 15 | 16 | Figure 2B | SaCas9 PE2* or Sa$^{KKH}$PE2* |
| EMX1 +4 6bp Insertion | GAGTCCGAGCAGAAGAAGAA | GAGCAGAAGAAGAAAAGCTTGGGCTCCCATCACAT | 14 | 21 | Figure 2C | PE2 or PE2* |
| EMX1 +1 6bp Insertion | CAGAAGCTGGAGGAGGAAGGGC | TGCTCGGACTCAGGCCAAGCTTCTTCCTCCTCCAGCT | 15 | 22 | Figure 2C | SaCas9 PE2* or Sa$^{KKH}$PE2* |
| CCR5-delta32-deletion | AGATGACTATCTTTAATGTC | ATTACACCTGCAGCTCTCATTTTCCTTATATTAAAGATAGTCATC | 16 | 29 | Figure 2D | PE2 or PE2* |
| CCR5-delta32-deletion | AAGATGACTATCTTTAATGTCT | AGCTCTCATTTTCCATACATTAAAGATAGTCATC | 17 | 17 | Figure 2D | Sa$^{KKH}$PE2* |
| Serpina1 G to A | TCCCCTCCAGGCCGTGCATA | TCTTGTCGATGGTCAGCACAGCTTTATGCACGGCCTGGAG | 13 | 27 | Figure 3A | PE2 or PE2* |
| Serpina1 G to A | CAGCTTCAGTCCCTTTCTCGT | ATCGACAAGAAAGGGACTGAAGCT | 15 | 9 | Figure 3A | Sa$^{KKH}$PE2* |
| Serpina1 A to G | TCCCCTCCAGGCCGTGCATA | TCTCGTCGATGGTCAGCACAGCTTTATGCACGGCCTGGAG | 13 | 27 | Figure 3B | PE2 or PE2* |
| Ctnnb1 C to T | AGGGTTGCCCTTGCCACTCA | GCTCCTTTCCTGAGTGGCAAGGGCAA | 13 | 13 | Figure 4A | PE2 or PE2* |
| Ctnnb1 C to T | AGGGTTGCCCTTGCCACTCA | ACAGCTCCTTTGAGTGGCAAGGGCAA | 13 | 13 | Figure 4D | PE2* |

| **Nicking sgRNA** | **spacer sequence (5'-3')** | **Figure** | **PE** |
|---|---|---|---|
| mCherry A to G | GCGCTTCAAGGTGCACATGGA | Figure 1B | PE2 or PE2* |
| mCherry A to G | GCTGTCCCCTCAGTTCATGTA | Figure 1B | PE2 or PE2* |
| mCherry A to G | GATGGAGGGCTCCGTGAACGGCC | Figure 1B | Sa$^{KKH}$PE2* |
| mCherry A to G | GTTCGCCTGGGACATCCTGTCCC | Figure 1B | Sa$^{KKH}$PE2* |
| GFP-insertion-sp-NK1 | GTAGGTCAGGGTGGTCACGA | Figure 1C | PE2 or PE2* |
| GFP-insertion-sp-NK2 | GCTCCTCGCCCTTGCTCACCA | Figure 1C | PE2 or PE2* |
| GFP-insertion-saKKH-NK1 | GCAAGGGCGAGGAGCTGTTCAC | Figure 1C | Sa$^{KKH}$PE2* |

| | | | |
|---|---|---|---|
| GFP-insertion-saKKH-NK2 | GTGACCACCCTGACCTACGGCG | Figure 1C | $Sa^{KKH}PE2^*$ |
| GFP-deletion-sp-NK1 | GAGAAGCCGTAGCCCATCACG | Figure 1D | PE2 or PE2* |
| GFP-deletion-sp-NK2 | GATCTTCATGGCGGGCATGG | Figure 1D | PE2 or PE2* |
| GFP-deletion-saKKH-NK1 | GATCACCGGCACCCTGAACGGCG | Figure 1D | $Sa^{KKH}PE2^*$ |
| GFP-deletion-saKKH-NK2 | GCAGCCCCTACCTGCTGAGCCA | Figure 1D | $Sa^{KKH}PE2^*$ |
| EMX1 +5 G to T-sp-NK1 | GTTGCCCACCCTAGTCATTGG | Figure 2A | PE2 or PE2* |
| EMX1 +5 G to T-sp-NK2 | GGCCGTTTGTACTTTGTCCTC | Figure 2A | PE2 or PE2* |
| EMX1 +3 +8 G to T-sa-NK1 | GCCTGGGCCAGGGAGGGAGGGGC | Figure 2A | SaCas9 PE2* or $Sa^{KKH}PE2^*$ |
| EMX1 +3 +8 G to T-sa-NK2 | GTGGTTGCCCACCCTAGTCATT | Figure 2A | SaCas9 PE2* or $Sa^{KKH}PE2^*$ |
| EMX1 +4 3bp deletion-sp-NK1 | GTTGCCCACCCTAGTCATTGG | Figure 2B | PE2 or PE2* |
| EMX1 +4 3bp deletion-sp-NK2 | GGCCGTTTGTACTTTGTCCTC | Figure 2B | PE2 or PE2* |
| EMX1 +1 3bp deletion-sa-NK1 | GCCTGGGCCAGGGAGGGAGGGGC | Figure 2B | SaCas9 PE2* or $Sa^{KKH}PE2^*$ |
| EMX1 +1 3bp deletion-sa-NK2 | GTGGTTGCCCACCCTAGTCATT | Figure 2B | SaCas9 PE2* or $Sa^{KKH}PE2^*$ |
| EMX1 +4 6bp Insertion-sp-NK1 | GTTGCCCACCCTAGTCATTGG | Figure 2C | PE2 or PE2* |
| EMX1 +4 6bp Insertion-sp-NK2 | GGCCGTTTGTACTTTGTCCTC | Figure 2C | PE2 or PE2* |
| EMX1 +1 6bp Insertion-sa-NK1 | GCCTGGGCCAGGGAGGGAGGGGC | Figure 2C | SaCas9 PE2* or $Sa^{KKH}PE2^*$ |
| EMX1 +1 6bp Insertion-sa-NK2 | GTGGTTGCCCACCCTAGTCATT | Figure 2C | SaCas9 PE2* or $Sa^{KKH}PE2^*$ |
| CCR5-deletion-sp-NK1 | GCTGTGTTTGCGTCTCTCCC | Figure 2E | PE2 or PE2* |
| CCR5-deletion-sp-NK2 | GACAAGTGTGATCACTTGGG | Figure 2E | PE2 or PE2* |
| CCR5-deletion-sp-NK3 | GCAGGACGGTCACCTTTGGGG | Figure 2E | PE2 or PE2* |
| CCR5-deletion-saKKH-NK1 | GCATCTTTACCAGATCTCAAAAA | Figure 2E | $Sa^{KKH}PE2^*$ |
| CCR5-deletion-saKKH-NK2 | GTGGCTGTGTTTGCGTCTCTCCC | Figure 2E | $Sa^{KKH}PE2^*$ |
| Serpina1 G to A-sp-NK1 | GGGGGGGATAGACATGGGTA | Figure 3A | PE2 or PE2* |
| Serpina1 G to A-sp-NK2 | GACCTCGGGGGGGATAGACA | Figure 3A | PE2 or PE2* |
| Serpina1 G to A-sp-NK3 | GGGTTTGTTGAACTTGACCT | Figure 3A | PE2 or PE2* |
| Serpina1 G to A-sp-NK4 | GTTCAATCATTAAGAAGACAA | Figure 3A | PE2 or PE2* |
| Serpina1 G to A | GCACGTGAGCCTTGCTCGAGGCC | Figure 3A | $Sa^{KKH}PE2^*$ |
| Serpina1 G to A | GCCCATGTCTATCCCCCCCGAGG | Figure 3A | $Sa^{KKH}PE2^*$ |
| Serpina1 A to G (mouse_injection) | GGGTTTGTTGAACTTGACCT | Figure 3B | PE2 or PE2* |
| Ctnnb1 C to T | GAAAAGCTGCTGTCAGCCAC | Figure 4A | PE2 or PE2* |

**Supplementary Table 2**. Sequences of primers used for pegRNA cloning

| pegRNA | Sequence (5'-3') | Figure | PE |
|---|---|---|---|
| mCherry A to G_F | TATCTTGTGGAAAGGACGAAACACCGCACCTTCAGCTTGGCGGTCTGTTTTA GAGCTAGAAATAG | Figure 1B | PE2 or PE2* |
| mCherry A to G_R | CGGTATCGATAAGCTTGATATCGAATTCAAAAAACTTCAGCTTGGCGGTTTG AGTGCCCTCGTAGGACCGACTCGGTCCCACTTTTTC | Figure 1B | PE2 or PE2* |
| mCherry A to G_F | TATCTTGTGGAAGGACGAAACACCGGTCACCTTCAGCTTGGCGGTGTTTTAG TACTCTGGAAACAG | Figure 1B | $Sa^{KKH}PE2^*$ |
| mCherry A to G_R | CGACGGTATCGATAAGCTTGATATCGAATTCAAAAAAATCACCTTCAGCTTG GCAGTCTGGGTGCCCTCGTATCTCGCCAACAAGTTGACGAGAT | Figure 1B | $Sa^{KKH}PE2^*$ |
| GFP-insertion_F | TGGAAAGGACGAAACACCGAAGTTCAGCGTGTCCGGCTTGTTTTAGAGCTAG AAATAG | Figure 1C | PE2 or PE2* |
| GFP-insertion_R | GTATCGATAAGCTTGATATCAAAAAATCAGCGTGTCCGGCGAAGGCGAGGGC GATGCCACCTACGGCAAGCTGACGGACCGACTCGGTCCCACTTTTTC | Figure 1C | PE2 or PE2* |
| GFP-insertion_F | GTGGAAGGACGAAACACCGTGAACTTCAGGGTCAGCTTGCCGTTTTAGTACT CTGGAAACAGA | Figure 1C | $Sa^{KKH}PE2^*$ |

| | | | |
|---|---|---|---|
| GFP-insertion_R | ATCTCGTCAACTTGTTGGCGAGAACAAGTTCAGCGTGTCCGGCGAGGGCGAG GGCGATGCCACGTACGGCAAGCTGACCCTGAAGTTCTTTTTTTGAATTCGAT ATCAAGCTTATCGATACCGTCG | Figure 1C | Sa$^{KKH}$PE2* |
| GFP-deletion_F | TGGAAAGGACGAAACACCGGCGGAGAGGGCACCCCCGAGTTTTAGAGCTAGA AATAG | Figure 1D | PE2 or PE2* |
| GFP-deletion_R | GTATCGATAAGCTTGATATCAAAAAAGGAGAGGGCACCCCCGAGCAGGGTCG CATGACCAACGGACCGACTCGGTCCCACTTTTTCAAG | Figure 1D | PE2 or PE2* |
| GFP-deletion_F | GTGGAAGGACGAAACACCGGGTCATGCGACCCTGCTCGGAGTTTTAGTACTC TGGAAACAG | Figure 1D | Sa$^{KKH}$PE2* |
| GFP-deletion_R | CGACGGTATCGATAAGCTTGATATCGAATTCAAAAAAACATGCGACCCTGCT CGGGGGTGCCCTCTCCGTCTCGCCAACAAGTTGACGAGAT | Figure 1D | Sa$^{KKH}$PE2* |
| EMX1 +5 G to T-sp_F | TGGAAAGGACGAAACACCGGAGTCCGAGCAGAAGAAGAAGTTTTAGAGCTAG AAATAG | Figure 2A | PE2 or PE2* |
| EMX1 +5 G to T-sp_R | GTATCGATAAGCTTGATATCAAAAAAGAGCAGAAGAAGAAGTGCTCCCATGG ACCGACTCGGTCCCACTTTTTC | Figure 2A | PE2 or PE2* |
| EMX1 +3 +8 G to T-sa_F | GTGGAAGGACGAAACACCGCAGAAGCTGGAGGAGGAAGGGCGTTTTAGTACT CTGGAAACAG | Figure 2A | SaCas9 PE2* or Sa$^{KKH}$PE2* |
| EMX1 +3 +8 G to T-sa_R | CGACGGTATCGATAAGCTTGATATCGAATTCAAAAAAAAGCTGGAGGAGGAA GGGTCTGATTCCGAGCATCTCGCCAACAAGTTGACGAGAT | Figure 2A | SaCas9 PE2* or Sa$^{KKH}$PE2* |
| EMX1 +4 3bp deletion-sp_F | TGGAAAGGACGAAACACCGGAGTCCGAGCAGAAGAAGAAGTTTTAGAGCTAG AAATAG | Figure 2B | PE2 or PE2* |
| EMX1 +4 3bp deletion-sp_R | GTATCGATAAGCTTGATATCAAAAAAGAGCAGAAGAAGAACTCCCATCACAT GGACCGACTCGGTCCCACTTTTTC | Figure 2B | PE2 or PE2* |
| EMX1 +1 3bp deletion-sa_F | GTGGAAGGACGAAACACCGCAGAAGCTGGAGGAGGAAGGGCGTTTTAGTACT CTGGAAACAG | Figure 2B | SaCas9 PE2* or Sa$^{KKH}$PE2* |
| EMX1 +1 3bp deletion-sa_R | CGACGGTATCGATAAGCTTGATATCGAATTCAAAAAAAAGCTGGAGGAGGAA GCTGAGTCCGAGCAGAATCTCGCCAACAAGTTGACGAGAT | Figure 2B | SaCas9 PE2* or Sa$^{KKH}$PE2* |
| EMX1 +4 6bp Insertion-sp_F | TGGAAAGGACGAAACACCGGAGTCCGAGCAGAAGAAGAAGTTTTAGAGCTAG AAATAG | Figure 2C | PE2 or PE2* |
| EMX1 +4 6bp Insertion-sp_R | GTATCGATAAGCTTGATATCAAAAAAGAGCAGAAGAAGAAAAGCTTGGGCTC CCATCACATGGACCGACTCGGTCCCACTTTTTC | Figure 2C | PE2 or PE2* |
| EMX1 +1 6bp Insertion-sa_F | GTGGAAGGACGAAACACCGCAGAAGCTGGAGGAGGAAGGGCGTTTTAGTACT CTGGAAACAG | Figure 2C | SaCas9 PE2* or Sa$^{KKH}$PE2* |
| EMX1 +1 6bp Insertion-sa_R | CGACGGTATCGATAAGCTTGATATCGAATTCAAAAAAAAGCTGGAGGAGGAA GAAGCTTGGCCTGAGTCCGAGCATCTCGCCAACAAGTTGACGAGAT | Figure 2C | SaCas9 PE2* or Sa$^{KKH}$PE2* |
| CCR5-deletion-sp_F | TGGAAAGGACGAAACACCGAGATGACTATCTTTAATGTCGTTTTAGAGCTAG AAATAG | Figure 2E | PE2 or PE2* |
| CCR5-deletion-sp_R | GTATCGATAAGCTTGATATCAAAAAAGATGACTATCTTTAATATAAGGAAAA TGAGAGCTGCAGGTGGACCGACTCGGTCCCACTTTTTC | Figure 2E | PE2 or PE2* |
| CCR5-deletion-saKKH_F | GTGGAAGGACGAAACACCGAAGATGACTATCTTTAATGTCTGTTTTAGTACT CTGGAAACAG | Figure 2E | Sa$^{KKH}$PE2* |
| CCR5-deletion-saKKH_R | GTATCGATAAGCTTGATATCAAAAAAGATGACTATCTTTAATATAAGGAAAA TGAGAGCTGCAGGTGTAATGGACCGACTCGGTCCCACTTTTTC | Figure 2E | Sa$^{KKH}$PE2* |
| Serpina1 G to A gblocks | TATATATCTTGTGGAAAGGACGAAACACCGtTATATATCTTGTGGAAAGGAC GAAACACCGtcccctccaggccgtgcatagttttagagctagaaatagcaag ttaaaataaggctagtccgttatcaacttg | Figure 3A | PE2 or PE2* |
| Serpina1 G to A gblocks | TATATATCTTGTGGAAAGGACGAAACACCGcaTATATATCTTGTGGAAAGGA CGAAACACCGtcccctccaggccgtgcatagttttagagctagaaatagcaa gttaaaataaggctagtccgttatcaacttg | Figure 3A | Sa$^{KKH}$PE2* |
| Serpina1 A to G gblocks | TATATATCTTGTGGAAAGGACGAAACACCGTATATATCTTGTGGAAAGGACG AAACACCGtcccctccaggccgtgcatagttttagagctagaaatagcaagt taaaataaggctagtccgttatcaacttg | Figure 3B | PE2 or PE2* |
| Ctnnb1 C to T gblocks | ATCTTGTGGAAAGGACGAAACACCGAGGGTTGCCCTTGCCACTCAgttttag agctagaaatagcaagttaaaataaggctagtccgttatcaacttgaaaaag tgggaccgagtcggtccGCTCCTTTCCTGAGTGGCAAGGGCAATTTTTTTGA ATTCGATATCAAGCTTATCGATACCGT | Figure 4A | PE2 or PE2* |

**Supplementary Table 3**. Sequences of primers used for genomic DNA amplification and high throughput sequencing.

| Figure | Description | PE | | Squence |
|---|---|---|---|---|
| Figure 2A~2C | EMX1_locus | PE2/PE2* or SaCas9 | 5p-DS1_EMX1 | ctacacgacgctcttccgatctCCTCCTGAGT TTCTCATCTGTGCC |
| Figure 2A~2C | | | 3p-DS1_EMX1 | agacgtgtgctcttccgatctTCTGCCCTCGT GGGTTTGTG |

| | | | | |
|---|---|---|---|---|
| Figure 2A~2C | | PE2*/SaKKH PE2* | 5p-DS2_EMX1 | ctacacgacgctcttccgatctGGACAAAGTA CAAACGGCAGAAGC |
| Figure 2A~2C | | | 3p-DS2_EMX1 | agacgtgtgctcttccgatctCAGCCAGCCCA TTGCTTGTC |
| Figure 2D~2E | CCR5-deletion | PE2/PE2* or SaKKHPE2* | DS-5P-CR5_UMI-deletion | CTACACGACGCTCTTCCGATCTNNWNNVHBGT CTCTCCCAGGAATCATCTTTACCAG |
| Figure 2D~2E | | | 3p_DS_constant | AGACGTGTGCTCTTCCGAT |
| Figure 2D~2E | | | 5p-DS_CCR5 | ctacacgacgctcttccgatctGTCTCTCCCA GGAATCATCTTTACCAG |
| Figure 2D~2E | | | 3p-DS_CCR5 | agacgtgtgctcttccgatctCGACACCGAAG CAGAGTTTTTAGGAT |
| Figure 3A~3D | Serpina1 G to A or Serpina1 A to G | PE2/PE2* or SaKKHPE2* | 5p-AAT-DS | ctacacgacgctcttccgatctATAAGGCTGT GCTGACCATCG |
| Figure 3A~3D | | | | |
| Figure 3A~3D | | | | |
| Figure 3A~3D | | | 3p-AAT-DS | agacgtgtgctcttccgatctGGGGAGACTTG GTATTTTGTTCA |
| Figure 3A~3D | | | | |
| Figure 3A~3D | | | | |
| Figure 4 | Ctnnb1 TCC deletion | PE2/PE2* | ctnnb1-Uditas-FWD | GTGACTGGAGTTCAGACGTGTGCTCTTCCGAT CTGCTTCTTCAGGTAGCATTTTCAGTTCAC |
| Figure 4 | | | ctnnb1-Uditas-REV | GTGACTGGAGTTCAGACGTGTGCTCTTCCGAT CTGCTTCCAAACACAAATGCTTTACCAG |
| Figure 5 | Serpina1 A to G | AAV | SerpinA1-Uditas-FWD | GTGACTGGAGTTCAGACGTGTGCTCTTCCGAT CTagccttacaacgtgtctctgcttc |
| Figure 5 | | | SerpinA1-Uditas-REV | GTGACTGGAGTTCAGACGTGTGCTCTTCCGAT CTgcagttatttttgggtgggattca |

**Supplementary Table 4.** Potential genetic engineered liver cancer models using prime editors.

| Strain | Target gene | Accompanying gene/mutation |
|---|---|---|
| FVB/B6 | p53 R270H | c-Myc |
| FVB/B6 | HrasG12V | c-Myc |
| FVB/B6 | NrasG12V | c-Myc |
| FVB/B6 | KrasG12V | p53 R270H |
| FVB/B6 | Ctnnb1 S45F | YapS127A |
| FVB/B6 | NrasG12V | Akt |
| FVB/B6 | Ctnnb1 S45F | Akt |
| FVB/B6 | Ctnnb1 S45F | c-Met |

**Supplementary Table 5.** In vivo on target analysis at SERPINA1 site by UdiTaS.

| Figure | Sample | total_reads | | precise_editing | | small_indels(<20bp)_or_substitutions | | Deletions between pegRNA and nicking sgRNA (<100bp) | | Deletion (large_deletions > 100bp) | | AAV_insertion | |
|---|---|---|---|---|---|---|---|---|---|---|---|---|---|
| Figure 5D and 5E | | total_# | Unique_UMI# | total_# | Unique_UMI# | total_# | Unique_UMI# | total_# | Unique_UMI# | total_# | Unique_UMI# | total_# | Unique_UMI# |
| Figure 5D and 5E | Neg-R1 | 1320768 | 46632 | 96 | 14 | 377 | 29 | 54 | 7 | 9 | 1 | 0 | 0 |
| Figure 5D and 5E | Neg-R2 | 1344784 | 71231 | 112 | 18 | 385 | 52 | 44 | 5 | 14 | 2 | 0 | 0 |

| | | | | | | | | | | | | | |
|---|---|---|---|---|---|---|---|---|---|---|---|---|---|
| Figure 5D and 5E | Neg-R3 | 1127485 | 51684 | 56 | 5 | 512 | 38 | 19 | 3 | 3 | 1 | 0 | 0 |
| Figure 5D and 5E | 6wks-R1 | 1817081 | 77854 | 19854 | 924 | 4352 | 211 | 1322 | 71 | 79 | 13 | 22 | 3 |
| Figure 5D and 5E | 6wks-R2 | 1945421 | 81256 | 17526 | 865 | 2464 | 238 | 1165 | 62 | 85 | 11 | 19 | 3 |
| Figure 5D and 5E | 6wks-R3 | 1521478 | 65424 | 11548 | 614 | 3716 | 189 | 1327 | 55 | 82 | 10 | 5 | 1 |
| Figure 5D and 5E | 10wks-R1 | 1325475 | 63367 | 25513 | 1256 | 7785 | 419 | 1985 | 112 | 124 | 17 | 61 | 7 |
| Figure 5D and 5E | 10wks-R2 | 1854788 | 74541 | 27452 | 1274 | 5562 | 298 | 1428 | 86 | 75 | 21 | 32 | 6 |
| Figure 5D and 5E | 10wks-R3 | 1654876 | 78458 | 36525 | 2115 | 12235 | 712 | 2565 | 177 | 228 | 54 | 216 | 18 |

**Supplementary Sequences 1.** Sequences of reporter cell line used in this study

**mCherry Reporter sequence for A-to-G transition :**

PAM-sgRNA stop_codon

ATGGTGAGCAAGGGCGAGGAGGACAACATGGCCATCATCAAGGAGTTCATGCGCTTCAAGGTGCACATGG
AGGGCTCCGTGAACGGCCACGAGTTCGAGATCGAGGGtGAGGGtGAGGGCCGaCCCTACGAGGGCACCtA
GACCGCCAAGCTGAAGGTGACCAAGGGCGGaCCCCTGCCCTTCGCCTGGGACATCCTGTCCCCTCAGTTC
ATGTACGGCTCCAAGGCCTACGTGAAGCACCCCGCCGACATCCCCGACTACTTGAAGCTGTCCTTCCCCG
AGGGCTTCAAGTGGGAGCGCGTGATGAACTTCGAGGACGGCGGCGTGGTGACCGTGACCCAGGACTCCTC
CCTGCAGGACGGCGAGTTCATCTACAAGGTGAAGCTGCGCGGCACCAACTTCCCCTCCGACGGCCCCGTA
ATGCAGAAGAAGACCATGGGCTGGGAGGCCTCCTCCGAGCGGATGTACCCCGAGGACGGCGCCCTGAAGG
GCGAGATCAAGCAGAGGCTGAAGCTGAAGGACGGCGGCCACTACGACGCCGAGGTCAAGACCACCTACAA
GGCCAAGAAGCCCGTGCAGCTGCCCGGCGCCTACAACGTCAACATCAAGCTGGACATCACCTCCCACAAC
GAGGACTACACCATCGTGGAACAGTACGAGCGCGCCGAGGGCCGCCACTCCACCGGCGGCATGGACGAGC
TGTACAAGTAA

**Traffic Light Reporter sequence for 47bp deletion:**

GFP-47bp_Insertion-P2A-mCherry

ATGCCCGCCATGAAGATCGAGTGCCGCATCACCGGCACCCTGAACGGCGTGGAGTTCGAGCTGGTGGGCG
GCGGAGAGGGCACCCCcgaggggccccaaatcacctgcctcgtgggagacaaaatacccgttcCGAGCAG
GGtCGCATGACCAACAAGATGAAGAGCACCAAAGGCGCCCTGACCTTCAGCCCCTACCTGCTGAGCCACG
TGATGGGCTACGGCTTCTACCACTTCGGCACCTACCCCAGCGGCTACGAGAACCCCTTCCTGCACGCCAT
CAACAACGGCGGCTACACCAACACCCGCATCGAGAAGTACGAGGACGGCGGCGTGCTGCACGTGAGCTTC
AGCTACCGCTACGAGGCCGGCCGCGTGATCGGCGACTTCAAGGTGGTGGGCACCGGCTTCCCCGAGGACA
GCGTGATCTTCACCGACAAGATCATCCGCAGCAACGCCACCGTGGAGCACCTGCACCCCATGGGCGAcAA
CGTGCTGGTGGGCAGCTTCGCCCGCACCTTCAGCCTGCGCGACGGCGGCTACTACAGCTTCGTGGTGGAC
AGCCACATGCACTTCAAGAGCGCCATCCACCCCAGCATCCTGCAGAACGGGGGCCCCATGTTCGCCTTCC

GCCGCGTGGAGGAGCTGCACAGCAACACCGAGCTGGGCATCGTGGAGTACCAGCACGCCTTCAAGACCCC
CATCGCCTTCGCCAGATCTCGAGCTCGAggccacgaatttctcgctactcaagcaggccggcgatgtcg
aggaaaaccctggtcctGTGAGCAAGGGCGAGGAGGACAACATGGCCATCATCAAGGAGTTCATGCGCTT
CAAGGTGCACATGGAGGGCTCCGTGAACGGCCACGAGTTCGAGATCGAGGGCGAGGGCGAGGGCCGCCCC
TACGAGGGCACCCAGACCGCCAAGCTGAAGGTGACCAAGGGCGGCCCCCTGCCCTTCGCCTGGGACATCC
TGTCCCCTCAGTTCATGTACGGCTCCAAGGCCTACGTGAAGCACCCCGCCGACATCCCCGACTACTTGAA
GCTGTCCTTCCCCGAGGGCTTCAAGTGGGAGCGCGTGATGAACTTCGAGGACGGCGGCGTGGTGACCGTG
ACCCAGGACTCCTCCCTGCAGGACGGCGAGTTCATCTACAAGGTGAAGCTGCGCGGCACCAACTTCCCCT
CCGACGGCCCCGTAATGCAGAAGAAGACCATGGGCTGGGAGGCCTCCTCCGAGCGGATGTACCCCGAGGA
CGGCGCCCTGAAGGGCGAGATCAAGCAGAGGCTGAAGCTGAAGGACGGCGGCCACTACGACGCCGAGGTC
AAGACCACCTACAAGGCCAAGAAGCCCGTGCAGCTGCCCGGCGCCTACAACGTCAACATCAAGCTGGACA
TCACCTCCCACAACGAGGACTACACCATCGTGGAACAGTACGAGCGCGCCGAGGGCCGCCACTCCACCGG
CGGCATGGACGAGCTGTACAAGTAATAG

**Traffic Light Reporter sequence for 39bp deletion + 18bp insertion:**

GFP-39bp_Insertion-T2A-mCherry

atggtgagcaagggcgaggagctgttcaccggggtggtgcccatcctggtcgagctggacggcgacgtaa
acggccacaagttcagcgtgtccggcTTTGGCGAGACAAATCACCTGCCTCGTGGAATACGGTAAaccta
cggcaagctgaccctgaagttcatctgcaccaccggcaagctgcccgtgccctggcccaccctcgtgacc
accctgacctacggcgtgcagtgcttcagccgctaccccgaccacatgaagcagcacgacttcttcaagt
ccgccatgcccgaaggctacgtccaggagcgcaccatcttcttcaaggacgacggcaactacaagacccg
cgccgaggtgaagttcgagggcgacaccctggtgaaccgcatcgagctgaagggcatcgacttcaaggag
gacggcaacatcctggggcacaagctggagtacaactacaacagccacaacgtctatatcatggccgaca
agcagaagaacggcatcaaggtgaacttcaagatccgccacaacatcgaggacggcagcgtgcagctcgc
cgaccactaccagcagaacacccccatcggcgacggccccgtgctgctgcccgacaaccactacctgagc
acccagtccgccctgagcaaagaccccaacgagaagcgcgatcacatggtcctgctggagttcgtgaccg
ccgccgggatcactctcggcatggacgagctgtacaagtaactggatccggtgagggcagaggaagtctt
ctaacatgcggtgacgtggaggagaatccgggccctgtgagcaagggcgaggaggataactccgccatca
tcaaggagttcctgcgcttcaaggtgcacatggagggctccgtgaacggccacgagttcgagatcgaggg
cgagggcgagggccgcccctacgagggcacccagaccgccaagctgaaggtgaccaagggtggcccccctg
cccttcgcctgggacatcctgtcccctcagttcatgtacggctccaaggcctacgtgaagcaccccgccg
acatccccgactacttgaagctgtccttccccgagggcttcaagtgggagcgcgtgatgaacttcgagga
cggcggcgtggtgaccgtgacccaggactcctctctgcaggacggcgagttcatctacaaggtgaagctg
cgcggcaccaacttcccctccgacggccccgtaatgcagaagaagaccatgggctgggaggcctcctccg
agcggatgtaccccgaggacggcgccctgaagggcgagatcaagcagaggctgaagctgaaggacggcgg
ccactacgacgctgaggtcaagaccacctacaaggccaagaagcccgtgcagctgcccggcgcctacaac
gtcaacatcaagttggacatcacctcccacaacgaggactacaccatcgtggaacagtacgaacgcgccg
agggccgccactccaccggcggcatggacgagctgtacaagtga

**Supplementary Sequences 2.** Sequence of backbone plasmid used for pegRNA and nicking sgRNA cloning

Backbone of pegRNA used for PE2 or PE2*: U6 promoter + spCas9-sgRNA scaffold

GTGGCACTTTTCGGGGAAATGTGCGCGGAACCCCTATTTGTTTATTTTTCTAAATACATTCAAA
TATGTATCCGCTCATGAGACAATAACCCTGATAAATGCTTCAATAATATTGAAAAAGGAAGAGT
ATGAGTATTCAACATTTCCGTGTCGCCCTTATTCCCTTTTTTGCGGCATTTTGCCTTCCTGTTT
TTGCTCACCCAGAAACGCTGGTGAAAGTAAAAGATGCTGAAGATCAGTTGGGTGCACGAGTGGG
TTACATCGAACTGGATCTCAACAGCGGTAAGATCCTTGAGAGTTTTCGCCCCGAAGAACGTTTT
CCAATGATGAGCACTTTTAAAGTTCTGCTATGTGGCGCGGTATTATCCCGTATTGACGCCGGGC
AAGAGCAACTCGGTCGCCGCATACACTATTCTCAGAATGACTTGGTTGAGTACTCACCAGTCAC
AGAAAAGCATCTTACGGATGGCATGACAGTAAGAGAATTATGCAGTGCTGCCATAACCATGAGT
GATAACACTGCGGCCAACTTACTTCTGACAACGATCGGAGGACCGAAGGAGCTAACCGCTTTTT
TGCACAACATGGGGGATCATGTAACTCGCCTTGATCGTTGGGAACCGGAGCTGAATGAAGCCAT
ACCAAACGACGAGCGTGACACCACGATGCCTGTAGCAATGGCAACAACGTTGCGCAAACTATTA
ACTGGCGAACTACTTACTCTAGCTTCCCGGCAACAATTAATAGACTGGATGGAGGCGGATAAAG
TTGCAGGACCACTTCTGCGCTCGGCCCTTCCGGCTGGCTGGTTTATTGCTGATAAATCTGGAGC
CGGTGAGCGTGGGTCTCGCGGTATCATTGCAGCACTGGGGCCAGATGGTAAGCCCTCCCGTATC
GTAGTTATCTACACGACGGGGAGTCAGGCAACTATGGATGAACGAAATAGACAGATCGCTGAGA
TAGGTGCCTCACTGATTAAGCATTGGTAACTGTCAGACCAAGTTTACTCATATATACTTTAGAT
TGATTTAAAACTTCATTTTTAATTTAAAAGGATCTAGGTGAAGATCCTTTTTGATAATCTCATG
ACCAAAATCCCTTAACGTGAGTTTTCGTTCCACTGAGCGTCAGACCCCGTAGAAAAGATCAAAG
GATCTTCTTGAGATCCTTTTTTTCTGCGCGTAATCTGCTGCTTGCAAACAAAAAAACCACCGCT
ACCAGCGGTGGTTTGTTTGCCGGATCAAGAGCTACCAACTCTTTTTCCGAAGGTAACTGGCTTC
AGCAGAGCGCAGATACCAAATACTGTCCTTCTAGTGTAGCCGTAGTTAGGCCACCACTTCAAGA
ACTCTGTAGCACCGCCTACATACCTCGCTCTGCTAATCCTGTTACCAGTGGCTGCTGCCAGTGG
CGATAAGTCGTGTCTTACCGGGTTGGACTCAAGACGATAGTTACCGGATAAGGCGCAGCGGTCG
GGCTGAACGGGGGGTTCGTGCACACAGCCCAGCTTGGAGCGAACGACCTACACCGAACTGAGAT
ACCTACAGCGTGAGCTATGAGAAAGCGCCACGCTTCCCGAAGGGAGAAAGGCGGACAGGTATCC
GGTAAGCGGCAGGGTCGGAACAGGAGAGCGCACGAGGGAGCTTCCAGGGGGAAACGCCTGGTAT
CTTTATAGTCCTGTCGGGTTTCGCCACCTCTGACTTGAGCGTCGATTTTTGTGATGCTCGTCAG
GGGGGCGGAGCCTATGGAAAAACGCCAGCAACGCGGCCTTTTTACGGTTCCTGGCCTTTTGCTG
GCCTTTTGCTCACATGTTCTTTCCTGCGTTATCCCCTGATTCTGTGGATAACCGTATTACCGCC
TTTGAGTGAGCTGATACCGCTCGCCGCAGCCGAACGACCGAGCGCAGCGAGTCAGTGAGCGAGG
AAGCGGAAGAGCGCCCAATACGCAAACCGCCTCTCCCCGCGCGTTGGCCGATTCATTAATGCAG
CTGGCACGACAGGTTTCCCGACTGGAAAGCGGGCAGTGAGCGCAACGCAATTAATGTGAGTTAG
CTCACTCATTAGGCACCCCAGGCTTTACACTTTATGCTTCCGGCTCGTATGTTGTGTGGAATTG
TGAGCGGATAACAATTTCACACAGGAAACAGCTATGACCATGATTACGCCAAGCGCGCAATTAA
CCCTCACTAAAGGGAACAAAAGCTGGAGCTCCACCGCGGTGGCGGCCGCCCCCTTCACCGAGGG
CCTATTTCCCATGATTCCTTCATATTTGCATATACGATACAAGGCTGTTAGAGAGATAATTGGA
ATTAATTTGACTGTAAACACAAAGATATTAGTACAAAATACGTGACGTAGAAAGTAATAATTTC
TTGGGTAGTTTGCAGTTTTAAAATTATGTTTTAAAATGGACTATCATATGCTTACCGTAACTTG
AAAGTATTTCGATTTCTTGGCTTTATATATCTTGTGGAAAGGACGAAACACCGGTGTGCAGGTG
AGTGATCCAAACGCCCGGCGGCAACCGAGCGTTCTGAACAAATCCAGATGGAGTTCTGAGGTCA
TTACTGGATCTATCAACAGGAGTCCAAGCGAGCTCTCGAACCCCAGAGTCCCGCTCAGAAGAAC
TCGTCAAGAAGGCGATAGAAGGCGATGCGCTGCGAATCGGGAGCGGCGATACCGTAAAGCACGA
GGAAGCGGTCAGCCCATTCGCCGCCAAGCTCTTCAGCAATATCACGGGTAGCCAACGCTATGTC
CTGATAGCGGTCCGCCACACCCAGCCGGCCACAGTCGATGAATCCAGAAAAGCGGCCATTTTCC
ACCATGATATTCGGCAAGCAGGCATCGCCATGGGTCACGACGAGATCCTCGCCGTCGGGCATGC

GCGCCTTGAGCCTGGCGAACAGTTCGGCTGGCGCGAGCCCCTGATGCTCTTCGTCCAGATCATC
CTGATCGACAAGACCGGCTTCCATCCGAGTACGTGCTCGCTCGATGCGATGTTTCGCTTGGTGG
TCGAATGGGCAGGTAGCCGGATCAAGCGTATGCAGCCGCCGCATTGCATCAGCCATGATGGATA
CTTTCTCGGCAGGAGCAAGGTGAGATGACAGGAGATCCTGCCCCGGCACTTCGCCCAATAGCAG
CCAGTCCCTTCCCGCTTCAGTGACAACGTCGAGCACAGCTGCGCAAGGAACGCCCGTCGTGGCC
AGCCACGATAGCCGCGCTGCCTCGTCCTGCAGTTCATTCAGGGCACCGGACAGGTCGGTCTTGA
CAAAAAGAACCGGGCGCCCCTGCGCTGACAGCCGGAACACGGCGGCATCAGAGCAGCCGATTGT
CTGTTGTGCCCAGTCATAGCCGAATAGCCTCTCCACCCAAGCGGCCGGAGAACCTGCGTGCAAT
CCATCTTGTTCAATCATGCGAAACGATCCTCATCCTGTCTCTTGATCAGATCTTGATCCCCTGC
GCCATCAGATCCTTGGCGGCAAGAAAGCCATCCAGTTTACTTTGCAGGGCTTCCCAACCTTACC
AGAGGGCGCCCCAGCTGGCAATTCCGACGGATCAAGAACCTGCTGACGTTTTAGAGCTAGAAATAG
CAAGTTAAAATAAGGCTAGTCCGTTATCAACTTGAAAAAGTGGGACCGAGTCGGTCCTTTTTTTGAATT
CGATATCAAGCTTATCGATACCGTCGACCTCGAGGGGGGGCCCGGTACCCAATTCGCCCTATAG
TGAGTCGTATTACGCGCGCTCACTGGCCGTCGTTTTACAACGTCGTGACTGGGAAAACCCTGGC
GTTACCCAACTTAATCGCCTTGCAGCACATCCCCCTTTCGCCAGCTGGCGTAATAGCGAAGAGG
CCCGCACCGATCGCCCTTCCCAACAGTTGCGCAGCCTGAATGGCGAATGGGACGCGCCCTGTAG
CGGCGCATTAAGCGCGGCGGGTGTGGTGGTTACGCGCAGCGTGACCGCTACACTTGCCAGCGCC
CTAGCGCCCGCTCCTTTCGCTTTCTTCCCTTCCTTTCTCGCCACGTTCGCCGGCTTTCCCCGTC
AAGCTCTAAATCGGGGGCTCCCTTTAGGGTTCCGATTTAGTGCTTTACGGCACCTCGACCCCAA
AAAACTTGATTAGGGTGATGGTTCACGTAGTGGGCCATCGCCCTGATAGACGGTTTTTCGCCCT
TTGACGTTGGAGTCCACGTTCTTTAATAGTGGACTCTTGTTCCAAACTGGAACAACACTCAACC
CTATCTCGGTCTATTCTTTTGATTTATAAGGGATTTTGCCGATTTCGGCCTATTGGTTAAAAAA
TGAGCTGATTTAACAAAAATTTAACGCGAATTTTAACAAAATATTAACGCTTACAATTTAG

Backbone of Nicking sgRNA used for PE2 or PE2*: U6 promoter + spCas9-sgRNA scaffold

GTGGCACTTTTCGGGGAAATGTGCGCGGAACCCCTATTTGTTTATTTTTCTAAATACATTCAAATATGTA
TCCGCTCATGAGACAATAACCCTGATAAATGCTTCAATAATATTGAAAAAGGAAGAGTATGAGTATTCAA
CATTTCCGTGTCGCCCTTATTCCCTTTTTTGCGGCATTTTGCCTTCCTGTTTTTGCTCACCCAGAAACGC
TGGTGAAAGTAAAAGATGCTGAAGATCAGTTGGGTGCACGAGTGGGTTACATCGAACTGGATCTCAACAG
CGGTAAGATCCTTGAGAGTTTTCGCCCCGAAGAACGTTTTCCAATGATGAGCACTTTTAAAGTTCTGCTA
TGTGGCGCGGTATTATCCCGTATTGACGCCGGGCAAGAGCAACTCGGTCGCCGCATACACTATTCTCAGA
ATGACTTGGTTGAGTACTCACCAGTCACAGAAAAGCATCTTACGGATGGCATGACAGTAAGAGAATTATG
CAGTGCTGCCATAACCATGAGTGATAACACTGCGGCCAACTTACTTCTGACAACGATCGGAGGACCGAAG
GAGCTAACCGCTTTTTTGCACAACATGGGGGATCATGTAACTCGCCTTGATCGTTGGGAACCGGAGCTGA
ATGAAGCCATACCAAACGACGAGCGTGACACCACGATGCCTGTAGCAATGGCAACAACGTTGCGCAAACT
ATTAACTGGCGAACTACTTACTCTAGCTTCCCGGCAACAATTAATAGACTGGATGGAGGCGGATAAAGTT
GCAGGACCACTTCTGCGCTCGGCCCTTCCGGCTGGCTGGTTTATTGCTGATAAATCTGGAGCCGGTGAGC
GTGGGTCTCGCGGTATCATTGCAGCACTGGGGCCAGATGGTAAGCCCTCCCGTATCGTAGTTATCTACAC
GACGGGGAGTCAGGCAACTATGGATGAACGAAATAGACAGATCGCTGAGATAGGTGCCTCACTGATTAAG
CATTGGTAACTGTCAGACCAAGTTTACTCATATATACTTTAGATTGATTTAAAACTTCATTTTTAATTTA
AAAGGATCTAGGTGAAGATCCTTTTTGATAATCTCATGACCAAAATCCCTTAACGTGAGTTTTCGTTCCA
CTGAGCGTCAGACCCCGTAGAAAAGATCAAAGGATCTTCTTGAGATCCTTTTTTTCTGCGCGTAATCTGC
TGCTTGCAAACAAAAAAACCACCGCTACCAGCGGTGGTTTGTTTGCCGGATCAAGAGCTACCAACTCTTT
TTCCGAAGGTAACTGGCTTCAGCAGAGCGCAGATACCAAATACTGTCCTTCTAGTGTAGCCGTAGTTAGG
CCACCACTTCAAGAACTCTGTAGCACCGCCTACATACCTCGCTCTGCTAATCCTGTTACCAGTGGCTGCT
GCCAGTGGCGATAAGTCGTGTCTTACCGGGTTGGACTCAAGACGATAGTTACCGGATAAGGCGCAGCGGT

CGGGCTGAACGGGGGGTTCGTGCACACAGCCCAGCTTGGAGCGAACGACCTACACCGAACTGAGATACCT
ACAGCGTGAGCTATGAGAAAGCGCCACGCTTCCCGAAGGGAGAAAGGCGGACAGGTATCCGGTAAGCGGC
AGGGTCGGAACAGGAGAGCGCACGAGGGAGCTTCCAGGGGGAAACGCCTGGTATCTTTATAGTCCTGTCG
GGTTTCGCCACCTCTGACTTGAGCGTCGATTTTTGTGATGCTCGTCAGGGGGGCGGAGCCTATGGAAAAA
CGCCAGCAACGCGGCCTTTTTACGGTTCCTGGCCTTTTGCTGGCCTTTTGCTCACATGTTCTTTCCTGCG
TTATCCCCTGATTCTGTGGATAACCGTATTACCGCCTTTGAGTGAGCTGATACCGCTCGCCGCAGCCGAA
CGACCGAGCGCAGCGAGTCAGTGAGCGAGGAAGCGGAAGAGCGCCCAATACGCAAACCGCCTCTCCCCGC
GCGTTGGCCGATTCATTAATGCAGCTGGCACGACAGGTTTCCCGACTGGAAAGCGGGCAGTGAGCGCAAC
GCAATTAATGTGAGTTAGCTCACTCATTAGGCACCCCAGGCTTTACACTTTATGCTTCCGGCTCGTATGT
TGTGTGGAATTGTGAGCGGATAACAATTTCACACAGGAAACAGCTATGACCATGATTACGCCAAGCGCGC
AATTAACCCTCACTAAAGGGAACAAAAGCTGGAGCTCCACCGCGGTGGCGGCCGCCCCCTTCACCGAGGG
CCTATTTCCCATGATTCCTTCATATTTGCATATACGATACAAGGCTGTTAGAGAGATAATTGGAATTAAT
TTGACTGTAAACACAAAGATATTAGTACAAAATACGTGACGTAGAAAGTAATAATTTCTTGGGTAGTTTG
CAGTTTTAAAATTATGTTTTAAAATGGACTATCATATGCTTACCGTAACTTGAAAGTATTTCGATTTCTT
GGCTTTATATATCTTGTGGAAAGGACGAAACACCGGTGTGCAGGTGAGTGATCCAAACGCCCGGCGGCAA
CCGAGCGTTCTGAACAAATCCAGATGGAGTTCTGAGGTCATTACTGGATCTATCAACAGGAGTCCAAGCG
AGCTCTCGAACCCCAGAGTCCCGCTCAGAAGAACTCGTCAAGAAGGCGATAGAAGGCGATGCGCTGCGAA
TCGGGAGCGGCGATACCGTAAAGCACGAGGAAGCGGTCAGCCCATTCGCCGCCAAGCTCTTCAGCAATAT
CACGGGTAGCCAACGCTATGTCCTGATAGCGGTCCGCCACACCCAGCCGGCCACAGTCGATGAATCCAGA
AAAGCGGCCATTTTCCACCATGATATTCGGCAAGCAGGCATCGCCATGGGTCACGACGAGATCCTCGCCG
TCGGGCATGCGCGCCTTGAGCCTGGCGAACAGTTCGGCTGGCGCGAGCCCCTGATGCTCTTCGTCCAGAT
CATCCTGATCGACAAGACCGGCTTCCATCCGAGTACGTGCTCGCTCGATGCGATGTTTCGCTTGGTGGTC
GAATGGGCAGGTAGCCGGATCAAGCGTATGCAGCCGCCGCATTGCATCAGCCATGATGGATACTTTCTCG
GCAGGAGCAAGGTGAGATGACAGGAGATCCTGCCCCGGCACTTCGCCCAATAGCAGCCAGTCCCTTCCCG
CTTCAGTGACAACGTCGAGCACAGCTGCGCAAGGAACGCCCGTCGTGGCCAGCCACGATAGCCGCGCTGC
CTCGTCCTGCAGTTCATTCAGGGCACCGGACAGGTCGGTCTTGACAAAAAGAACCGGGCGCCCCTGCGCT
GACAGCCGGAACACGGCGGCATCAGAGCAGCCGATTGTCTGTTGTGCCCAGTCATAGCCGAATAGCCTCT
CCACCCAAGCGGCCGGAGAACCTGCGTGCAATCCATCTTGTTCAATCATGCGAAACGATCCTCATCCTGT
CTCTTGATCAGATCTTGATCCCCTGCGCCATCAGATCCTTGGCGGCAAGAAAGCCATCCAGTTTACTTTG
CAGGGCTTCCCAACCTTACCAGAGGGCGCCCCAGCTGGCAATTCCGACGGATCAAGAACCTGCTGACGTT
TTAGAGCTAGAAATAGCAAGTTAAAATAAGGCTAGTCCGTTATCAACTTGAAAAAGTGGCACCGAGTCGG
TGCTTTTTTTGAATTCGATATCAAGCTTATCGATACCGTCGACCTCGAGGGGGGGCCCGGTACCCAATTC
GCCCTATAGTGAGTCGTATTACGCGCGCTCACTGGCCGTCGTTTTACAACGTCGTGACTGGGAAAACCCT
GGCGTTACCCAACTTAATCGCCTTGCAGCACATCCCCCTTTCGCCAGCTGGCGTAATAGCGAAGAGGCCC
GCACCGATCGCCCTTCCCAACAGTTGCGCAGCCTGAATGGCGAATGGGACGCGCCCTGTAGCGGCGCATT
AAGCGCGGCGGGTGTGGTGGTTACGCGCAGCGTGACCGCTACACTTGCCAGCGCCCTAGCGCCCGCTCCT
TTCGCTTTCTTCCCTTCCTTTCTCGCCACGTTCGCCGGCTTTCCCCGTCAAGCTCTAAATCGGGGGCTCC
CTTTAGGGTTCCGATTTAGTGCTTTACGGCACCTCGACCCCAAAAAACTTGATTAGGGTGATGGTTCACG
TAGTGGGCCATCGCCCTGATAGACGGTTTTTCGCCCTTTGACGTTGGAGTCCACGTTCTTTAATAGTGGA
CTCTTGTTCCAAACTGGAACAACACTCAACCCTATCTCGGTCTATTCTTTTGATTTATAAGGATTTTGC
CGATTTCGGCCTATTGGTTAAAAAATGAGCTGATTTAACAAAAATTTAACGCGAATTTTAACAAAATATT
AACGCTTACAATTTAG

Backbone of pegRNA or Nicking sgRNA used for for SaPE2* and SaPE2*: U6 promoter + SaCas9 sgRNA scaffold

GTGGCACTTTTCGGGGAAATGTGCGCGGAACCCCTATTTGTTTATTTTTCTAAATACATTCAAATATGTA
TCCGCTCATGAGACAATAACCCTGATAAATGCTTCAATAATATTGAAAAAGGAAGAGTATGAGTATTCAA
CATTTCCGTGTCGCCCTTATTCCCTTTTTTGCGGCATTTTGCCTTCCTGTTTTTGCTCACCCAGAAACGC
TGGTGAAAGTAAAAGATGCTGAAGATCAGTTGGGTGCACGAGTGGGTTACATCGAACTGGATCTCAACAG
CGGTAAGATCCTTGAGAGTTTTCGCCCCGAAGAACGTTTTCCAATGATGAGCACTTTTAAAGTTCTGCTA
TGTGGCGCGGTATTATCCCGTATTGACGCCGGGCAAGAGCAACTCGGTCGCCGCATACACTATTCTCAGA
ATGACTTGGTTGAGTACTCACCAGTCACAGAAAAGCATCTTACGGATGGCATGACAGTAAGAGAATTATG
CAGTGCTGCCATAACCATGAGTGATAACACTGCGGCCAACTTACTTCTGACAACGATCGGAGGACCGAAG
GAGCTAACCGCTTTTTTGCACAACATGGGGGATCATGTAACTCGCCTTGATCGTTGGGAACCGGAGCTGA
ATGAAGCCATACCAAACGACGAGCGTGACACCACGATGCCTGTAGCAATGGCAACAACGTTGCGCAAACT
ATTAACTGGCGAACTACTTACTCTAGCTTCCCGGCAACAATTAATAGACTGGATGGAGGCGGATAAAGTT
GCAGGACCACTTCTGCGCTCGGCCCTTCCGGCTGGCTGGTTTATTGCTGATAAATCTGGAGCCGGTGAGC
GTGGGTCTCGCGGTATCATTGCAGCACTGGGGCCAGATGGTAAGCCCTCCCGTATCGTAGTTATCTACAC
GACGGGGAGTCAGGCAACTATGGATGAACGAAATAGACAGATCGCTGAGATAGGTGCCTCACTGATTAAG
CATTGGTAACTGTCAGACCAAGTTTACTCATATATACTTTAGATTGATTTAAAACTTCATTTTTAATTTA
AAAGGATCTAGGTGAAGATCCTTTTTGATAATCTCATGACCAAAATCCCTTAACGTGAGTTTTCGTTCCA
CTGAGCGTCAGACCCCGTAGAAAAGATCAAAGGATCTTCTTGAGATCCTTTTTTTCTGCGCGTAATCTGC
TGCTTGCAAACAAAAAAACCACCGCTACCAGCGGTGGTTTGTTTGCCGGATCAAGAGCTACCAACTCTTT
TTCCGAAGGTAACTGGCTTCAGCAGAGCGCAGATACCAAATACTGTCCTTCTAGTGTAGCCGTAGTTAGG
CCACCACTTCAAGAACTCTGTAGCACCGCCTACATACCTCGCTCTGCTAATCCTGTTACCAGTGGCTGCT
GCCAGTGGCGATAAGTCGTGTCTTACCGGGTTGGACTCAAGACGATAGTTACCGGATAAGGCGCAGCGGT
CGGGCTGAACGGGGGGTTCGTGCACACAGCCCAGCTTGGAGCGAACGACCTACACCGAACTGAGATACCT
ACAGCGTGAGCTATGAGAAAGCGCCACGCTTCCCGAAGGGAGAAAGGCGGACAGGTATCCGGTAAGCGGC
AGGGTCGGAACAGGAGAGCGCACGAGGGAGCTTCCAGGGGGAAACGCCTGGTATCTTTATAGTCCTGTCG
GGTTTCGCCACCTCTGACTTGAGCGTCGATTTTTGTGATGCTCGTCAGGGGGGCGGAGCCTATGGAAAAA
CGCCAGCAACGCGGCCTTTTTACGGTTCCTGGCCTTTTGCTGGCCTTTTGCTCACATGTTCTTTCCTGCG
TTATCCCCTGATTCTGTGGATAACCGTATTACCGCCTTTGAGTGAGCTGATACCGCTCGCCGCAGCCGAA
CGACCGAGCGCAGCGAGTCAGTGAGCGAGGAAGCGGAAGAGCGCCCAATACGCAAACCGCCTCTCCCCGC
GCGTTGGCCGATTCATTAATGCAGCTGGCACGACAGGTTTCCCGACTGGAAAGCGGGCAGTGAGCGCAAC
GCAATTAATGTGAGTTAGCTCACTCATTAGGCACCCCAGGCTTTACACTTTATGCTTCCGGCTCGTATGT
TGTGTGGAATTGTGAGCGGATAACAATTTCACACAGGAAACAGCTATGACCATGATTACGCCAAGCGCGC
AATTAACCCTCACTAAAGGGAACAAAAGCTGGAGCTCCACCGCGGTGGCGGCCGCCCCCTTCACCGAGGG
CCTATTTCCCATGATTCCTTCATATTTGCATATACGATACAAGGCTGTTAGAGAGATAATTGGAATTAAT
TTGACTGTAAACACAAAGATATTAGTACAAAATACGTGACGTAGAAAGTAATAATTTCTTGGGTAGTTTG
CAGTTTTAAAATTATGTTTTAAAATGGACTATCATATGCTTACCGTAACTTGAAAGTATTTCGATTTCTT
GGCTTTATATATCTTGTGGAAAGGACGAAACACCGGTGTGCAGGTGAGTGATCCAAACGCCCGGCGGCAA
CCGAGCGTTCTGAACAAATCCAGATGGAGTTCTGAGGTCATTACTGGATCTATCAACAGGAGTCCAAGCG
AGCTCTCGAACCCCAGAGTCCCGCTCAGAAGAACTCGTCAAGAAGGCGATAGAAGGCGATGCGCTGCGAA
TCGGGAGCGGCGATACCGTAAAGCACGAGGAAGCGGTCAGCCCATTCGCCGCCAAGCTCTTCAGCAATAT
CACGGGTAGCCAACGCTATGTCCTGATAGCGGTCCGCCACACCCAGCCGGCCACAGTCGATGAATCCAGA
AAAGCGGCCATTTTCCACCATGATATTCGGCAAGCAGGCATCGCCATGGGTCACGACGAGATCCTCGCCG
TCGGGCATGCGCGCCTTGAGCCTGGCGAACAGTTCGGCTGGCGCGAGCCCCTGATGCTCTTCGTCCAGAT
CATCCTGATCGACAAGACCGGCTTCCATCCGAGTACGTGCTCGCTCGATGCGATGTTTCGCTTGGTGGTC
GAATGGGCAGGTAGCCGGATCAAGCGTATGCAGCCGCCGCATTGCATCAGCCATGATGGATACTTTCTCG
GCAGGAGCAAGGTGAGATGACAGGAGATCCTGCCCCGGCACTTCGCCCAATAGCAGCCAGTCCCTTCCCG
CTTCAGTGACAACGTCGAGCACAGCTGCGCAAGGAACGCCCGTCGTGGCCAGCCACGATAGCCGCGCTGC

CTCGTCCTGCAGTTCATTCAGGGCACCGGACAGGTCGGTCTTGACAAAAAGAACCGGGCGCCCCTGCGCT
GACAGCCGGAACACGGCGGCATCAGAGCAGCCGATTGTCTGTTGTGCCCAGTCATAGCCGAATAGCCTCT
CCACCCAAGCGGCCGGAGAACCTGCGTGCAATCCATCTTGTTCAATCATGCGAAACGATCCTCATCCTGT
CTCTTGATCAGATCTTGATCCCCTGCGCCATCAGATCCTTGGCGGCAAGAAAGCCATCCAGTTTACTTTG
CAGGGCTTCCCAACCTTACCAGAGGGCGCCCCAGCTGGCAATTCCGACGGATCAgctagcAGAACCTGCT
GACGTTTTAGTACTCTGGAAACAGAATCTACTAAAACAAGGCAAAATGCCGTGTTTATCTCGTCAACTTG
TTGGCGAGATTTTTTTGAATTCGATATCAAGCTTATCGATACCGTCGACCTCGAGGGGGGGCCCGGTACC
CAATTCGCCCTATAGTGAGTCGTATTACGCGCGCTCACTGGCCGTCGTTTTACAACGTCGTGACTGGGAA
AACCCTGGCGTTACCCAACTTAATCGCCTTGCAGCACATCCCCCTTTCGCCAGCTGGCGTAATAGCGAAG
AGGCCCGCACCGATCGCCCTTCCCAACAGTTGCGCAGCCTGAATGGCGAATGGGACGCGCCCTGTAGCGG
CGCATTAAGCGCGGCGGGTGTGGTGGTTACGCGCAGCGTGACCGCTACACTTGCCAGCGCCCTAGCGCCC
GCTCCTTTCGCTTTCTTCCCTTCCTTTCTCGCCACGTTCGCCGGCTTTCCCCGTCAAGCTCTAAATCGGG
GGCTCCCTTTAGGGTTCCGATTTAGTGCTTTACGGCACCTCGACCCCAAAAAACTTGATTAGGGTGATGG
TTCACGTAGTGGGCCATCGCCCTGATAGACGGTTTTTCGCCCTTTGACGTTGGAGTCCACGTTCTTTAAT
AGTGGACTCTTGTTCCAAACTGGAACAACACTCAACCCTATCTCGGTCTATTCTTTTGATTTATAAGGGA
TTTTGCCGATTTCGGCCTATTGGTTAAAAAATGAGCTGATTTAACAAAAATTTAACGCGAATTTTAACAA
AATATTAACGCTTACAATTTAG

PegRNA used for for SaPE2* : U6 promoter-sgRNA-Scaffold-RT_template-PBS (CCR5 pegRNA as an example)

GTGGCACTTTTCGGGGAAATGTGCGCGGAACCCCTATTTGTTTATTTTTCTAAATACATTCAAATATGTA
TCCGCTCATGAGACAATAACCCTGATAAATGCTTCAATAATATTGAAAAAGGAAGAGTATGAGTATTCAA
CATTTCCGTGTCGCCCTTATTCCCTTTTTTGCGGCATTTTGCCTTCCTGTTTTTGCTCACCCAGAAACGC
TGGTGAAAGTAAAAGATGCTGAAGATCAGTTGGGTGCACGAGTGGGTTACATCGAACTGGATCTCAACAG
CGGTAAGATCCTTGAGAGTTTTCGCCCCGAAGAACGTTTTCCAATGATGAGCACTTTTAAAGTTCTGCTA
TGTGGCGCGGTATTATCCCGTATTGACGCCGGGCAAGAGCAACTCGGTCGCCGCATACACTATTCTCAGA
ATGACTTGGTTGAGTACTCACCAGTCACAGAAAAGCATCTTACGGATGGCATGACAGTAAGAGAATTATG
CAGTGCTGCCATAACCATGAGTGATAACACTGCGGCCAACTTACTTCTGACAACGATCGGAGGACCGAAG
GAGCTAACCGCTTTTTTGCACAACATGGGGGATCATGTAACTCGCCTTGATCGTTGGGAACCGGAGCTGA
ATGAAGCCATACCAAACGACGAGCGTGACACCACGATGCCTGTAGCAATGGCAACAACGTTGCGCAAACT
ATTAACTGGCGAACTACTTACTCTAGCTTCCCGGCAACAATTAATAGACTGGATGGAGGCGGATAAAGTT
GCAGGACCACTTCTGCGCTCGGCCCTTCCGGCTGGCTGGTTTATTGCTGATAAATCTGGAGCCGGTGAGC
GTGGGTCTCGCGGTATCATTGCAGCACTGGGGCCAGATGGTAAGCCCTCCCGTATCGTAGTTATCTACAC
GACGGGGAGTCAGGCAACTATGGATGAACGAAATAGACAGATCGCTGAGATAGGTGCCTCACTGATTAAG
CATTGGTAACTGTCAGACCAAGTTTACTCATATATACTTTAGATTGATTTAAAACTTCATTTTTAATTTA
AAAGGATCTAGGTGAAGATCCTTTTTGATAATCTCATGACCAAAATCCCTTAACGTGAGTTTTCGTTCCA
CTGAGCGTCAGACCCCGTAGAAAAGATCAAAGGATCTTCTTGAGATCCTTTTTTTCTGCGCGTAATCTGC
TGCTTGCAAACAAAAAAACCACCGCTACCAGCGGTGGTTTGTTTGCCGGATCAAGAGCTACCAACTCTTT
TTCCGAAGGTAACTGGCTTCAGCAGAGCGCAGATACCAAATACTGTCCTTCTAGTGTAGCCGTAGTTAGG
CCACCACTTCAAGAACTCTGTAGCACCGCCTACATACCTCGCTCTGCTAATCCTGTTACCAGTGGCTGCT
GCCAGTGGCGATAAGTCGTGTCTTACCGGGTTGGACTCAAGACGATAGTTACCGGATAAGGCGCAGCGGT
CGGGCTGAACGGGGGGTTCGTGCACACAGCCCAGCTTGGAGCGAACGACCTACACCGAACTGAGATACCT
ACAGCGTGAGCTATGAGAAAGCGCCACGCTTCCCGAAGGGAGAAAGGCGGACAGGTATCCGGTAAGCGGC
AGGGTCGGAACAGGAGAGCGCACGAGGGAGCTTCCAGGGGGAAACGCCTGGTATCTTTATAGTCCTGTCG
GGTTTCGCCACCTCTGACTTGAGCGTCGATTTTTGTGATGCTCGTCAGGGGGGCGGAGCCTATGGAAAAA

CGCCAGCAACGCGGCCTTTTTACGGTTCCTGGCCTTTTGCTGGCCTTTTGCTCACATGTTCTTTCCTGCG
TTATCCCCTGATTCTGTGGATAACCGTATTACCGCCTTTGAGTGAGCTGATACCGCTCGCCGCAGCCGAA
CGACCGAGCGCAGCGAGTCAGTGAGCGAGGAAGCGGAAGAGCGCCCAATACGCAAACCGCCTCTCCCCGC
GCGTTGGCCGATTCATTAATGCAGCTGGCACGACAGGTTTCCCGACTGGAAAGCGGGCAGTGAGCGCAAC
GCAATTAATGTGAGTTAGCTCACTCATTAGGCACCCCAGGCTTTACACTTTATGCTTCCGGCTCGTATGT
TGTGTGGAATTGTGAGCGGATAACAATTTCACACAGGAAACAGCTATGACCATGATTACGCCAAGCGCGC
AATTAACCCTCACTAAAGGGAACAAAAGCTGGAGCTCCACCGCGGTGGCGGCCGCCCCCTTCACCGAGGG
CCTATTTCCCATGATTCCTTCATATTTGCATATACGATACAAGGCTGTTAGAGAGATAATTGGAATTAAT
TTGACTGTAAACACAAAGATATTAGTACAAAATACGTGACGTAGAAAGTAATAATTTCTTGGGTAGTTTG
CAGTTTTAAAATTATGTTTTAAAATGGACTATCATATGCTTACCGTAACTTGAAAGTATTTCGATTTCTT
GGCTTTATATATCTTGTGGAAGGACGAAACACCGAAGATGACTATCTTTAATGTCTGTTTTAGTACTCTGGAA
ACAGAATCTACTAAAACAAGGCAAAATGCCGTGTTTATCTCGTCAACTTGTTGGCGAGAAGCTCTCATTTTCCATACATT
AAAGATAGTCATCTTTTTTTGAATTCGATATCAAGCTTATCGATACCGTCGACCTCGAGGGGGGGCCCGG
TACCCAATTCGCCCTATAGTGAGTCGTATTACGCGCGCTCACTGGCCGTCGTTTTACAACGTCGTGACTG
GGAAAACCCTGGCGTTACCCAACTTAATCGCCTTGCAGCACATCCCCCTTTCGCCAGCTGGCGTAATAGC
GAAGAGGCCCGCACCGATCGCCCTTCCCAACAGTTGCGCAGCCTGAATGGCGAATGGGACGCGCCCTGTA
GCGGCGCATTAAGCGCGGCGGGTGTGGTGGTTACGCGCAGCGTGACCGCTACACTTGCCAGCGCCCTAGC
GCCCGCTCCTTTCGCTTTCTTCCCTTCCTTTCTCGCCACGTTCGCCGGCTTTCCCCGTCAAGCTCTAAAT
CGGGGGCTCCCTTTAGGGTTCCGATTTAGTGCTTTACGGCACCTCGACCCCAAAAAACTTGATTAGGGTG
ATGGTTCACGTAGTGGGCCATCGCCCTGATAGACGGTTTTTCGCCCTTTGACGTTGGAGTCCACGTTCTT
TAATAGTGGACTCTTGTTCCAAACTGGAACAACACTCAACCCTATCTCGGTCTATTCTTTTGATTTATAA
GGGATTTTGCCGATTTCGGCCTATTGGTTAAAAAATGAGCTGATTTAACAAAAATTTAACGCGAATTTTA
ACAAAATATTAACGCTTACAATTTAG

**Supplementary Sequences 3.** Protein Sequences of prime editors

**PE2:** BPSV40_NLS-SpCas9H840A-linker-M-MLV_reverse_transcriptase-BPSV40

KRTADGSEFESPKKKRKVDKKYSIGLDIGTNSVGWAVITDEYKVPSKKFKVLGNTDRHSIKKNLIGALLF
DSGETAEATRLKRTARRRYTRRKNRICYLQEIFSNEMAKVDDSFFHRLEESFLVEEDKKHERHPIFGNIV
DEVAYHEKYPTIYHLRKKLVDSTDKADLRLIYLALAHMIKFRGHFLIEGDLNPDNSDVDKLFIQLVQTYN
QLFEENPINASGVDAKAILSARLSKSRRLENLIAQLPGEKKNGLFGNLIALSLGLTPNFKSNFDLAEDAK
LQLSKDTYDDDLDNLLAQIGDQYADLFLAAKNLSDAILLSDILRVNTEITKAPLSASMIKRYDEHHQDLT
LLKALVRQQLPEKYKEIFFDQSKNGYAGYIDGGASQEEFYKFIKPILEKMDGTEELLVKLNREDLLRKQR
TFDNGSIPHQIHLGELHAILRRQEDFYPFLKDNREKIEKILTFRIPYYVGPLARGNSRFAWMTRKSEETI
TPWNFEEVVDKGASAQSFIERMTNFDKNLPNEKVLPKHSLLYEYFTVYNELTKVKYVTEGMRKPAFLSGE
QKKAIVDLLFKTNRKVTVKQLKEDYFKKIECFDSVEISGVEDRFNASLGTYHDLLKIIKDKDFLDNEENE
DILEDIVLTLTLFEDREMIEERLKTYAHLFDDKVMKQLKRRRYTGWGRLSRKLINGIRDKQSGKTILDFL
KSDGFANRNFMQLIHDDSLTFKEDIQKAQVSGQGDSLHEHIANLAGSPAIKKGILQTVKVVDELVKVMGR
HKPENIVIEMARENQTTQKGQKNSRERMKRIEEGIKELGSQILKEHPVENTQLQNEKLYLYYLQNGRDMY
VDQELDINRLSDYDVDAIVPQSFLKDDSIDNKVLTRSDKNRGKSDNVPSEEVVKKMKNYWRQLLNAKLIT
QRKFDNLTKAERGGLSELDKAGFIKRQLVETRQITKHVAQILDSRMNTKYDENDKLIREVKVITLKSKLV
SDFRKDFQFYKVREINNYHHAHDAYLNAVVGTALIKKYPKLESEFVYGDYKVYDVRKMIAKSEQEIGKAT
AKYFFYSNIMNFFKTEITLANGEIRKRPLIETNGETGEIVWDKGRDFATVRKVLSMPQVNIVKKTEVQTG
GFSKESILPKRNSDKLIARKKDWDPKKYGGFDSPTVAYSVLVVAKVEKGKSKKLKSVKELLGITIMERSS
FEKNPIDFLEAKGYKEVKKDLIIKLPKYSLFELENGRKRMLASAGELQKGNELALPSKYVNFLYLASHYE

KLKGSPEDNEQKQLFVEQHKHYLDEIIEQISEFSKRVILADANLDKVLSAYNKHRDKPIREQAENIIHLF
TLTNLGAPAAFKYFDTTIDRKRYTSTKEVLDATLIHQSITGLYETRIDLSQLGGDSGGSSGGSSGSETPG
TSESATPESSGGSSGGSSTLNIEDEYRLHETSKEPDVSLGSTWLSDFPQAWAETGGMGLAVRQAPLIIPL
KATSTPVSIKQYPMSQEARLGIKPHIQRLLDQGILVPCQSPWNTPLLPVKKPGTNDYRPVQDLREVNKRV
EDIHPTVPNPYNLLSGLPPSHQWYTVLDLKDAFFCLRLHPTSQPLFAFEWRDPEMGISGQLTWTRLPQGF
KNSPTLFNEALHRDLADFRIQHPDLILLQYVDDLLLAATSELDCQQGTRALLQTLGNLGYRASAKKAQIC
QKQVKYLGYLLKEGQRWLTEARKETVMGQPTPKTPRQLREFLGKAGFCRLFIPGFAEMAAPLYPLTKPGT
LFNWGPDQQKAYQEIKQALLTAPALGLPDLTKPFELFVDEKQGYAKGVLTQKLGPWRRPVAYLSKKLDPV
AAGWPPCLRMVAAIAVLTKDAGKLTMGQPLVILAPHAVEALVKQPPDRWLSNARMTHYQALLLDTDRVQF
GPVVALNPATLLPLPEEGLQHNCLDILAEAHGTRPDLTDQPLPDADHTWYTDGSSLLQEGQRKAGAAVTT
ETEVIWAKALPAGTSAQRAELIALTQALKMAEGKKLNVYTDSRYAFATAHIHGEIYRRRGWLTSEGKEIK
NKDEILALLKALFLPKRLSIIHCPGHQKGHSAEARGNRMADQAARKAAITETPDTSTLLIENSSPSGGSK
RTADGSEFEPKKKRKV

**PE2*:** Cmyc_NLS-BPSV40_NLS-SpCas9H840A-linker-M-MLV_reverse_transcriptase-vBPSV40_NLS-SV40

PAAKRVKLDGGKRTADGSEFESPKKKRKVDKKYSIGLDIGTNSVGWAVITDEYKVPSKKFKVLGNTDRHS
IKKNLIGALLFDSGETAEATRLKRTARRRYTRRKNRICYLQEIFSNEMAKVDDSFFHRLEESFLVEEDKK
HERHPIFGNIVDEVAYHEKYPTIYHLRKKLVDSTDKADLRLIYLALAHMIKFRGHFLIEGDLNPDNSDVD
KLFIQLVQTYNQLFEENPINASGVDAKAILSARLSKSRRLENLIAQLPGEKKNGLFGNLIALSLGLTPNF
KSNFDLAEDAKLQLSKDTYDDDLDNLLAQIGDQYADLFLAAKNLSDAILLSDILRVNTEITKAPLSASMI
KRYDEHHQDLTLLKALVRQQLPEKYKEIFFDQSKNGYAGYIDGGASQEEFYKFIKPILEKMDGTEELLVK
LNREDLLRKQRTFDNGSIPHQIHLGELHAILRRQEDFYPFLKDNREKIEKILTFRIPYYVGPLARGNSRF
AWMTRKSEETITPWNFEEVVDKGASAQSFIERMTNFDKNLPNEKVLPKHSLLYEYFTVYNELTKVKYVTE
GMRKPAFLSGEQKKAIVDLLFKTNRKVTVKQLKEDYFKKIECFDSVEISGVEDRFNASLGTYHDLLKIIK
DKDFLDNEENEDILEDIVLTLTLFEDREMIEERLKTYAHLFDDKVMKQLKRRRYTGWGRLSRKLINGIRD
KQSGKTILDFLKSDGFANRNFMQLIHDDSLTFKEDIQKAQVSGQGDSLHEHIANLAGSPAIKKGILQTVK
VVDELVKVMGRHKPENIVIEMARENQTTQKGQKNSRERMKRIEEGIKELGSQILKEHPVENTQLQNEKLY
LYYLQNGRDMYVDQELDINRLSDYDVDAIVPQSFLKDDSIDNKVLTRSDKNRGKSDNVPSEEVVKKMKNY
WRQLLNAKLITQRKFDNLTKAERGGLSELDKAGFIKRQLVETRQITKHVAQILDSRMNTKYDENDKLIRE
VKVITLKSKLVSDFRKDFQFYKVREINNYHHAHDAYLNAVVGTALIKKYPKLESEFVYGDYKVYDVRKMI
AKSEQEIGKATAKYFFYSNIMNFFKTEITLANGEIRKRPLIETNGETGEIVWDKGRDFATVRKVLSMPQV
NIVKKTEVQTGGFSKESILPKRNSDKLIARKKDWDPKKYGGFDSPTVAYSVLVVAKVEKGKSKKLKSVKE
LLGITIMERSSFEKNPIDFLEAKGYKEVKKDLIIKLPKYSLFELENGRKRMLASAGELQKGNELALPSKY
VNFLYLASHYEKLKGSPEDNEQKQLFVEQHKHYLDEIIEQISEFSKRVILADANLDKVLSAYNKHRDKPI
REQAENIIHLFTLTNLGAPAAFKYFDTTIDRKRYTSTKEVLDATLIHQSITGLYETRIDLSQLGGDSGGS
SGGSSGSETPGTSESATPESSGGSSGGSSTLNIEDEYRLHETSKEPDVSLGSTWLSDFPQAWAETGGMGL
AVRQAPLIIPLKATSTPVSIKQYPMSQEARLGIKPHIQRLLDQGILVPCQSPWNTPLLPVKKPGTNDYRP
VQDLREVNKRVEDIHPTVPNPYNLLSGLPPSHQWYTVLDLKDAFFCLRLHPTSQPLFAFEWRDPEMGISG
QLTWTRLPQGFKNSPTLFNEALHRDLADFRIQHPDLILLQYVDDLLLAATSELDCQQGTRALLQTLGNLG
YRASAKKAQICQKQVKYLGYLLKEGQRWLTEARKETVMGQPTPKTPRQLREFLGKAGFCRLFIPGFAEMA
APLYPLTKPGTLFNWGPDQQKAYQEIKQALLTAPALGLPDLTKPFELFVDEKQGYAKGVLTQKLGPWRRP
VAYLSKKLDPVAAGWPPCLRMVAAIAVLTKDAGKLTMGQPLVILAPHAVEALVKQPPDRWLSNARMTHYQ
ALLLDTDRVQFGPVVALNPATLLPLPEEGLQHNCLDILAEAHGTRPDLTDQPLPDADHTWYTDGSSLLQE
GQRKAGAAVTTETEVIWAKALPAGTSAQRAELIALTQALKMAEGKKLNVYTDSRYAFATAHIHGEIYRRR

GWLTSEGKEIKNKDEILALLKALFLPKRLSIIHCPGHQKGHSAEARGNRMADQAARKAAITETPDTSTLL
IENSSPSGGSKRTADGSEKRTADSQHSTPPKTKRKVEFEPKKKRKV

**SaPE2*:** Cmyc_NLS-BPSV40_NLS-SaCas9N580A-linker-M-MLV_reverse_transcriptase-vBPSV40_NLS-SV40

PAAKRVKLDGGKRTADGSEFESPKKKRKVGIHGVPAAKRNYILGLDIGITSVGYGIIDYETRDVIDAGVR
LFKEANVENNEGRRSKRGARRLKRRRRHRIQRVKKLLFDYNLLTDHSELSGINPYEARVKGLSQKLSEEE
FSAALLHLAKRRGVHNVNEVEEDTGNELSTKEQISRNSKALEEKYVAELQLERLKKDGEVRGSINRFKTS
DYVKEAKQLLKVQKAYHQLDQSFIDTYIDLLETRRTYYEGPGEGSPFGWKDIKEWYEMLMGHCTYFPEEL
RSVKYAYNADLYNALNDLNNLVITRDENEKLEYYEKFQIIENVFKQKKKPTLKQIAKEILVNEEDIKGYR
VTSTGKPEFTNLKVYHDIKDITARKEIIENAELLDQIAKILTIYQSSEDIQEELTNLNSELTQEEIEQIS
NLKGYTGTHNLSLKAINLILDELWHTNDNQIAIFNRLKLVPKKVDLSQQKEIPTTLVDDFILSPVVKRSF
IQSIKVINAIIKKYGLPNDIIIELAREKNSKDAQKMINEMQKRNRQTNERIEEIIRTTGKENAKYLIEKI
KLHDMQEGKCLYSLEAIPLEDLLNNPFNYEVDHIIPRSVSFDNSFNNKVLVKQEEASKKGNRTPFQYLSS
SDSKISYETFKKHILNLAKGKGRISKTKKEYLLEERDINRFSVQKDFINRNLVDTRYATRGLMNLLRSYF
RVNNLDVKVKSINGGFTSFLRRKWKFKKERNKGYKHHAEDALIIANADFIFKEWKKLDKAKKVMENQMFE
EKQAESMPEIETEQEYKEIFITPHQIKHIKDFKDYKYSHRVDKKPNRELINDTLYSTRKDDKGNTLIVNN
LNGLYDKDNDKLKKLINKSPEKLLMYHHDPQTYQKLKLIMEQYGDEKNPLYKYYEETGNYLTKYSKKDNG
PVIKKIKYYGNKLNAHLDITDDYPNSRNKVVKLSLKPYRFDVYLDNGVYKFVTVKNLDVIKKENYYEVNS
KCYEEAKKLKKISNQAEFIASFYNNDLIKINGELYRVIGVNNDLLNRIEVNMIDITYREYLENMNDKRPP
RIIKTIASKTQSIKKYSTDILGNLYEVKSKKHPQIIKKGSGGSSGGSSGSETPGTSESATPESSGGSSGG
SSTLNIEDEYRLHETSKEPDVSLGSTWLSDFPQAWAETGGMGLAVRQAPLIIPLKATSTPVSIKQYPMSQ
EARLGIKPHIQRLLDQGILVPCQSPWNTPLLPVKKPGTNDYRPVQDLREVNKRVEDIHPTVPNPYNLLSG
LPPSHQWYTVLDLKDAFFCLRLHPTSQPLFAFEWRDPEMGISGQLTWTRLPQGFKNSPTLFNEALHRDLA
DFRIQHPDLILLQYVDDLLLAATSELDCQQGTRALLQTLGNLGYRASAKKAQICQKQVKYLGYLLKEGQR
WLTEARKETVMGQPTPKTPRQLREFLGKAGFCRLFIPGFAEMAAPLYPLTKPGTLFNWGPDQQKAYQEIK
QALLTAPALGLPDLTKPFELFVDEKQGYAKGVLTQKLGPWRRPVAYLSKKLDPVAAGWPPCLRMVAAIAV
LTKDAGKLTMGQPLVILAPHAVEALVKQPPDRWLSNARMTHYQALLLDTDRVQFGPVVALNPATLLPLPE
EGLQHNCLDILAEAHGTRPDLTDQPLPDADHTWYTDGSSLLQEGQRKAGAAVTTETEVIWAKALPAGTSA
QRAELIALTQALKMAEGKKLNVYTDSRYAFATAHIHGEIYRRRGWLTSEGKEIKNKDEILALLKALFLPK
RLSIIHCPGHQKGHSAEARGNRMADQAARKAAITETPDTSTLLIENSSPSGGSKRTADGSEKRTADSQHS
TPPKTKRKVEFEPKKKRKV

**Sa$^{KKH}$PE*:** Cmyc_NLS-BPSV40_NLS-SaCas9$^{KKH}$N580A-linker-M-MLV_reverse_transcriptase-vBPSV40_NLS-SV40

PAAKRVKLDGGKRTADGSEFESPKKKRKVGIHGVPAAKRNYILGLDIGITSVGYGIIDYETRDVIDAGVR
LFKEANVENNEGRRSKRGARRLKRRRRHRIQRVKKLLFDYNLLTDHSELSGINPYEARVKGLSQKLSEEE
FSAALLHLAKRRGVHNVNEVEEDTGNELSTKEQISRNSKALEEKYVAELQLERLKKDGEVRGSINRFKTS
DYVKEAKQLLKVQKAYHQLDQSFIDTYIDLLETRRTYYEGPGEGSPFGWKDIKEWYEMLMGHCTYFPEEL
RSVKYAYNADLYNALNDLNNLVITRDENEKLEYYEKFQIIENVFKQKKKPTLKQIAKEILVNEEDIKGYR
VTSTGKPEFTNLKVYHDIKDITARKEIIENAELLDQIAKILTIYQSSEDIQEELTNLNSELTQEEIEQIS
NLKGYTGTHNLSLKAINLILDELWHTNDNQIAIFNRLKLVPKKVDLSQQKEIPTTLVDDFILSPVVKRSF
IQSIKVINAIIKKYGLPNDIIIELAREKNSKDAQKMINEMQKRNRQTNERIEEIIRTTGKENAKYLIEKI
KLHDMQEGKCLYSLEAIPLEDLLNNPFNYEVDHIIPRSVSFDNSFNNKVLVKQEEASKKGNRTPFQYLSS

SDSKISYETFKKHILNLAKGKGRISKTKKEYLLEERDINRFSVQKDFINRNLVDTRYATRGLMNLLRSYF
RVNNLDVKVKSINGGFTSFLRRKWKFKKERNKGYKHHAEDALIIANADFIFKEWKKLDKAKKVMENQMFE
EKQAESMPEIETEQEYKEIFITPHQIKHIKDFKDYKYSHRVDKKPNRKLINDTLYSTRKDDKGNTLIVNN
LNGLYDKDNDKLKKLINKSPEKLLMYHHDPQTYQKLKLIMEQYGDEKNPLYKYYEETGNYLTKYSKKDNG
PVIKKIKYYGNKLNAHLDITDDYPNSRNKVVKLSLKPYRFDVYLDNGVYKFVTVKNLDVIKKENYYEVNS
KCYEEAKKLKKISNQAEFIASFYKNDLIKINGELYRVIGVNNDLLNRIEVNMIDITYREYLENMNDKRPP
HIIKTIASKTQSIKKYSTDILGNLYEVKSKKHPQIIKKGSGGSSGGSSGSETPGTSESATPESSGGSSGG
SSTLNIEDEYRLHETSKEPDVSLGSTWLSDFPQAWAETGGMGLAVRQAPLIIPLKATSTPVSIKQYPMSQ
EARLGIKPHIQRLLDQGILVPCQSPWNTPLLPVKKPGTNDYRPVQDLREVNKRVEDIHPTVPNPYNLLSG
LPPSHQWYTVLDLKDAFFCLRLHPTSQPLFAFEWRDPEMGISGQLTWTRLPQGFKNSPTLFNEALHRDLA
DFRIQHPDLILLQYVDDLLLAATSELDCQQGTRALLQTLGNLGYRASAKKAQICQKQVKYLGYLLKEGQR
WLTEARKETVMGQPTPKTPRQLREFLGKAGFCRLFIPGFAEMAAPLYPLTKPGTLFNWGPDQQKAYQEIK
QALLTAPALGLPDLTKPFELFVDEKQGYAKGVLTQKLGPWRRPVAYLSKKLDPVAAGWPPCLRMVAAIAV
LTKDAGKLTMGQPLVILAPHAVEALVKQPPDRWLSNARMTHYQALLLDTDRVQFGPVVALNPATLLPLPE
EGLQHNCLDILAEAHGTRPDLTDQPLPDADHTWYTDGSSLLQEGQRKAGAAVTTETEVIWAKALPAGTSA
QRAELIALTQALKMAEGKKLNVYTDSRYAFATAHIHGEIYRRRGWLTSEGKEIKNKDEILALLKALFLPK
RLSIIHCPGHQKGHSAEARGNRMADQAARKAAITETPDTSTLLIENSSPSGGSKRTADGSEKRTADSQHS
TPPKTKRKVEFEPKKKRKV

pU6-Ctnnb1_pegRNA_S45F: U6 promoter + Spacer + sgRNA scaffold + RT +PBS

GAGGGCCTATTTCCCATGATTCCTTCATATTTGCATATACGATACAAGGCTGTTAGAGAGAT
AATTGGAATTAATTTGACTGTAAACACAAAGATATTAGTACAAAATACGTGACGTAGAAAGT
AATAATTTCTTGGGTAGTTTGCAGTTTTAAAATTATGTTTTAAAATGGACTATCATATGCTTA
CCGTAACTTGAAAGTATTTCGATTTCTTGGCTTTATATATCTTGTGGAAAGGACGAAACACC
GAGGGTTGCCCTTGCCACTCAgttttagagctagaaatagcaagttaaaataaggctagtccgttatcaacttgaaaaa
gtgggaccgagtcggtccGCTCCTTTCCTGAGTGGCAAGGGCAATTTTTTT

U6-pegAAT-U6-Nicking-U1A-Cmyc_NLS-SpCas9(N)-Nter-Npu-intein

cctgcaggcagctgcgcgctcgctcgctcactgaggccgcccgggcaaagcccgggcgtcgggcgacctt
tggtcgcccggcctcagtgagcgagcgagcgcgcagagagggagtggccaactccatcactaggggttcc
tgcggcctctagaggtaccGAGGGCCTATTTCCCATGATTCCTTCATATTTGCATATACGATACAAGGCT
GTTAGAGAGATAATTGGAATTAATTTGACTGTAAACACAAAGATATTAGTACAAAATACGTGACGTAGAA
AGTAATAATTTCTTGGGTAGTTTGCAGTTTTAAAATTATGTTTTAAAATGGACTATCATATGCTTACCGT
AACTTGAAAGTATTTCGATTTCTTGGCTTTATATATCTTGTGGAAAGGACGAAACACCGtcccctccagg
ccgtgcatagttttagagctagaaatagcaagttaaaataaggctagtccgttatcaacttgaaaaagtg
ggaccgagtcggtcctctcgtcgatggtcagcacagcTttatgcacggcctggagTTTTTTTGAATTCGA
TATCAAGCTTATCGATACCGTCGACCTCGgctgccgctggaggtgctcaaagagatggaGAGGGCCTATT
TCCCATGATTCCTTCATATTTGCATATACGATACAAGGCTGTTAGAGAGATAATTGGAATTAATTTGACT
GTAAACACAAAGATATTAGTACAAAATACGTGACGTAGAAAGTAATAATTTCTTGGGTAGTTTGCAGTTT
TAAAATTATGTTTTAAAATGGACTATCATATGCTTACCGTAACTTGAAAGTATTTCGATTTCTTGGCTTT
ATATATCTTGTGGAAAGGACGAAACACCGTTCAATCATTAAGAAGACAAgttttagagctagaaatagca
agttaaaataaggctagtccgttatcaacttgaaaaagtgggaccgagtcggtgcTTTTTTTGAATTCGA
TATCAAGCTTATCGATACCGTCgctagcccgggtcgactagtgatcagtgtgagggagtgtaaagctggt
ttaaacgttgttgtttggggttgaattactcttctagaccgcggcgcgcctccatggatatcaagcttat

ggaggcggtactatgtagatgagaattcaggagcaaactgggaaaagcaactgcttccaaatatttgtga
tttttacagtgtagttttggaaaaactcttagcctaccaattcttctaagtgttttaaaatgtgggagcc
agtacacatgaagttatagagtgttttaatgaggcttaaatatttaccgtaactatgaaatgctacgcat
atcatgctgttcaggctccgtggccacgcaactcatactaccggtGCCACCATGgctagcCCCGCCGCCA
AGCGCGTGAAGCTGGACGACAAGAAGTACAGCATCGGCCTGGACATCGGCACCAACTCTGTGGGCTGGGC
CGTGATCACCGACGAGTACAAGGTGCCCAGCAAGAAATTCAAGGTGCTGGGCAACACCGACCGGCACAGC
ATCAAGAAGAACCTGATCGGAGCCCTGCTGTTCGACAGCGGCGAAACAGCCGAGGCCACCCGGCTGAAGA
GAACCGCCAGAAGAAGATACACCAGACGGAAGAACCGGATCTGCTATCTGCAAGAGATCTTCAGCAACGA
GATGGCCAAGGTGGACGACAGCTTCTTCCACAGACTGGAAGAGTCCTTCCTGGTGGAAGAGGATAAGAAG
CACGAGCGGCACCCCATCTTCGGCAACATCGTGGACGAGGTGGCCTACCACGAGAAGTACCCCACCATCT
ACCACCTGAGAAAGAAACTGGTGGACAGCACCGACAAGGCCGACCTGCGGCTGATCTATCTGGCCCTGGC
CCACATGATCAAGTTCCGGGGCCACTTCCTGATCGAGGGCGACCTGAACCCCGACAACAGCGACGTGGAC
AAGCTGTTCATCCAGCTGGTGCAGACCTACAACCAGCTGTTCGAGGAAAACCCCATCAACGCCAGCGGCG
TGGACGCCAAGGCCATCCTGTCTGCCAGACTGAGCAAGAGCAGACGGCTGGAAAATCTGATCGCCCAGCT
GCCCGGCGAGAAGAAGAATGGCCTGTTCGGAAACCTGATTGCCCTGAGCCTGGGCCTGACCCCCAACTTC
AAGAGCAACTTCGACCTGGCCGAGGATGCCAAACTGCAGCTGAGCAAGGACACCTACGACGACGACCTGG
ACAACCTGCTGGCCCAGATCGGCGACCAGTACGCCGACCTGTTTCTGGCCGCCAAGAACCTGTCCGACGC
CATCCTGCTGAGCGACATCCTGAGAGTGAACACCGAGATCACCAAGGCCCCCCTGAGCGCCTCTATGATC
AAGAGATACGACGAGCACCACCAGGACCTGACCCTGCTGAAAGCTCTCGTGCGGCAGCAGCTGCCTGAGA
AGTACAAAGAGATTTTCTTCGACCAGAGCAAGAACGGCTACGCCGGCTACATTGACGGCGGAGCCAGCCA
GGAAGAGTTCTACAAGTTCATCAAGCCCATCCTGGAAAAGATGGACGGCACCGAGGAACTGCTCGTGAAG
CTGAACAGAGAGGACCTGCTGCGGAAGCAGCGGACCTTCGACAACGGCAGCATCCCCCACCAGATCCACC
TGGGAGAGCTGCACGCCATTCTGCGGCGGCAGGAAGATTTTTACCCATTCCTGAAGGACAACCGGGAAAA
GATCGAGAAGATCCTGACCTTCCGCATCCCCTACTACGTGGGCCCTCTGGCCAGGGGAAACAGCAGATTC
GCCTGGATGACCAGAAAGAGCGAGGAAACCATCACCCCCTGGAACTTCGAGGAAGTGGTGGACAAGGGCG
CTTCCGCCCAGAGCTTCATCGAGCGGATGACCAACTTCGATAAGAACCTGCCCAACGAGAAGGTGCTGCC
CAAGCACAGCCTGCTGTACGAGTACTTCACCGTGTATAACGAGCTGACCAAAGTGAAATACGTGACCGAG
GGAATGAGAAAGCCCGCCTTCCTGAGCGGCGAGCAGAAAAAGGCCATCGTGGACCTGCTGTTCAAGACCA
ACCGGAAAGTGACCGTGAAGCAGCTGAAAGAGGACTACTTCAAGAAAATCGAGTGCTTCGACTCCGTGGA
AATCTCCGGCGTGGAAGATCGGTTCAACGCCTCCCTGGGCACATACCACGATCTGCTGAAAATTATCAAG
GACAAGGACTTCCTGGACAATGAGGAAAACGAGGACATTCTGGAAGATATCGTGCTGACCCTGACACTGT
TTGAGGACAGAGAGATGATCGAGGAACGGCTGAAAACCTATGCCCACCTGTTCGACGACAAAGTGATGAA
GCAGCTGAAGCGGCGGAGATACACCGGCTGGGGCAGGCTGAGCCGGAAGCTGATCAACGGCATCCGGGAC
AAGCAGTCCGGCAAGACAATCCTGGATTTCCTGAAGTCCGACGGCTTCGCCAACAGAAACTTCATGCAGC
TGATCCACGACGACAGCCTGACCTTTAAAGAGGACATCCAGAAAGCCCAGGTGTGCCTGTCCTACGAGAC
AGAGATCCTGACAGTGGAGTATGGCCTGCTGCCAATCGGCAAGATCGTGGAGAAGAGGATCGAGTGTACC
GTGTACTCTGTGGATAACAATGGCAACATCTATACACAGCCCGTGGCACAGTGGCACGATAGGGGAGAGC
AGGAGGTGTTCGAGTATTGCCTGGAGGACGGCAGCCTGATCAGGGCAACCAAGGACCACAAGTTCATGAC
AGTGGATGGCCAGATGCTGCCCATCGACGAGATTTTCGAGCGGGAGCTGGACCTGATGAGAGTGGATAAC
CTGCCTAATtgagaattcctagagctcgctgatcagcctcgactgtgccttctagttgccagccatctgt
tgtttgcccctcccccgtgccttccttgaccctggaaggtgccactcccactgtcctttcctaataaaat
gaggaaattgcatcgcattgtctgagtaggtgtcattctattctggggggtggggtggggcaggacagca
agggggaggattgggaagagaatagcaggcatgctggggagcggccgcaggaacccctagtgatggagtt
ggccactccctctctgcgcgctcgctcgctcactgaggccgggcgaccaaaggtcgcccgacgcccgggc
tttgcccgggcggcctcagtgagcgagcgagcgcgcagctgcctgcaggggcgcctgatgcggtattttc
tccttacgcatctgtgcggtatttcacaccgcatacgtcaaagcaaccatagtacgcgccctgtagcggc
gcattaagcgcggcgggtgtggtggttacgcgcagcgtgaccgctacacttgccagcgccctagcgcccg
ctcctttcgctttcttcccttcctttctcgccacgttcgccggctttccccgtcaagctctaaatcgggg

gctccctttagggttccgatttagtgctttacggcacctcgaccccaaaaaacttgatttgggtgatggt
tcacgtagtgggccatcgccctgatagacggtttttcgccctttgacgttggagtccacgttctttaata
gtggactcttgttccaaactggaacaacactcaaccctatctcgggctattcttttgatttataagggat
tttgccgatttcggcctattggttaaaaaatgagctgatttaacaaaaatttaacgcgaattttaacaaa
atattaacgtttacaattttatggtgcactctcagtacaatctgctctgatgccgcatagttaagccagc
cccgacacccgccaacacccgctgacgcgccctgacgggcttgtctgctcccggcatccgcttacagaca
agctgtgaccgtctccgggagctgcatgtgtcagaggttttcaccgtcatcaccgaaacgcgcgagacga
aagggcctcgtgatacgcctatttttataggttaatgtcatgataataatggtttcttagacgtcaggtg
gcacttttcggggaaatgtgcgcggaacccctatttgtttatttttctaaatacattcaaatatgtatcc
gctcatgagacaataaccctgataaatgcttcaataatattgaaaaaggaagagtatgagtattcaacat
ttccgtgtcgcccttattcccttttttgcggcattttgccttcctgtttttgctcacccagaaacgctgg
tgaaagtaaaagatgctgaagatcagttgggtgcacgagtgggttacatcgaactggatctcaacagcgg
taagatccttgagagttttcgccccgaagaacgttttccaatgatgagcacttttaaagttctgctatgt
ggcgcggtattatcccgtattgacgccgggcaagagcaactcggtcgccgcatacactattctcagaatg
acttggttgagtactcaccagtcacagaaaagcatcttacggatggcatgacagtaagagaattatgcag
tgctgccataaccatgagtgataacactgcggccaacttacttctgacaacgatcggaggaccgaaggag
ctaaccgcttttttgcacaacatgggggatcatgtaactcgccttgatcgttgggaaccggagctgaatg
aagccataccaaacgacgagcgtgacaccacgatgcctgtagcaatggcaacaacgttgcgcaaactatt
aactggcgaactacttactctagcttcccggcaacaattaatagactggatggaggcggataaagttgca
ggaccacttctgcgctcggcccttccggctggctggtttattgctgataaatctggagccggtgagcgtg
gaagccgcggtatcattgcagcactggggccagatggtaagccctcccgtatcgtagttatctacacgac
ggggagtcaggcaactatggatgaacgaaatagacagatcgctgagataggtgcctcactgattaagcat
tggtaactgtcagaccaagtttactcatatatactttagattgatttaaaacttcatttttaatttaaaa
ggatctaggtgaagatcctttttgataatctcatgaccaaaatcccttaacgtgagttttcgttccactg
agcgtcagaccccgtagaaaagatcaaaggatcttcttgagatcctttttttctgcgcgtaatctgctgc
ttgcaaacaaaaaaaccaccgctaccagcggtggtttgtttgccggatcaagagctaccaactctttttc
cgaaggtaactggcttcagcagagcgcagataccaaatactgtccttctagtgtagccgtagttaggcca
ccacttcaagaactctgtagcaccgcctacatacctcgctctgctaatcctgttaccagtggctgctgcc
agtggcgataagtcgtgtcttaccgggttggactcaagacgatagttaccggataaggcgcagcggtcgg
gctgaacggggggttcgtgcacacagcccagcttggagcgaacgacctacaccgaactgagatacctaca
gcgtgagctatgagaaagcgccacgcttcccgaagggagaaaggcggacaggtatccggtaagcggcagg
gtcggaacaggagagcgcacgagggagcttccagggggaaacgcctggtatctttatagtcctgtcgggt
ttcgccacctctgacttgagcgtcgatttttgtgatgctcgtcaggggggcggagcctatggaaaaacgc
cagcaacgcggcctttttacggttcctggccttttgctggccttttgctcacatgt

Cter-Npu-intein-SpCas9(C)- linker- M-MLV_reverse_transcriptase-BPSV40_NLS

cctgcaggcagctgcgcgctcgctcgctcactgaggccgcccgggcaaagcccgggcgtcgggcgacctt
tggtcgcccggcctcagtgagcgagcgagcgcgcagagagggagtggccaactccatcactaggggttcc
ttctagaAtggaggcggtactatgtagatgagaattcaggagcaaactgggaaaagcaactgcttccaaa
tatttgtgatttttacagtgtagttttggaaaaactcttagcctaccaattcttctaagtgttttaaaat
gtgggagccagtacacatgaagttatagagtgttttaatgaggcttaaatatttaccgtaactatgaaat
gctacgcatatcatgctgttcaggctccgtggccacgcaactcatactaccggtGCCACCATGATCAAGA
TTGCTACACGGAAATACCTGGGAAAGCAGAACGTGTACGACATCGGCGTGGAGCGGATCACAACTTCGC
CCTGAAGAATGGCTTTATCGCCAGCAATTCCGGCCAGGGCGATAGCCTGCACGAGCACATTGCCAATCTG
GCCGGCAGCCCCGCCATTAAGAAGGGCATCCTGCAGACAGTGAAGGTGGTGGACGAGCTCGTGAAAGTGA
TGGGCCGGCACAAGCCCGAGAACATCGTGATCGAAATGGCCAGAGAGAACCAGACCACCCAGAAGGACA

GAAGAACAGCCGCGAGAGAATGAAGCGGATCGAAGAGGGCATCAAAGAGCTGGGCAGCCAGATCCTGAAA
GAACACCCCGTGGAAAACACCCAGCTGCAGAACGAGAAGCTGTACCTGTACTACCTGCAGAATGGGCGGG
ATATGTACGTGGACCAGGAACTGGACATCAACCGGCTGTCCGACTACGATGTGGACGCTATCGTGCCTCA
GAGCTTTCTGAAGGACGACTCCATCGACAACAAGGTGCTGACCAGAAGCGACAAGAACCGGGGCAAGAGC
GACAACGTGCCCTCCGAAGAGGTCGTGAAGAAGATGAAGAACTACTGGCGGCAGCTGCTGAACGCCAAGC
TGATTACCCAGAGAAAGTTCGACAATCTGACCAAGGCCGAGAGAGGCGGCCTGAGCGAACTGGATAAGGC
CGGCTTCATCAAGAGACAGCTGGTGGAAACCCGGCAGATCACAAAGCACGTGGCACAGATCCTGGACTCC
CGGATGAACACTAAGTACGACGAGAATGACAAGCTGATCCGGGAAGTGAAAGTGATCACCCTGAAGTCCA
AGCTGGTGTCCGATTTCCGGAAGGATTTCCAGTTTTACAAAGTGCGCGAGATCAACAACTACCACCACGC
CCACGACGCCTACCTGAACGCCGTCGTGGGAACCGCCCTGATCAAAAAGTACCCTAAGCTGGAAAGCGAG
TTCGTGTACGGCGACTACAAGGTGTACGACGTGCGGAAGATGATCGCCAAGAGCGAGCAGGAAATCGGCA
AGGCTACCGCCAAGTACTTCTTCTACAGCAACATCATGAACTTTTTCAAGACCGAGATTACCCTGGCCAA
CGGCGAGATCCGGAAGCGGCCTCTGATCGAGACAAACGGCGAAACCGGGGAGATCGTGTGGGATAAGGGC
CGGGATTTTGCCACCGTGCGGAAAGTGCTGAGCATGCCCCAAGTGAATATCGTGAAAAAGACCGAGGTGC
AGACAGGCGGCTTCAGCAAAGAGTCTATCCTGCCCAAGAGGAACAGCGATAAGCTGATCGCCAGAAAGAA
GGACTGGGACCCTAAGAAGTACGGCGGCTTCGACAGCCCCACCGTGGCCTATTCTGTGCTGGTGGTGGCC
AAAGTGGAAAAGGGCAAGTCCAAGAAACTGAAGAGTGTGAAAGAGCTGCTGGGGATCACCATCATGGAAA
GAAGCAGCTTCGAGAAGAATCCCATCGACTTTCTGGAAGCCAAGGGCTACAAAGAAGTGAAAAAGGACCT
GATCATCAAGCTGCCTAAGTACTCCCTGTTCGAGCTGGAAAACGGCCGGAAGAGAATGCTGGCCTCTGCC
GGCGAACTGCAGAAGGGAAACGAACTGGCCCTGCCCTCCAAATATGTGAACTTCCTGTACCTGGCCAGCC
ACTATGAGAAGCTGAAGGGCTCCCCCGAGGATAATGAGCAGAAACAGCTGTTTGTGGAACAGCACAAGCA
CTACCTGGACGAGATCATCGAGCAGATCAGCGAGTTCTCCAAGAGAGTGATCCTGGCCGACGCTAATCTG
GACAAAGTGCTGTCCGCCTACAACAAGCACCGGGATAAGCCCATCAGAGAGCAGGCCGAGAATATCATCC
ACCTGTTTACCCTGACCAATCTGGGAGCCCCTGCCGCCTTCAAGTACTTTGACACCACCATCGACCGGAA
GAGGTACACCAGCACCAAAGAGGTGCTGGACGCCACCCTGATCCACCAGAGCATCACCGGCCTGTACGAG
ACACGGATCGACCTGTCTCAGCTGGGAGGTGACTCTGGAGGATCTAGCGGAGGATCCTCTGGCAGCGAGA
CACCAGGAACAAGCGAGTCAGCAACACCAGAGAGCAGTGGCGGCAGCAGCGGCGGCAGCAGCACCCTAAA
TATAGAAGATGAGTATCGGCTACATGAGACCTCAAAAGAGCCAGATGTTTCTCTAGGGTCCACATGGCTG
TCTGATTTTCCTCAGGCCTGGGCGGAAACCGGGGGCATGGGACTGGCAGTTCGCCAAGCTCCTCTGATCA
TACCTCTGAAAGCAACCTCTACCCCCGTGTCCATAAAACAATACCCCATGTCACAAGAAGCCAGACTGGG
GATCAAGCCCCACATACAGAGACTGTTGGACCAGGGAATACTGGTACCCTGCCAGTCCCCCTGGAACACG
CCCCTGCTACCCGTTAAGAAACCAGGGACTAATGATTATAGGCCTGTCCAGGATCTGAGAGAAGTCAACA
AGCGGGTGGAAGACATCCACCCCACCGTGCCCAACCCTTACAACCTCTTGAGCGGGCTCCCACCGTCCCA
CCAGTGGTACACTGTGCTTGATTTAAAGGATGCCTTTTTCTGCCTGAGACTCCACCCCACCAGTCAGCCT
CTCTTCGCCTTTGAGTGGAGAGATCCAGAGATGGGAATCTCAGGACAATTGACCTGGACCAGACTCCCAC
AGGGTTTCAAAAACAGTCCCACCCTGTTTAATGAGGCACTGCACAGAGACCTAGCAGACTTCCGGATCCA
GCACCCAGACTTGATCCTGCTACAGTACGTGGATGACTTACTGCTGGCCGCCACTTCTGAGCTAGACTGC
CAACAAGGTACTCGGGCCCTGTTACAAACCCTAGGGAACCTCGGGTATCGGGCCTCGGCCAAGAAAGCCC
AAATTTGCCAGAAACAGGTCAAGTATCTGGGGTATCTTCTAAAAGAGGGTCAGAGATGGCTGACTGAGGC
CAGAAAAGAGACTGTGATGGGGCAGCCTACTCCGAAGACCCCTCGACAACTAAGGGAGTTCCTAGGGAAG
GCAGGCTTCTGTCGCCTCTTCATCCCTGGGTTTGCAGAAATGGCAGCCCCCCTGTACCCTCTCACCAAAC
CGGGGACTCTGTTTAATTGGGGCCCAGACCAACAAAAGGCCTATCAAGAAATCAAGCAAGCTCTTCTAAC
TGCCCCAGCCCTGGGGTTGCCAGATTTGACTAAGCCCTTTGAACTCTTTGTCGACGAGAAGCAGGGCTAC
GCCAAAGGTGTCCTAACGCAAAAACTGGGACCTTGGCGTCGGCCGGTGGCCTACCTGTCCAAAAAGCTAG
ACCCAGTAGCAGCTGGGTGGCCCCCTTGCCTACGGATGGTAGCAGCCATTGCCGTACTGACAAAGGATGC
AGGCAAGCTAACCATGGGACAGCCACTAGTCATTCTGGCCCCCCATGCAGTAGAGGCACTAGTCAAACAA
CCCCCCGACCGCTGGCTTTCCAACGCCCGGATGACTCACTATCAGGCCTTGCTTTTGGACACGGACCGGG
TCCAGTTCGGACCGGTGGTAGCCCTGAACCCGGCTACGCTGCTCCCACTGCCTGAGGAAGGGCTGCAACA

CAACTGCCTTGATATCCTGGCCGAAGCCCACGGAACCCGACCCGACCTAACGGACCAGCCGCTCCCAGAC
GCCGACCACACCTGGTACACGGATGGAAGCAGTCTCTTACAAGAGGGACAGCGTAAGGCGGGAGCTGCGG
TGACCACCGAGACCGAGGTAATCTGGGCTAAAGCCCTGCCAGCCGGGACATCCGCTCAGCGGGCTGAACT
GATAGCACTCACCCAGGCCCTAAAGATGGCAGAAGGTAAGAAGCTAAATGTTTATACTGATAGCCGTTAT
GCTTTTGCTACTGCCCATATCCATGGAGAAATATACAGAAGGCGTGGGTGGCTCACATCAGAAGGCAAAG
AGATCAAAAATAAAGACGAGATCTTGGCCCTACTAAAAGCCCTCTTTCTGCCCAAAAGACTTAGCATAAT
CCATTGTCCAGGACATCAAAAGGGACACAGCGCCGAGGCTAGAGGCAACCGGATGGCTGACCAAGCGGCC
CGAAAGGCAGCCATCACAGAGACTCCAGACACCTCTACCCTCCTCATAGAAAATTCATCACCCTCTGGCG
GCTCAAAAAGAACCGCCGACGGCAGCGAATTCGAGCCCAAGAAGAAGAGGAAAGTCTAAgaaaataaagg
aaatttattttcattgcaatagtgtgttggaattttttgtgtctctcagcggccgcaggaacccctagtg
atggagttggccactccctctctgcgcgctcgctcgctcactgaggccgggcgaccaaaggtcgcccgac
gcccgggctttgcccgggcggcctcagtgagcgagcgagcgcgcagctgcctgcaggggcgcctgatgcg
gtattttctccttacgcatctgtgcggtatttcacaccgcatacgtcaaagcaaccatagtacgcgccct
gtagcggcgcattaagcgcggcgggtgtggtggttacgcgcagcgtgaccgctacacttgccagcgccct
agcgcccgctcctttcgctttcttcccttcctttctcgccacgttcgccggctttccccgtcaagctcta
aatcgggggctccctttagggttccgatttagtgctttacggcacctcgaccccaaaaaacttgatttgg
gtgatggttcacgtagtgggccatcgccctgatagacggtttttcgccctttgacgttggagtccacgtt
ctttaatagtggactcttgttccaaactggaacaacactcaaccctatctcgggctattcttttgattta
taagggattttgccgatttcggcctattggttaaaaaatgagctgatttaacaaaaatttaacgcgaatt
ttaacaaaatattaacgtttacaattttatggtgcactctcagtacaatctgctctgatgccgcatagtt
aagccagccccgacacccgccaacacccgctgacgcgccctgacgggcttgtctgctcccggcatccgct
tacagacaagctgtgaccgtctccgggagctgcatgtgtcagaggttttcaccgtcatcaccgaaacgcg
cgagacgaaagggcctcgtgatacgcctatttttataggttaatgtcatgataataatggtttcttagac
gtcaggtggcacttttcggggaaatgtgcgcggaacccctatttgtttatttttctaaatacattcaaat
atgtatccgctcatgagacaataaccctgataaatgcttcaataatattgaaaaaggaagagtatgagta
ttcaacatttccgtgtcgcccttattcccttttttgcggcattttgccttcctgtttttgctcacccaga
aacgctggtgaaagtaaaagatgctgaagatcagttgggtgcacgagtgggttacatcgaactggatctc
aacagcggtaagatccttgagagttttcgccccgaagaacgttttccaatgatgagcacttttaaagttc
tgctatgtggcgcggtattatcccgtattgacgccgggcaagagcaactcggtcgccgcatacactattc
tcagaatgacttggttgagtactcaccagtcacagaaaagcatcttacggatggcatgacagtaagagaa
ttatgcagtgctgccataaccatgagtgataacactgcggccaacttacttctgacaacgatcggaggac
cgaaggagctaaccgcttttttgcacaacatgggggatcatgtaactcgccttgatcgttgggaaccgga
gctgaatgaagccataccaaacgacgagcgtgacaccacgatgcctgtagcaatggcaacaacgttgcgc
aaactattaactggcgaactacttactctagcttcccggcaacaattaatagactggatggaggcggata
aagttgcaggaccacttctgcgctcggcccttccggctggctggtttattgctgataaatctggagccgg
tgagcgtggaagccgcggtatcattgcagcactggggccagatggtaagccctcccgtatcgtagttatc
tacacgacggggagtcaggcaactatggatgaacgaaatagacagatcgctgagataggtgcctcactga
ttaagcattggtaactgtcagaccaagtttactcatatatactttagattgatttaaaacttcattttta
atttaaaaggatctaggtgaagatcctttttgataatctcatgaccaaaatcccttaacgtgagttttcg
ttccactgagcgtcagaccccgtagaaaagatcaaaggatcttcttgagatcctttttttctgcgcgtaa
tctgctgcttgcaaacaaaaaaaccaccgctaccagcggtggtttgtttgccggatcaagagctaccaac
tctttttccgaaggtaactggcttcagcagagcgcagataccaaatactgtccttctagtgtagccgtag
ttaggccaccacttcaagaactctgtagcaccgcctacatacctcgctctgctaatcctgttaccagtgg
ctgctgccagtggcgataagtcgtgtcttaccgggttggactcaagacgatagttaccggataaggcgca
gcggtcgggctgaacggggggttcgtgcacacagcccagcttggagcgaacgacctacaccgaactgaga
tacctacagcgtgagctatgagaaagcgccacgcttcccgaagggagaaaggcggacaggtatccggtaa
gcggcagggtcggaacaggagagcgcacgagggagcttccagggggaaacgcctggtatctttatagtcc

tgtcgggtttcgccacctctgacttgagcgtcgatttttgtgatgctcgtcaggggggcggagcctatgg
aaaaacgccagcaacgcggcctttttacggttcctggccttttgctggccttttgctcacatgt

**Supplementary Note 1**. FACS gating examples for GFP-positive or Cherry-positive cells.

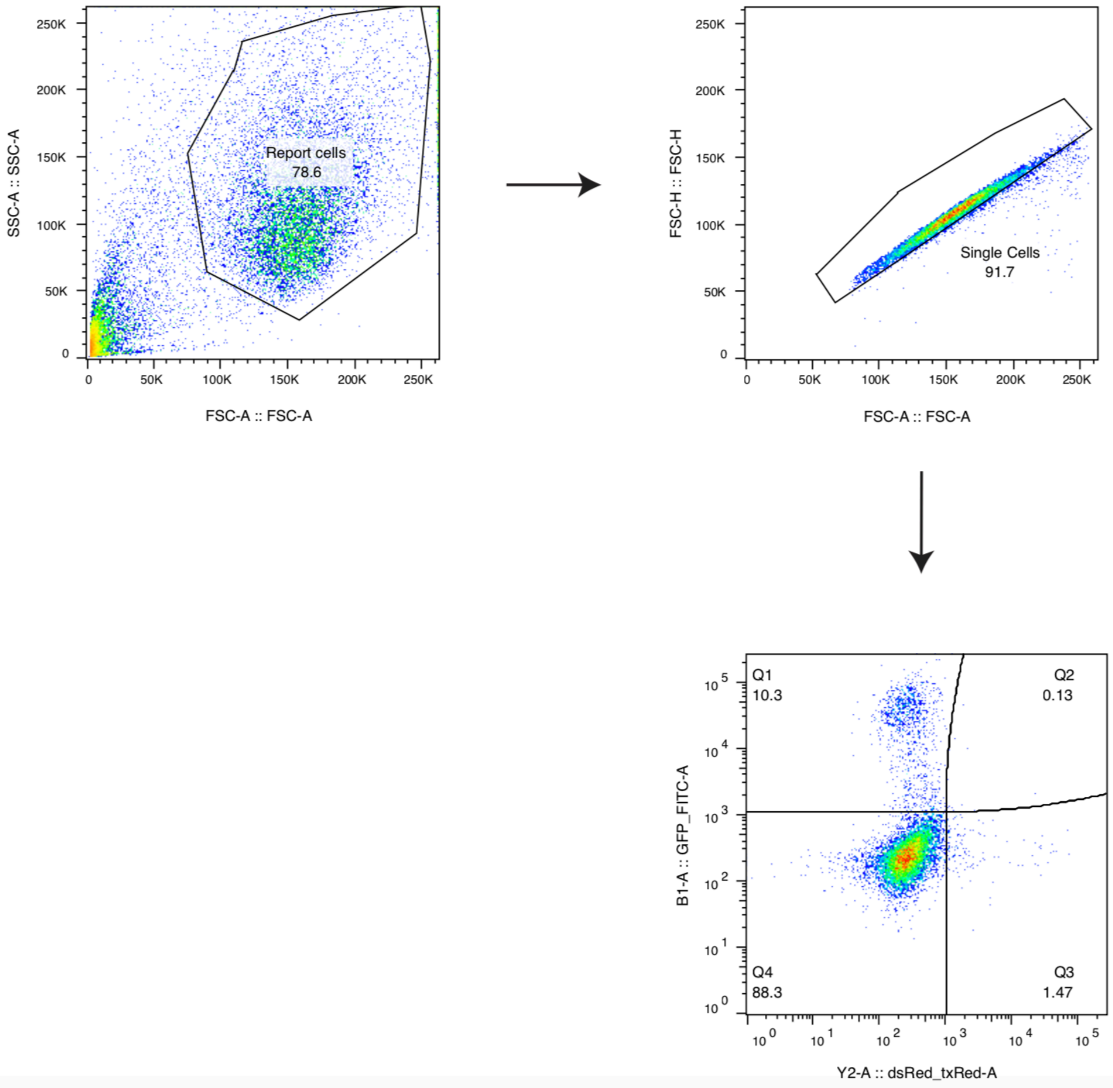
